# Reference genome choice and filtering thresholds jointly influence phylogenomic analyses

**DOI:** 10.1101/2022.03.10.483737

**Authors:** Jessica A. Rick, Chad D. Brock, Alexander L. Lewanski, Jimena Golcher-Benavides, Catherine E. Wagner

## Abstract

Molecular phylogenies are a cornerstone of modern comparative biology and are commonly employed to investigate a range of biological phenomena, such as diversification rates, patterns in trait evolution, biogeography, and community assembly. Recent work has demonstrated that significant biases may be introduced into downstream phylogenetic analyses from processing genomic data; however, it remains unclear whether there are interactions among bioinformatic parameters or biases introduced through the choice of reference genome for sequence alignment and variant-calling. We address these knowledge gaps by employing a combination of simulated and empirical data sets to investigate to what extent the choice of reference genome in upstream bioinformatic processing of genomic data influences phylogenetic inference, as well as the way that reference genome choice interacts with bioinformatic filtering choices and phylogenetic inference method. We demonstrate that more stringent minor allele filters bias inferred trees away from the true species tree topology, and that these biased trees tend to be more imbalanced and have a higher center of gravity than the true trees. We find greatest topological accuracy when filtering sites for minor allele count *>*3–4 in our 51-taxa data sets, while tree center of gravity was closest to the true value when filtering for sites with minor allele count *>*1–2. In contrast, filtering for missing data increased accuracy in the inferred topologies; however, this effect was small in comparison to the effect of minor allele filters and may be undesirable due to a subsequent mutation spectrum distortion. The bias introduced by these filters differs based on the reference genome used in short read alignment, providing further support that choosing a reference genome for alignment is an important bioinformatic decision with implications for downstream analyses. These results demonstrate that attributes of the study system and dataset (and their interaction) add important nuance for how best to assemble and filter short read genomic data for phylogenetic inference.

---

Phylogenetic history provides a critical framework for understanding the myriad components of a lineage’s evolution, and this “tree thinking” approach is a cornerstone of modern comparative biology (O’Hara, 1997; Pagel, 1999). Large multilocus data sets have become increasingly important for inferring phylogenetic histories (e.g., Edwards, 2009; Lemmon and Lemmon, 2013; Chan et al., 2020; Cloutier et al., 2019; Huang et al., 2020), and our ability to resolve relationships among rapidly diversifying taxa has greatly improved as a result of advances in high-throughput sequencing methods (e.g., Wagner et al., 2013; Rokas et al., 2003; Philippe et al., 2004; Prasad et al., 2008). However, inferring phylogenetic relationships is not a trivial task, even with genomic data. These large data sets also present novel—and in some cases, unexplored—challenges to phylogenetic analyses (e.g., Kumar et al., 2012; Jeffroy et al., 2006).

Phylogenetic inferences may be sensitive to both sampling and analytical approaches, and previous work has demonstrated that factors such as taxon sampling (Rannala et al., 1998; Zwickl and Hillis, 2002; Cusimano and Renner, 2010; Brock et al., 2011; Heath et al., 2008a,b), missing data (Wiens, 1998, 2003; Pybus and Harvey, 2000; Wiens and Moen, 2008; Wiens and Tiu, 2012; Grievink et al., 2013; Lemmon et al., 2009), and method of phylogenetic reconstruction (Huelsenbeck and Kirkpatrick, 1996; Revell et al., 2005; Rüber and Zardoya, 2005; Grievink et al., 2013; Leaché et al., 2015) can affect phylogenetic accuracy and downstream inferences about the tempo and mode of evolution. While bioinformatic filters—such as those for read depth, missing data per site, missing data per individual, and minor allele count or frequency—are often employed to guard against sequencing and alignment errors, these choices can have unintended consequences. Not only do these filters tend to reduce the number of loci in the data set, they can also bias the characteristics of those loci that are retained for downstream analyses. In population genetics, rare variants (i.e., those at *<*5% frequency in a population) are common and often of interest (Biddanda et al., 2020), particularly in disease association studies (e.g., Nelson et al., 2012; Momozawa and Mizukami, 2021; Weiner et al., 2023) and analyses of fine-scale population structure (e.g., Slatkin, 1985; Novembre et al., 2008), and the inclusion or removal of these low-frequency sites can affect estimates of gene flow (Slatkin, 1985) or differentiation among populations (Bhatia et al., 2013). Several recent studies have also demonstrated that discarding sites with low minor allele frequencies truncates the inferred allele frequency spectrum by removing the most recent mutations (Linck and Battey, 2019; Shafer et al., 2017), which can lead to biased estimates of historical demography and evolutionary processes, particularly in the recent past (Boitard et al., 2016).

Similarly, missing data in reduced-representation sequencing are expected to be non-randomly distributed across species and loci, with the amount of missing data proportional to the genetic distance between taxa (Eaton et al., 2017) and to locus-specific mutation rates. Filtering loci by amount of missing data thus disproportionately excludes loci with high mutation rates, which artificially truncates the mutational spectrum of the remaining loci (Huang and Knowles, 2016). This has important implications for analyses explicitly using the mutational spectrum for modeling evolutionary history (e.g., inferences of historical demography, Gutenkunst et al., 2009; Hotaling et al., 2018; Boitard et al., 2016; McTavish and Hillis, 2015; DeWitt et al., 2021), as well as for phylogenetic analyses where shared mutations provide information about evolutionary relationships and branch lengths. Loci with high mutation rates tend to be more phylogenetically informative, and therefore contribute more to accurate tree estimation (Lanier et al., 2014). Consequently, removing these loci may then lead to severe drops in tree estimation accuracy. Consistent with this, Molloy and Warnow (2018) demonstrated that filtering genes based on missing data generally reduces the accuracy of both species tree and concatenation-based phylogenetic analyses, with a greater effect in trees with more incomplete lineage sorting.

Recent advances in sequencing technologies along with falling sequencing costs have made it increasingly feasible to sequence and assemble reference genomes for non-model species (Feng et al., 2020; Kolora et al., 2021; Hotaling et al., 2021; Formenti et al., 2022; Rhie et al., 2021). Non-model reference genomes can then be used to assemble short reads from reduced-representation or whole genome sequencing into shared loci across individuals, and an increasing number of studies use a reference genome of a focal study taxon for assembly. Assembly of reads to a reference genome is based on sequence similarity (Catchen et al., 2011), and reads with higher mutation rates and higher variability will tend to have lower alignment scores (Nielsen et al., 2011; Huang and Knowles, 2016). Therefore, the choice of an ingroup (i.e., closely-related) versus an outgroup (i.e., more distant) reference genome with which to align reads may have substantial consequences for downstream phylogenetic analyses. For example, there are likely to be more reads with lower alignment scores when aligning to a more distant genome than when using an ingroup reference, and conserved regions with low mutation rates are more likely to be kept post-filtering (McCormack et al., 2009; Huang and Knowles, 2016). A consequence of this is that DNA fragments carrying the reference allele will be more likely to map successfully and pass quality filters, and therefore be retained in analyses (Ros-Freixedes et al., 2018; Brandt et al., 2015). This bias is expected to increase with increasing genetic distance to the reference genome and to be more prevalent in regions that are highly polymorphic. Following this prediction, several recent studies have demonstrated that reference genome choices can bias population genetic inferences, such as in estimates of historical demography, heterozygosity, and runs of homozygosity (Prasad et al., 2021; Günther and Nettelblad, 2019; Reid et al., 2021; Shafer et al., 2017). In phylogenetic analyses, the use of more divergent reference genomes has been shown to influence the accuracy of heterozygote calls (Duchen and Salamin, 2021). Additionally, in bacteria, the failure of more divergent regions of query sequences to map to the reference sequence can bias both topological accuracy and branch length estimation (Bertels et al., 2014). With this knowledge, a more general treatment of biases introduced by reference genome choices in phylogenetics is warranted.

Systematic biases in sites retained for analyses may hinder our ability to recover the true evolutionary history of a clade, and in doing so, may bias inferences made about the tempo and mode of evolution. Small changes in the sites retained can have large influences on the inferred topology and branch lengths (Shen et al., 2017) and therefore the diversification patterns inferred from that topology. For example, more stringent minor allele count or frequency filters may remove a greater proportion of data from branches at the tips of a phylogeny, leading to the truncation of these branches and an erroneous signal of increasing diversification in the recent past. Employing a more distant reference genome may have a similar effect, as this could lead to lower alignment scores for the more rapidly evolving sites most likely to change in recently diverged groups. Alternatively, as missing data is likely to increase with increasing genetic distance between taxa (e.g., due to loss of restriction cut sites; Huang and Knowles, 2016; Eaton et al., 2017), missing data filters may disproportionately affect branches toward the base of the tree. Stringent filtering may reduce data set size, as well as the distribution of signal across a phylogeny. As phylogenetically informative sites are lost, increasing noise could lead to more imbalanced topologies, on average, because there are more possibilities for a tree to be unbalanced than balanced (Heard and Mooers, 1996; Huelsenbeck and Kirkpatrick, 1996; Alanzi and Degnan, 2017; Alanzi, 2020).

Unfortunately, there is a lack of consensus on how best to process data for phylogenetic analysis. For example, some studies recommend filtering data stringently to ensure sequence accuracy (Davey et al., 2013). Others demonstrate that filtering can bias important features of the resulting data (Nielsen, 2000; Mastretta-Yanes et al., 2015; Huang and Knowles, 2016; Chan et al., 2020), and therefore recommend against filtering stringently. It may also be the case that there is not a universally “best” approach to these decisions (Nazareno and Knowles, 2021); variation in empirical data attributes such as divergence times, incomplete lineage sorting, and genetic diversity may mean that different filtering decisions are optimal in different scenarios. For example, Huang and Knowles (2016) show that high missing data filters are particularly harmful for data sets with short coalescent times and it is likely that the consequences of other filters are similarly context dependent. Consequently, there may not be a one-size-fits-all prescription that can be made for the treatment of data for phylogenetics and the proper filtering choices may be contingent on the focus of the analysis.

Here, we assess the influence of bioinformatic choices on phylogenetic inference by combining simulations and empirical data sets to assess the interacting influences of reference genome choice, minor allele filters, and missing data filters on inferred phylogenies. We assess our ability to recover both the true tree and the true summary statistics describing tree shape and imbalance, and address three main questions: (1) To what extent do bioinformatic choices affect our ability to recover the ‘true’ phylogenetic relationships? (2) How do bioinformatic choices bias inferences of diversification patterns based on tree shape and node distribution? and (3) How do these biases depend on the diversification history within the clade? Our results demonstrate that decisions made by researchers during the assembly and processing of genomic data, as well as characteristics of the true divergence history among taxa, can bias the data set retained and the inferred phylogenetic tree in ways that are consequential for macroevolutionary analyses. Finally, while our inferences focus on single nucleotide polymorphism (SNP) data sets derived from reduced-representation data, we outline how these findings are also applicable to the whole genome data sets that are becoming more common in modern phylogenomics.

## Materials and Methods

### Overview

We inferred phylogenies for simulated data sets aligned to reference genomes at varying genetic distances to the ingroup taxa. We quantified differences between the true species tree and trees inferred from 210 different combinations of alignment and filtering options for 10 simulated data sets (Fig. 1), two empirical data sets, and for two different choices of reference genome in each data set. We quantified topological differences between phylogenies for all simulated and empirical trees using three different tree distance metrics. To summarize multi-dimensional tree shapes and branching patterns, we also calculate summary statistics for tree shape, imbalance, and overall bootstrap support. For describing tree shape and branching patterns, we focus on three metrics: Colless’ imbalance statistic (*I_c_*; Colless, 1982), the Sackin imbalance statistic (Sackin, 1972), and the *γ* statistic (Pybus and Harvey, 2000). The Colless and Sackin statistics both increase with increasing tree imbalance (i.e., more pectinate topologies). The *γ* statistic is a measure of the distribution of branching events within a phylogeny; a constant-rate pure-birth tree has an expected *γ* = 0, and an increase in *γ* corresponds to branching times in a tree being closer to the tips (i.e., a less bottom-heavy topology with higher center of gravity), while a negative *γ* estimate indicates a tree with more diversification events toward the root (i.e., a more bottom-heavy topology).

**Fig. 1.**
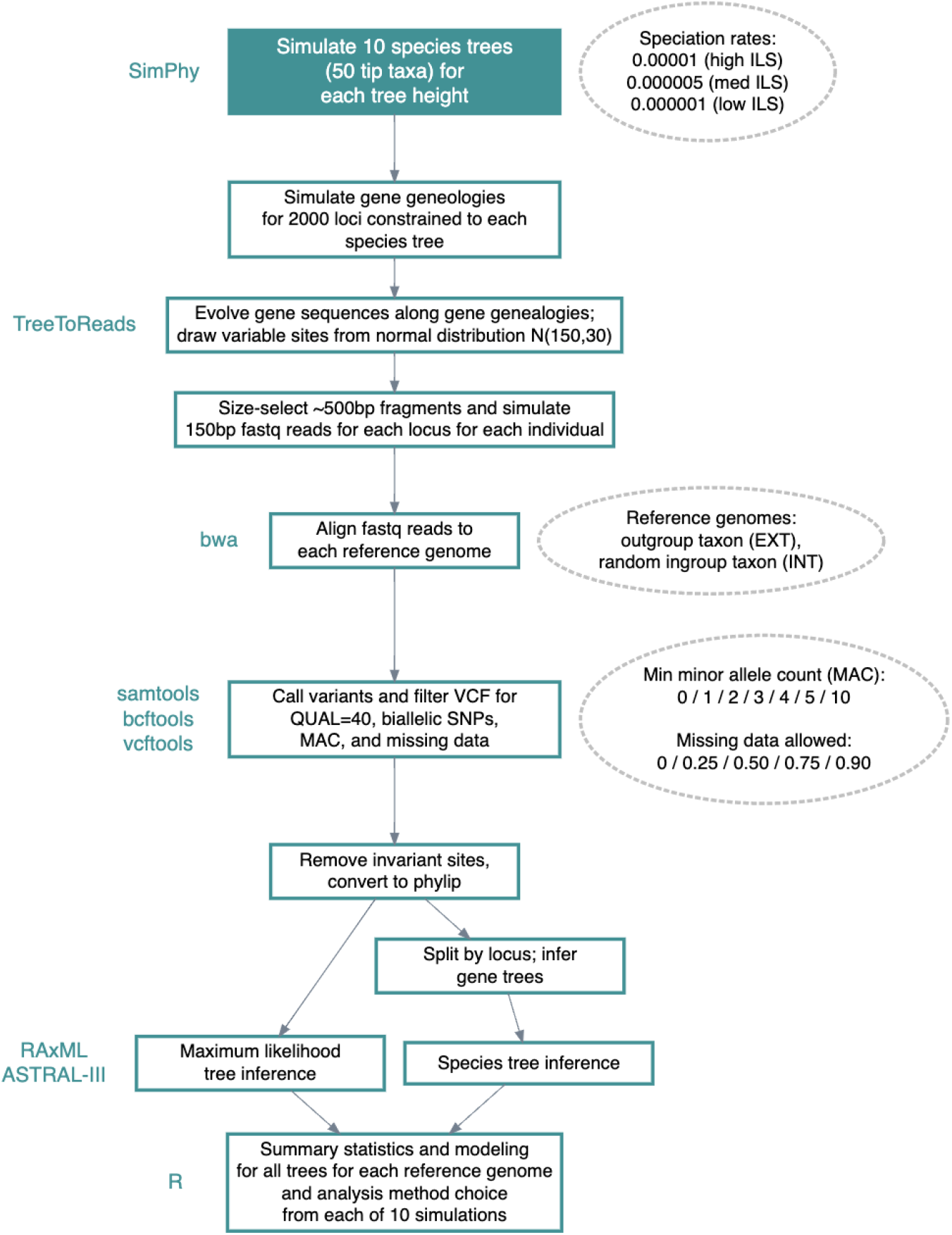
We used simulated data and varying bioinformatic filters to compare inferred phylogenies to the true topology used to generate data. This schematic shows parameter choices that we iterated over (rounded shapes) and steps performed, from species tree simulation to data analysis (square shapes). Each round of simulation resulted in 210 filtered phylogenetic trees for analysis from each combination of reference genome choice and phylogenetic inference method, and the simulation was replicated for 10 different simulated species trees at each speciation rate. *QUAL = mapping quality; ILS = incomplete lineage sorting; VCF = variant call format*.

### Simulated Data

To replicate the various properties of reduced-representation genotyping-by-sequencing (GBS) or restriction-associated-digest sequencing (RADseq), we simulated 60 independent data sets using the following procedure (Fig. 1): (1) simulate a species tree for 50 individuals plus one outgroup taxon at 3 different levels of incomplete lineage sorting; (2) simulate 2,000 gene genealogies for 5,000bp loci under the multi-species coalescent conditioned on the simulated species tree from (1); (3) simulate genome sequences and 150bp fastq reads for each individual for each gene genealogy from (2); (4) align simulated fastq reads to one ingroup and one outgroup reference genome; (5) call variants separately on ingroup and outgroup reference data sets; (6) filter variants with different parameter combinations; (7) infer a phylogeny either via concatenation and maximum likelihood or by first estimating individual gene trees that are then used to estimate the species tree; and finally, (8) calculate summary statistics on the output trees and model the contributions of each bioinformatic filter to differences in these statistics. Each of these steps is explained in detail below.

#### Species and gene trees

We simulated 50-taxon species trees (with one individual per taxon) using `SimPhy` (Mallo et al., 2016) and employing the Yule process with three different birth rates (-SB 0.00001, 0.000005, 0.000001) to represent three different levels of shared polymorphism due to incomplete lineage sorting (ILS; high, medium, and low, respectively, sensu Mirarab and Warnow 2015). All simulations had a tree-wide effective population size of 50,000 (`-SP f:50000`) and a ratio of 10 between the ingroup height and the branch from the root to the ingroup (`-SO f:10`). We then simulated genealogies for 2,000 loci of 5,000bp each, conditioned on the species tree for the given simulation and using a tree-wide substitution rate of 3 *×* 10*^−^*^9^ per generation (similar to that estimated in Lake Malawi cichlid fishes; Malinsky et al., 2018) for converting branch lengths to the expected number of substitutions per site. The ancestral gene sequences for these loci were pulled randomly without replacement from the empirical *Lates calcarifer* reference genome (Vij et al., 2016) (see Supplemental Methods). With these simulation parameters, ILS levels and gene tree discordance varied from a mean normalized Robinson-Foulds (RF) distance among gene trees of 0.061 to 0.845 (low ILS, mean = 0.165; medium ILS, mean = 0.568; high ILS, mean = 0.761).

#### Generating fastqs

We simulated fastq reads at each locus for each individual using the `TreeToReads` pipeline (McTavish et al., 2017). `TreeToReads` uses `Seq-Gen` (v1.3.4; Rambaut and Grass, 1997) for simulating variable sites and `ART` (vGreatSmokyMountains; Huang et al., 2012) to simulate Illumina reads. The pipeline generates simulated Illumina short-read data for individuals along an input gene genealogy, using a given number of variable sites, model of evolution, read length, and fragment length (McTavish et al., 2017). We simulated 500bp (s.d. 50) fragment lengths, to reflect size selection for *∼*500bp fragments in generating reduced-representation data. Reads were then simulated as 150bp single-end reads using the Illumina HiSeq2500 profile in `ART`. We pulled the number of variable sites for each fragment from a normal distribution with mean of 150 and standard deviation of 30. The location of mutations in the genome were drawn from a uniform distribution and generated using a generalized time reversible (GTR) model of sequence evolution with gamma rate heterogeneity (Lanave et al., 1984; Yang, 1993). While the substitution rate specified in `SeqGen` is used to translate branch lengths to number of substitutions for the input gene trees, the number of variable sites specified in `TreeToReads` translates those numbers of substitutions into mutations observed at in each locus, allowing for stochasticity in the location of mutations, and processes like “multiple hits”– mutations occurring at the same site. The read depth for each individual at each locus was drawn from a uniform distribution between 2 and 20, to generate heterogeneity in read depth and introduce missing data. We specified a sequencing error file parameterized using the empirical GBS data in Rick et al. (2022), to mimic empirical errors introduced in the sequencing process.

#### Aligning fastqs and calling variants

For each simulated tree, we randomly chose an ingroup taxon to be the ingroup reference, and used the simulated outgroup as our outgroup reference. We aligned all of the simulated fastq reads to each reference genome using `bwa mem` (v0.7.17, Li and Durbin 2009) and `samtools mpileup` (v1.6, Li 2011) with default parameters. We then filtered to retain only variable sites and called variants using `bcftools` (v1.9, Li et al. 2009). We retained variants in the raw VCF file that were biallelic (`--min-alleles 2` and `--max-alleles 2`), had a genotype quality score (`--minGQ`) *>*10, and had a minimum mapping quality (`--minQ`) *>*40.

We then further filtered variant sites using minor allele count (MAC) and missing data (miss) filters, based on parameter ranges observed in published studies. We chose seven different minimum minor allele count cutoffs (0, 1, 2, 3, 4, 5, 10) and five different missing data cutoffs (0, 0.25, 0.5, 0.75, 0.90; corresponding to the percent of genotyped taxa required for a site to be retained) in `vcftools` (v0.1.14, Danecek et al. 2011), and created SNP data sets using all combinations of these filtering parameters. Individual genotypes were only called with a read depth ⩾5 (`--minDP 5`). Following filtering, the resulting VCFs were converted to phylip format and invariant sites were removed using the `raxml ascbias.py` script v1.0 from https://github.com/btmartin721/raxml_ascbias. As a quantitative measure of divergence between the reference and ingroup lineages, we calculated for each unfiltered data set (MAC = 0, miss = 0) the mean pairwise genome-wide divergence, *d_xy_* (or *π_xy_*; Nei and Li, 1979) between the reference genome and all ingroup individuals using the `parseVCF.py` and `distMat.py` scripts from https://github.com/simonhmartin/genomics_general. Pairwise *d_xy_* is an absolute measure of divergence and reflects the proportion of shared sites that differ between two sequences (Cruickshank and Hahn, 2014).

#### Tree estimation

We used two different methods for tree estimation: a concatenation-maximum likelihood method (i.e., `RAxML`) and a summary species tree method (i.e., `ASTRAL-III`). Concatenation analyses ignore variance in patterns of coalescence across the genome due to incomplete lineage sorting (ILS), while species tree methods more explicitly account for ILS and gene tree discordance resulting from ILS (Liu et al., 2015). Because we simulated trees with varying amounts of ILS, we would expect a species tree method to perform better at recovering trees with high ILS due to explicitly accounting for ILS in its inference and thus it is of interest to determine whether concatenation and species tree methods differ in their sensitivity to our simulation parameters and filtering criteria.

For concatenation analyses, we concatenated variant sites and inferred phylogenies for each of the 4,200 simulation and filtering combinations under a GTRCAT model of rate heterogeneity with Lewis-type ascertainment bias correction (Leaché et al., 2015) in a full search for the best scoring ML tree with rapid bootstrap analysis (100 bootstraps) in the hybrid MPI/Pthreads version of `RAxML` (v8.2.12, Stamatakis, 2014). For species tree analyses, we constructed individual gene trees using `RAxML` with a GTRCAT model of rate heterogeneity and Lewis-type ascertainment bias correction. We then used `ASTRAL-III` (Zhang et al., 2018) to estimate the unrooted species tree given this set of gene trees. `ASTRAL` is statistically consistent under the multi-species coalescent model and finds the species tree that has the maximum number of shared quartet trees with the set of gene trees.

To measure the effect that each of the bioinformatic choices and tree construction methods has on downstream phylogenetic analyses, we calculated the information-corrected normalized RF distance (`InfoRobinsonFoulds` from the `TreeDist` package in `R`; Smith, 2020) between each inferred tree and the corresponding species tree (the “true” tree). Due to the inability of RF distance to capture all aspects of tree differences (Smith, 2020), we used two additional metrics of distance to the true tree: quartet distance (Estabrook et al., 1985) and clustering information distance (Smith, 2020). Quartet distance was calculated using `QuartetDivergence` using the tqDist algorithm (Sand et al., 2014) from the `Quartet` package in `R` (Smith, 2019), and is a count of the number of four-taxon statements (quartets) that differ between two trees. Clustering information distance scores a pair of trees based on the amount of information that one partition contains about the other and is an information-based tree distance metric that recognizes similarity in tree structure even when every possible pairing of splits conflicts (Smith, 2020), making it more sensitive than RF distance when trees are very dissimilar from one another. We also calculated two measures of imbalance, the Sackin statistic (Sackin, 1972) and Colless’ imbalance statistic (*I_c_*, Colless, 1982), as well as the tree center of gravity via the *γ* statistic (`gammaStat` from `ape`; Pybus and Harvey, 2000) and branch length distributions. These statistics are often used in diversification rate analyses, and therefore any systematic bias in their direction or magnitude may have important consequences for evolutionary inferences made in these types of analyses. The Sackin and Colless indices were standardized using the PDA model (Blum et al., 2006). We additionally calculated imbalance and *γ* statistics using only the ingroup taxa and standardized these statistics by subtracting the value of each statistic for the true tree from the value calculated for the inferred tree, thus making those values the deviation from the true statistic (sensu Stadler et al., 2016). Because trees inferred in `ASTRAL` have no terminal branch lengths and non-terminal coalescent unit branch lengths that are not robust to gene tree estimation error (Sayyari and Mirarab, 2016), we calculated only those summary statistics that are not dependent on branch length (i.e., omitted *γ* and tree height calculations) for these sets of trees.

### Statistical Analyses

All statistical analyses were conducted in R (v4.0.4, R Core Team, 2021) and plotted using `ggplot2` (Wickham, 2016). Scripts associated with simulations and analyses, as well as simulation outputs, are available at https://github.com/jessicarick/refbias_scripts.

#### Biases in lost loci

We first examined the effect of bioinformatic filters on the number of loci (size of the data matrix) and the corresponding mutational spectrum of retained SNPs. We quantified locus-specific mutation rate as the number of simulated variable sites for a given 5,000bp locus. We then compared mutation rates for SNPs pre- and post-filtering to investigate whether there were biases in the mutation rates of SNPs lost via different filters. We used pairwise Kolmogorov-Smirnov tests (`ks.test` in `R`) to statistically compare distributions of mutation rates from lost SNPs at each MAC and missing data cutoff. We also calculated the distribution of mutation rates of loci lost with increasing MAC and missing data filters to determine whether loci with higher mutation rates are more likely to be lost first, as has recently been demonstrated in other studies (e.g., Linck and Battey 2019; Huang and Knowles 2016; Shafer et al. 2017).

To investigate whether there is a pattern to signal loss in the phylogeny due to filtering, we compared the mean number of descendants and bootstrap support for nodes retained in each filtering step for the `RAxML`-inferred trees. If signal is disproportionally lost at the tips of the tree due to the MAC or missing data filters, then we would expect the nodes with fewer descendants to disproportionally be lost in the inferred trees. We used the `phytools` package (Revell, 2012) to compare each output tree to the original simulated tree (using the `matchNodes` function), which provides us with information about which nodes from the original tree are missing in the output tree. We then recorded the bootstrap support (from `RAxML` output) and number of node descendants (using `getDescendants` from `phytools`) for each node in the original tree. For each combination of ILS level, MAC, missing data, and reference genome choice, we created a null distribution for the expected mean node descendants and expected mean nodal bootstrap support using 100 bootstrap replicates, by taking the mean of *n* values drawn from the possible values for each of these, where *n* is the number of nodes lost across all trees with the given parameters choices. From these values, we calculated a z-score for the observed mean to quantify deviation from the bootstrapped mean value. We repeated the bootstrapping procedure for the nodes retained in each parameter combination, and calculated z-scores for these as well.

#### Tree space analyses

To further characterize dissimilarity among the inferred phylogenetic trees within each simulation, we calculated the information-corrected normalized (Smith, 2020) RF distances (Robinson and Foulds, 1981) between all pairwise comparisons of the output trees that had the same starting species tree (`InfoRobinsonFoulds` from `TreeDist`; Smith, 2020). As noted above, we also compared each output tree to their starting (“true”) species tree and then performed a principal coordinates analysis (PCoA) on all of these pairwise distances across the starting trees (using `pcoa` from `ape`, Paradis and Schliep 2019). Using a PCoA approach, we can visualize differences between trees in multidimensional “tree space” and determine the differentiation among trees with different parameter combinations. These results complement our direct comparisons to the “true” species tree described above by allowing us to additionally assess the level of variation across output trees under each bioinformatic scenario. We calculated the correlation between tree position on the first two PCoA axes and each of our filtering parameters and reference genome choice for each group of trees to assess the extent to which each of these parameters drives the clustering of the inferred trees in tree space.

#### Linear modeling

To characterize the relationship between parameter choices (predictors) and the resulting trees and tree statistics (responses), we used linear mixed models (implemented using `lme4`; Bates et al., 2015) in `R`. In our linear mixed models, for each tree height, our predictor variables included reference distance (mean *d_xy_* between the ingroup taxa and the reference taxon), minor allele count cutoff (MAC), and missing data cutoff, as well as pairwise interaction terms between reference distance and each of MAC and missing data. We additionally included a random intercept for simulation number, such that the overall model was *X ∼ d_xy_* + *MAC* + *miss* + *d_xy_*: *MAC* + *d_xy_*: *miss* + (1*|sim*), where *X* is our tree statistic of interest, *d_xy_* is mean distance to the reference genome, *MAC* is minor allele count, *miss* is missing data threshold, and *sim* is simulation replicate number. We assessed model fit using conditional pseudo-*R*^2^ values (using `r.squaredGLMM` from `MuMIn`; Bartoń 2022) and assessed the contribution of each predictor variable to the response statistic using regression coefficients. We additionally assessed bivariate correlations between all variable pairs using Pearson’s correlation coefficient, implemented using cor.test in R.

### Accounting for the influence of data set size

Because each filtering step removes SNPs, and therefore decreases the amount of input data for phylogenetic analyses (Fig. S2), we performed further steps to parse out whether the observed effects were simply due to the loss of power in smaller data sets, or if additional effects could be attributed to the specific filters applied and a bias as to which sites are retained. To investigate the relative influence of each filter on the three response variables (RF distance to true tree, imbalance, and gamma), as well as the mutation spectrum, we subsampled each filtered data set to retain only 6,000 random SNPs. We then ran phylogenetic analyses in `RAxML` as described above. With these subsampled trees, we performed the same statistical and modeling steps described above for the full data sets, and compared these results to those with the full data sets.

### Empirical data

To investigate to what extent the same patterns observed in our simulated data may affect empirical data sets, we used two sets of GBS data from clades with similar ages, but different number of species and expected levels of gene tree discordance: the radiation of *Lates* fishes endemic to Lake Tanganyika and the radiation of cichlid fishes in the tribe Tropheini, also endemic to Lake Tanganyika. Both of these clades have an estimated origin around *∼*1–2 Ma (Koblmüller et al., 2021, 2010; Irisarri et al., 2018, although Ronco et al. 2021 estimates the origin of the tropheine radiation to be *∼*3.5–5.2 Ma), but very different species richness (n=4 *Lates spp.*; n*≈*40 described and undescribed tropheine spp., as estimated in Ronco et al. 2020). Thus, the tropheines represent a clade with high speciation rates, shorter internodes, and thus potentially higher amounts of ILS, while ILS is expected to be lower in the more slowly diversifying *Lates* clade. Sequences for the *Lates spp.* were generated as described in Rick et al. (2022) (NCBI Sequence Read Archive PRJNA776855) and data for the tropheines were generated in a similar manner following the GBS protocol in Parchman et al. (2012), as described in the Supplementary Methods.

Using the same methods as for the simulated data, the *Lates* fastq reads were aligned to the *L. calcarifer* (distant outgroup) and *L. mariae* (ingroup) reference genomes. The tropheine fastq reads were aligned to the *Oreochromis niloticus* (distant outgroup) and *Pundamilia nyererei* (closely related outgroup) reference genomes (Brawand et al., 2015). We randomly chose 49 individuals for each of 10 data sets for each clade out of the n = 229 high-quality (i.e., *>*100,000 reads assembled to each reference genome) sequenced individuals for the *Lates spp.* and n = 463 high-quality sequenced individuals for the tropheines. We then added the alignments for the two reference genome individuals to these data sets, so that the total number of tip taxa was n = 51, matching our simulations. Following alignment of fastq reads to the reference genomes, we iterated through missing data (miss = 0, 0.25, 0.50, 0.75, 0.90) and minor allele (MAC = 0, 1, 2, 3, 4, 5, 10) filters and inferred the phylogeny for each filtered data set using `RAxML` and `ASTRAL-III`, using the same procedures and parameter choices as with the simulated data. For the two empirical data sets, we retained only scaffolds greater than 100kb and used 10kb windows for gene tree inference in `RAxML` prior to `ASTRAL` inference. We then calculated the RF distance between all trees for each data set and inference method and visualized differences among the trees using PCoA in R. We calculated (for both `RAxML` and `ASTRAL` trees) and *γ* and tree height (for `RAxML trees`), and again used linear mixed models to assess the influence that each of our reference genome choices and bioinformatic filters had on these tree characteristics. With the empirical results, we treated the identity of the reference genome as a binary option (i.e., ingroup/close versus outgroup/distant reference), rather than as a continuous *d_xy_* variable, such that our models were *X ∼ ref* + *MAC* + *miss* + *ref*: *MAC* + *ref*: *miss* + (1*|sim*), where *ref* indicates reference genome choice and the ingroup/close reference was coded as 1 and the outgroup/distant reference was coded as 0.

## Results

### Simulations

Our simulation scheme resulted in a total of 4,200 phylogenetic trees across 10 replicate simulations and two phylogenetic inference methods, which we analyzed to determine the influence of reference genome and filtering choices on topology and tree shape metrics. Following filtering and removal of invariant sites, our simulated data sets retained an average of 77,549 SNPs (95% quantile = 8,320–177,830). The mean outgroup reference genome distance (*d_xy_*) was 0.40 (95% quantile = 0.30–0.53), while the ingroup reference genomes were an average distance of *d_xy_* = 0.039 (95% quantile = 0.031–0.047 from the other ingroup taxa. As was expected based on simulation parameters, the mean distance to the outgroup reference was greatest in the low-ILS trees (mean=0.45, range 0.39–0.53), lower in the medium-ILS trees (mean=0.38, range 0.33–0.46), and lowest in the high-ILS trees (mean=0.37, range 0.30–0.46), although all of these distributions overlapped (Fig. S7). The mean distance to the ingroup reference was similar among all of the tree heights (low-ILS trees: mean=0.040, range 0.031–0.046; medium-ILS trees: mean=0.039, range 0.035–0.047; high-ILS trees: mean=0.040, range=0.035–0.044; Fig. S7). Consistent with expectations, the average amount of gene tree discordance (i.e., mean pairwise normalized Robinson-Foulds distance between all gene trees) in the low-ILS trees was 0.164 (range=0.096–0.241), in medium-ILS trees was 0.606 (range=0.480–0.746), and in high-ILS trees was 0.771 (range=0.661–0.845). Our simulated species trees had overlapping distributions of normalized ingroup imbalance and gamma among tree heights (Fig. 2), with a mean ingroup *I_c_* of 0.412 and mean ingroup *γ* of -0.159. Example filtered phylogenies can be found in Fig. S1.

**Fig. 2.**
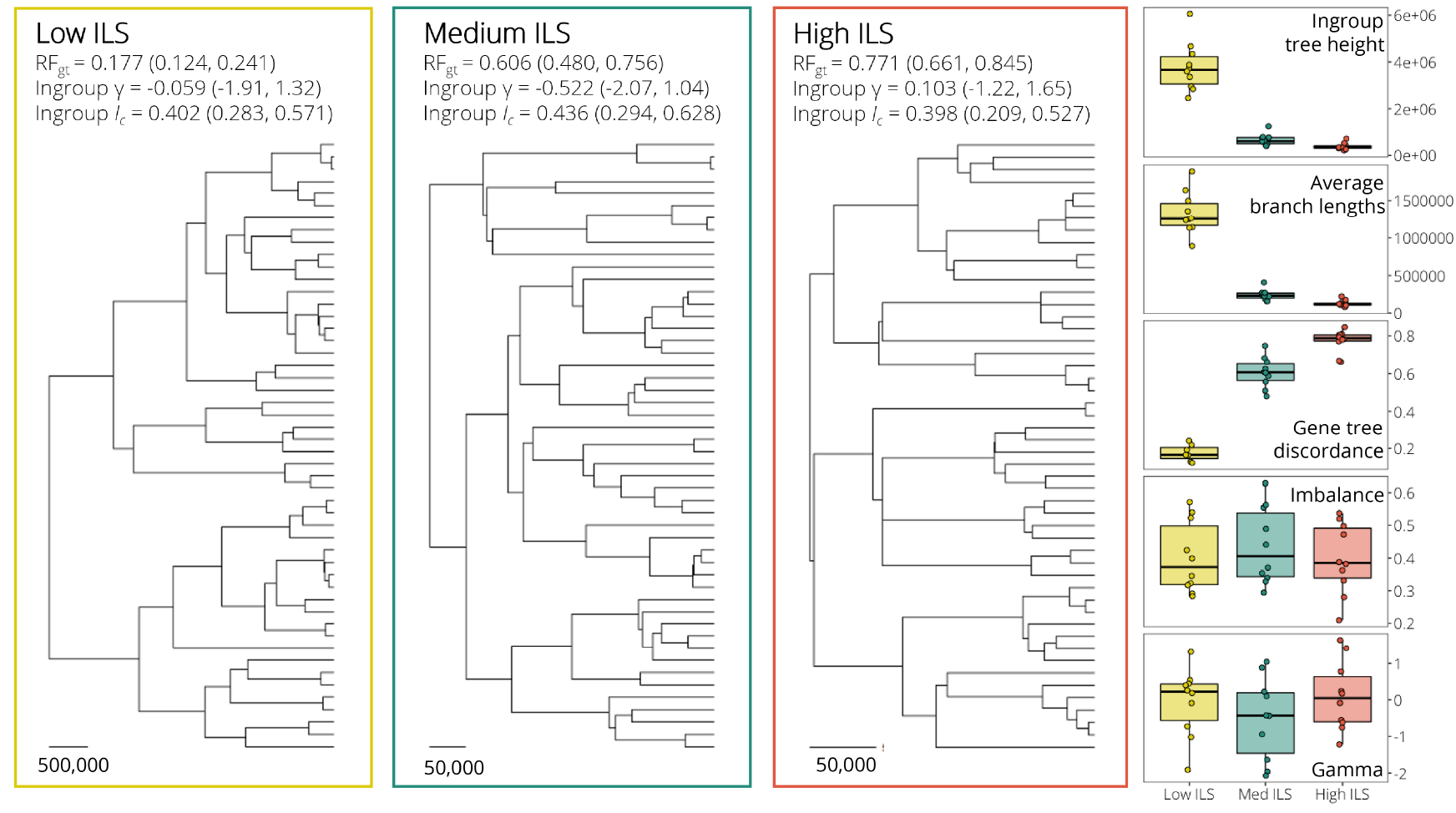
Example simulated species trees from `SimPhy` for low ILS (speciation rate = 0.000001; yellow), medium ILS (speciation rate = 0.000005; green), and high ILS (speciation rate = 0.00001; red) trees (sensu Mirarab and Warnow, 2015), and summary statistics for all simulated species trees. All trees were simulated with 50 taxa and an outgroup (not shown in phylogenies), on which the ingroup clade was rooted. For each tree, the mean across 10 simulated species trees for the gene tree discordance (as measured by mean pairwise Robinson-Foulds distance between gene trees; *RF_gt_*), mean ingroup gamma (*γ*), and mean Colless imbalance statistic (*I_c_*), along with their ranges, are given. Note that scales (in coalescent units) are not consistent among the trees shown. In the righthand panel, the distribution of values across all 30 simulated “true trees” is shown for ingroup tree height, average branch lengths, mean gene tree discordance, ingroup Colless imbalance, and ingroup gamma.

Correlations among all variables for the filtered data sets and `RAxML`-inferred trees are visualized in Figure S3a. Among all of our filtered data sets, minor allele count (MAC) threshold and number of SNPs retained were highly negatively correlated (Pearson’s *r*(2098) = *−*0.734*, p <* 2.2*e −* 16; Fig. S2,S3a). Our two measures of imbalance (i.e., Colless imbalance and Sackin imbalance) were highly correlated as well (Pearson’s *r*(2098) = 0.988*, p <* 2.2*e −* 16), and these were positively correlated with metrics of topological distance from the inferred tree to the true tree (i.e., Robinson-Foulds distance, quartet distance, and clustering information distance; mean Pearson’s *r*(2098) = 0.252, all *p <* 0.001), although quartet distance was less strongly correlated with imbalance than the other topological distance metrics. Ingroup *γ* was inversely correlated to tree height (Pearson’s *r*(2098) = 0.781, *p <* 2.2*e −* 16)—as is expected given our simulation methods—and slightly positively correlated to imbalance and topological distance metrics (Colless imbalance: Pearson’s *r*(2098) = 0.319, *p <* 2.2*e −* 16; RF distance: Pearson’s *r*(2098) = 0.329, *p <* 2.2*e −* 16). Topological distance from the inferred tree to the true tree was negatively correlated with average bootstrap support (Pearson’s *r*(2098) = 0.779, *p <* 2.2*e −* 16). Heatmaps visualizing the way in which each of the inferred tree characteristics varied across MAC and missing data thresholds for the `RAxML` trees can be found in Fig. S8–S15. Correlations among statistics were similar for the `ASTRAL`-inferred phylogenies (Fig. S3b), with the direction of the relatinoship between terms consistent between the two inference methods for 52 of the 72 correlations. Furthermore, the adjusted RV coefficient (Mayer et al., 2011) between the `RAxML` and `ASTRAL` correlation matrices (Fig. S3) was 0.72, indicating substantial similarity among matrices.

#### Distance to true tree

In our analyses, the ability to recover the input species tree topology in simulated data sets varied widely across filtering parameters and reference genome distance. All four of our linear mixed models for Robinson-Foulds distance between the `RAxML` trees and their true trees explained a majority of the variation in the data (mean conditional pseudo-*R*^2^ = 0.669, range 0.558–0.758; Table S1), and minor allele count had the greatest effect on the distance of the inferred phylogeny to the true tree. MAC was an important predictor variable in our linear models for distance to the true tree across all three simulated tree heights, with increased MAC corresponding to topologies more distant to the true tree (Fig. 3). Missing data had less of an effect on topological accuracy, and was not an important predictor variable in our models for all trees combined, or for each of the three tree heights individually (Fig. 3; Table S1). While we only used RF distance in our linear mixed models, we observed similar trends across MAC in quartet and clustering information distances, such that all three topological distance statistics increased with increasing minor allele count threshold (Fig. S22).

**Fig. 3.**
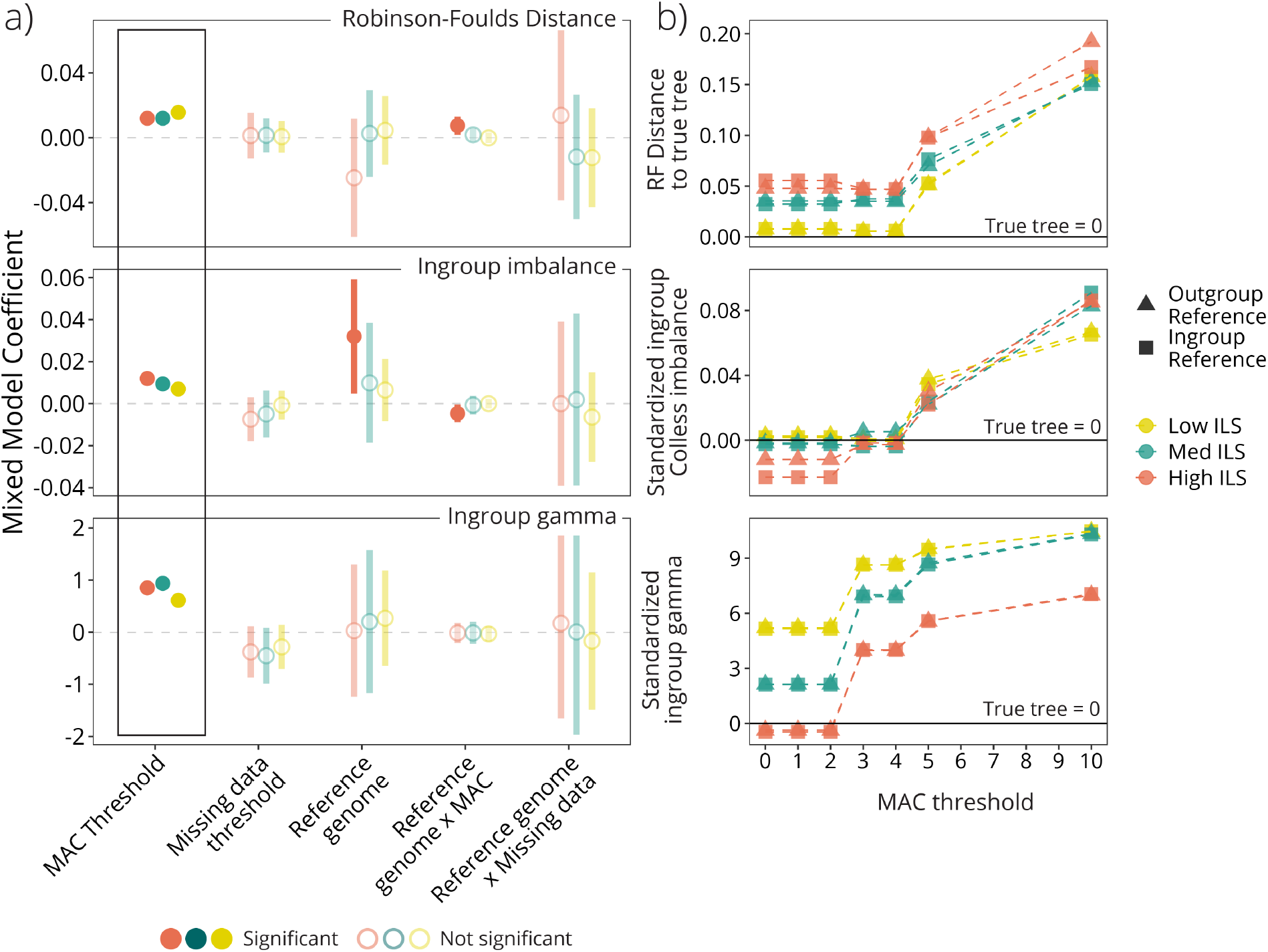
Minor allele count (MAC) threshold had the greatest effect on the calculated tree statistics for our simulated data. MAC threshold was a significant predictor of (1) distance from the inferred tree to the true tree, (2) the imbalance, and (3) the *γ* statistic of the output tree. MAC threshold also interacted with the distance to the reference genome and tree height. Plots in (a) show coefficient estimates and 95% confidence intervals for each predictor variable from linear mixed models for the `RAxML` trees using the indicated statistic as the response variable (Robinson-Foulds (RF) distance to the true tree, the gamma statistic for tree shape (*γ*), and Colless’ imbalance statistic (*I_c_*)). Filled in and darkened symbols indicate significant positive or negative relationships, while faded open symbols are predictors with confidence intervals that overlap zero. Plots in (b) show patterns of variation in the three response variables for the ingroup across minor allele counts, with shapes indicating the reference genome used (EXT, external reference; INT, internal reference) and colors indicating the level of ILS. In (b), the statistics are standardized so that the true tree is 0, as indicated with the solid horizontal lines.

Similar to the modeling results for `RAxML` trees, all four of the linear mixed models for RF distance between the `ASTRAL` trees and their true trees explained a majority of the variation in the data (mean conditional pseudo-*R*^2^ = 0.742, range 0.636–0.813; Table S4). Minor allele count threshold again had the greatest effect on the distance of the inferred phylogenies to the true trees (with more stringent MAC thresholds leading to trees more different from the true topology), while missing data also was a significant predictor for the low ILS trees, high ILS trees, and models with all of the trees combined (Fig. S5b,d).

The influence of reference genome choice had the greatest effect in high-ILS trees and the weakest effect in low-ILS trees for the `RAxML` trees (Fig. 3a). Across all three topological distance metrics, trees constructed with the ingroup reference genome were generally similar distances from the true tree as those using the outgroup reference genome, with some variation among MAC thresholds and tree height (Fig. S22a,c,e). High ILS trees aligned to the ingroup reference and filtered using low MAC thresholds were more distant to the true tree than outgroup-aligned reference trees, but at high MAC thresholds the ingroup reference trees are more similar to the true tree (i.e., have higher topological accuracy). Robinson-Foulds and clustering information distances identified the greatest difference between trees inferred using ingroup versus outgroup reference genomes at high MAC thresholds. Quartet distance metrics (Fig. S22c) indicate that at low MAC thresholds for medium and high ILS trees, trees constructed from data aligned to an outgroup reference genome were generally more similar to the true tree than those aligned to an ingroup reference genome. In contrast, at high MAC thresholds, ingroup reference genomes produced topologies closer to the true species tree for medium and high ILS trees. Trees with low ILS had a smaller quartet distance difference between ingroup and outgroup reference genomes, and ingroup reference genomes produced trees more similar to the true tree across all MAC thresholds for these low ILS trees (Fig. S22c). While the `ASTRAL`-inferred trees generally showed the same patterns, the low ILS trees with high MAC thresholds (i.e., most stringent filtering) were much more distant from the true tree than those inferred using `RAxML`. Across all MAC thresholds, the low ILS trees inferred using `ASTRAL` were more distant from the true tree than those inferred using `RAxML` (Fig. S26a,b).

Using PCoA to visualize tree space for `RAxML`-inferred trees (Fig. 4), we find that the first two PCoA axes for the majority of simulations (90%) for trees with low ILS are correlated with minor allele count and reference genome choice (Pearson’s correlation, *p <* 0.01; Fig. 4b), while only 10% of simulations had missing data correlated with separation among trees on either of the first two PCoA axes. For medium ILS trees, results were similar to those in our low ILS trees, but minor allele count and reference genome were more strongly correlated with distance among trees on the first two PCoA axes: minor allele count was strongly correlated with the first PCoA axis for 100% of simulations and reference genome identity was correlated with one of the first two PCoA axes for 90% of simulations. For high ILS trees, we found that the first two axes were correlated with minor allele count and reference genome in 100% of simulations, while missing data was correlated with one of the first two axes in 60% of simulations. The first two PCoA axes explained an average of 54.9% of the variation in the data for `RAxML`-inferred phylogenies (high ILS trees, 73.9%; medium ILS trees, 36.8%; low ILS trees, 54.0%), and an average of 81.9% of the variation in the data for `ASTRAL`-inferred phylogenies (high ILS trees, 77.3%; medium ILS trees, 82.4%; low ILS trees, 86.1%). In treespace for the `ASTRAL`-inferred phylogenies, the first PCoA axes were generally correlated with MAC threshold (96% of simulations), the second PCoA axes were generally correlated with reference genome identity (50% of simulations) and missing data (16.7% of simulations; Fig. S18).

**Fig. 4.**
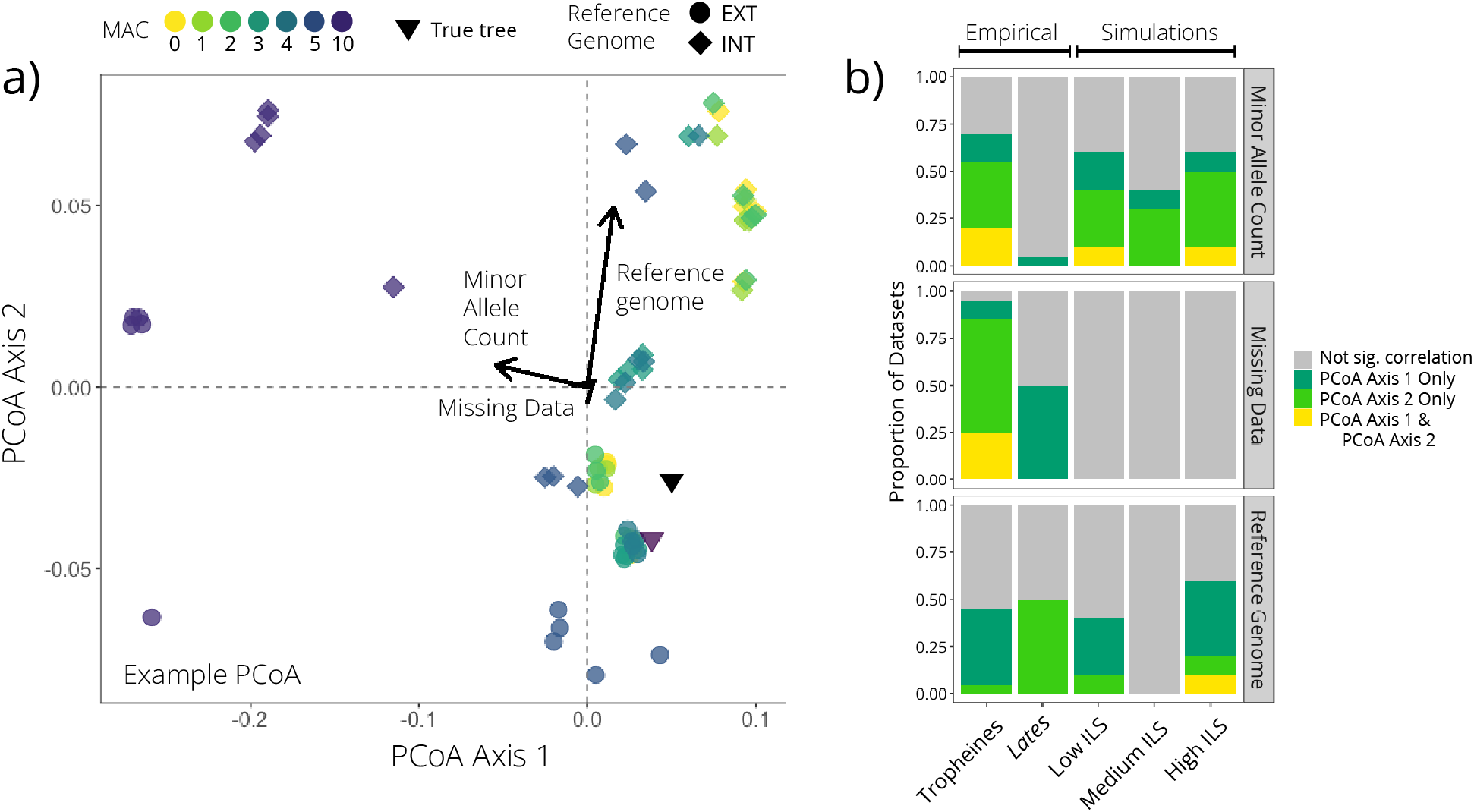
Principal coordinate analyses (PCoAs) based on pairwise Robinson-Foulds distances between `RAxML`-inferred trees demonstrate the relationship between simulated and true trees in two dimensional tree space and emphasize the role of minor allele count (MAC) threshold, missing data threshold, and reference genome on tree topologies. (a) Example PCoA plot of trees from one iteration of simulated data (high ILS, simulation 18), with the true tree indicated and correlations between PCoA axes and each of reference genome choice, MAC threshold, and missing data threshold visualized via arrows. (b) Summary of correlations between each of the three bioinformatic choices and the first two PCoA axes for all simulated and empirical iterations, demonstrating the relative importance of minor allele count in the simulated data and missing data in the empirical data sets, while reference genome was important for all. Barplots show the proportion of iterations for each data set type that had a significant correlation with each of the first two PCoA axes, assessed using Pearson’s correlation coefficients. Dark green, *p*-value *<*0.01 for axis 1; light green, *p*-value *<*0.01 for axis 2; yellow, *p*-value *<*0.01 for both axis 1 and axis 2; gray, *p*-value ⩾ 0.01 for both axis 1 and axis 2. Correlation coefficients are visualized in Fig. S16 and plots for `ASTRAL`-inferred trees are in Fig. S17.

As with the full data sets, both MAC and the interaction between MAC and reference genome distance were significant predictors of topological accuracy in our linear models for our subsampled data sets, where we controlled for the correlation between MAC and data set size by subsampling all data sets to 6,000 SNPs (Fig. 5a; Table S2). Within the subsampled data sets, there was an even clearer improvement in tree topology at moderate MAC thresholds over low or high MAC thresholds, with all three distance metrics indicating that the most accurate topologies for all tree heights and both reference choices occurred at MAC = 3–4 (Fig. S22a,c,e). The subsampled data sets also indicate that in data sets without a MAC filter (i.e., MAC=0), trees constructed with the ingroup reference genome are closer to the true tree than those using the outgroup reference genome for high and medium ILS trees (Fig. 5b).

**Fig. 5.**
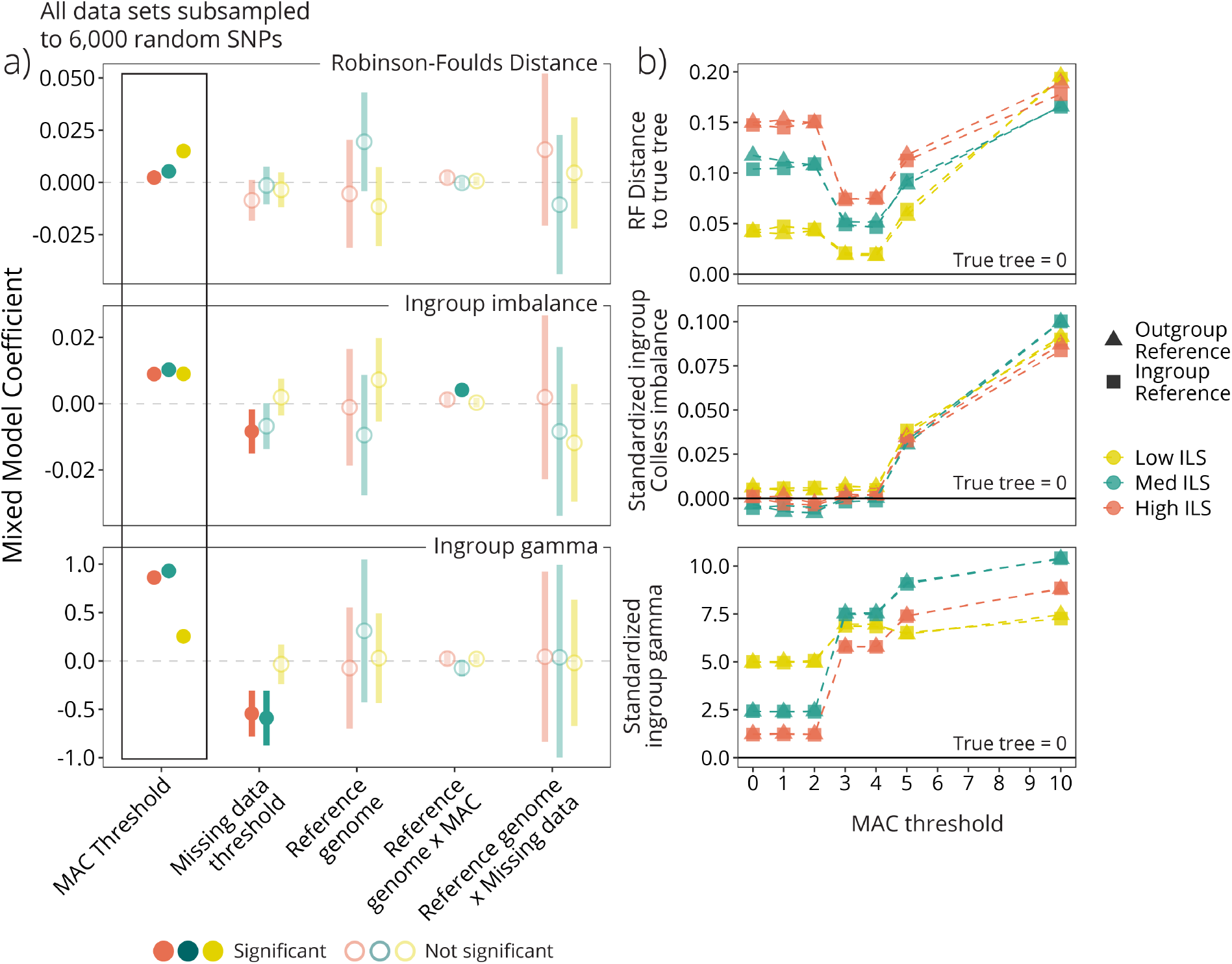
Models for data sets subsampled to 6000 SNPs supported the role of minor allele count (MAC) threshold as a strong predictor of tree topology and shape. We conducted these analyses to address the role of dataset size versus filtering *per se* in influencing downstream results. Subsampled datasets—similar to the findings for datasets differing in size (e.g., Fig. 3)—support a role for MAC’s influence on topology and inferred imbalance. This suggests that MAC filtering does bias the characteristics of the retained SNPs, in addition to its role in reducing the number of retained SNPs. Plots in (a) show model coefficients for linear mixed models for the given tree statistic. Color intensity indicates whether coefficients for the given variable in our linear mixed models were significant (filled in and darker circles) or not significant (faded and open circles). For all three tree statistics, values closer to 0 indicate trees that are closer to the true tree. In (b), the trends in summary statistics across MAC filtering thresholds demonstrate that all three become increasingly distant from the true tree statistic as the threshold for minimum MAC is increased (i.e., with more stringent filtering).

#### Phylogenetic tree imbalance

Our inference of tree imbalance also varied widely across filtering parameters and reference genome choices, and additionally showed different patterns between `RAxML`- and `ASTRAL`-inferred trees. For the full data sets using `RAxML`, minor allele count (MAC) was the strongest predictor variable in our models (Fig. 3a; Table S1) and the linear mixed models explained slightly over half of the variation in the data (mean pseudo-*R*^2^ = 0.566, range 0.466–0.669). The average distance of the reference genome to the ingroup taxa was only an important predictor in our models for imbalance when all tree heights were combined, such that an increased distance to the reference genome increased *I_c_*. In high ILS trees, the interaction between MAC and the distance to the reference genome was also a significant predictor of tree imbalance (Fig. 3b). Filtering for missing data tended to reduce imbalance in the inferred trees, but the effect of this filter on ingroup imbalance was not significant. Both low and medium ILS trees were closest to the true tree imbalance statistic at 0 ⩽ MAC ⩽ 2, while the high ILS trees were closest to the imbalance of the true tree at 3 ⩽ MAC ⩽ 4 (Fig. 3b). These patterns were the same for both Colless and Sackin measures of imbalance (Fig. S22).

In contrast, linear mixed models for imbalance with the `ASTRAL` trees showed differing patterns for the three categories of ILS. With all three ILS levels combined, only missing data was a significant predictor of imbalance, with lower missing data thresholds (i.e., less stringent filtering) producing trees with lower imbalance (Fig. S5f). This pattern is driven predominantly by the effect of missing data in low ILS trees, although the trend is similar in medium and high ILS trees as well (Fig. S5h). In high ILS trees, the reference genome identity also interacted with both missing data and MAC, such that the pattern across MAC and missing data differed between data sets aligned to the ingroup versus outgroup reference genome. In examining effect plots (Fig. S26), the imbalance for all three tree heights across both reference genomes was closest to that of the true tree at 3 ⩽ MAC ⩽ 5, although this effect is weak when compared to the `RAxML` results.

In the subsampled data sets, MAC again was the largest driver of standardized ingroup tree imbalance, with higher MAC thresholds resulting in more imbalanced trees and trees farther from the true tree statistic (Fig. 5; Table S2). The interaction term between MAC and distance to the reference genome was an important predictor again, but the relationship was the inverse of the trend seen in the full data sets (i.e., the interaction term was negative in the full data sets, but positive in the subsampled data sets). In addition, missing data had a significant influence on imbalance in the subsampled data sets, such that increasing the missing data threshold reduces tree imbalance (Fig. S25b,d), a trend that was weaker but in the same direction in the full data sets (Fig. S23b,d). In the subsampled data sets, trees of all three ILS levels and both reference genome types were closest to the true tree imbalance at 3 ⩽ MAC ⩽ 4, as well as when the missing data threshold was most stringent. However, the range of *I_c_* statistics across all missing data thresholds was only *∼*0.005 for high and medium ILS trees, and only *∼*0.001 for low ILS trees, in contrast to the greater variation in *I_c_* for all tree heights across MAC thresholds (*∼*0.05).

#### Influence of parameters on gamma

As stated previously, we are only able to calculate *γ* for the `RAxML`-inferred trees and not those inferred in `ASTRAL`, as `ASTRAL` does not infer terminal branch lengths. Similar to topological accuracy and imbalance, the linear mixed models for standardized *γ* of the ingroup explained a large proportion of the variation in the data for the `RAxML` trees (mean pseudo-*R*^2^ = 0.782, range 0.727-0.823) and had MAC as the strongest predictor of deviation from the true tree’s *γ* statistic (Fig. 3), which was the only significant predictor variable in any of our *γ* models. Across all three tree heights, the values for *γ* were closest to the true tree at 0 ⩽ MAC ⩽ 2 and increased with increasing MAC (Fig. 3b). The change in *γ* across MAC values was greatest in the high and medium ILS trees, and lowest in the low ILS trees, although the low ILS trees were the most biased away from the true *γ* statistic across all MAC thresholds. The reference genome had little influence on *γ* on its own or in interactions with MAC or missing data. Increased missing data thresholds tended to reduce *γ* estimates, although the relationship was weak and the variation in *γ* statistics across missing data values was *<*1 (Fig. S23f).

We see similar patterns for *γ* statistics in the subsampled data sets, with both MAC and missing data being strong predictors of deviation from the *γ* of the true tree (Fig. 5). As with the full data sets, increasing the stringency of the MAC filter increased ingroup *γ*, while increasing the stringency of the missing data filter reduced *γ*. The bias introduced by both filters was strongest in high and medium ILS trees, while low ILS trees had the most consistent *γ* values across filter thresholds (Fig. S24f,S25f). Across all filtering combinations, the inferred topologies tended to be biased to more positive *γ* values than the true trees, which were simulated under a pure-birth process and thus had *γ ≈* 0, or just smaller than 0 (Fig. 2). However, values for *γ* in medium and high ILS trees at 0 ⩽ MAC ⩽ 2 ranged from *γ*=-6.00 to *γ*=-1.00, with a mean *γ*=-4.4 in high ILS trees and mean *γ*=-3.2 in medium ILS trees. At MAC=10, these means then became significantly positive: in high ILS trees, mean *γ*=3.6 (range 1.2-6.8) and in medium ILS trees, mean *γ*=5.25 (range 0.3-7.9). In contrast, low ILS trees only varied from a mean *γ*=-0.4 (range -2.8-1.5) at 0 ⩽ MAC ⩽ 2 to a mean *γ*=1.9 (range -1.08-5.6) at MAC=10.

In the *γ* models including all tree heights, we find that the distance to the reference genome and the interaction between MAC and distance to the reference genome were strong predictors in our models (Fig. 5a). However, neither term is significant in any of our three models for individual tree heights (Fig. 5a), suggesting that the strength in the all-height model may be an artifact of the correlation between mean distance to the reference genome and tree height (which is not included as a predictor in these models; Fig. S3), rather than being due to the distance to the reference genome itself. This hypothesis was supported by a lack of difference between *γ* for trees aligned to the ingroup versus outgroup reference genomes within each tree height across MAC (Fig. S24f) or missing data (Fig. S25f)

#### Influence of parameters on mutation spectra

The patterns we observed in tree statistics suggest that there may be a bias in the characteristics of SNPs lost across filtering parameters, and therefore we examined the mutation rates of SNPs lost at each MAC and missing data cutoff, as well as how each of these varies across reference genome choice. We found that SNPs lost at low MAC and missing data cutoffs were biased toward those located on high mutation rate loci (Fig. 6a), which resulted in MAC and missing data filters truncating the mutation spectra of retained SNPs toward those with lower mutation rates. The change in mean mutation rate of retained loci was most dramatic from MAC=2 to MAC=3, and from miss=0.75 to miss=0.9 (Fig. 6).

**Fig. 6.**
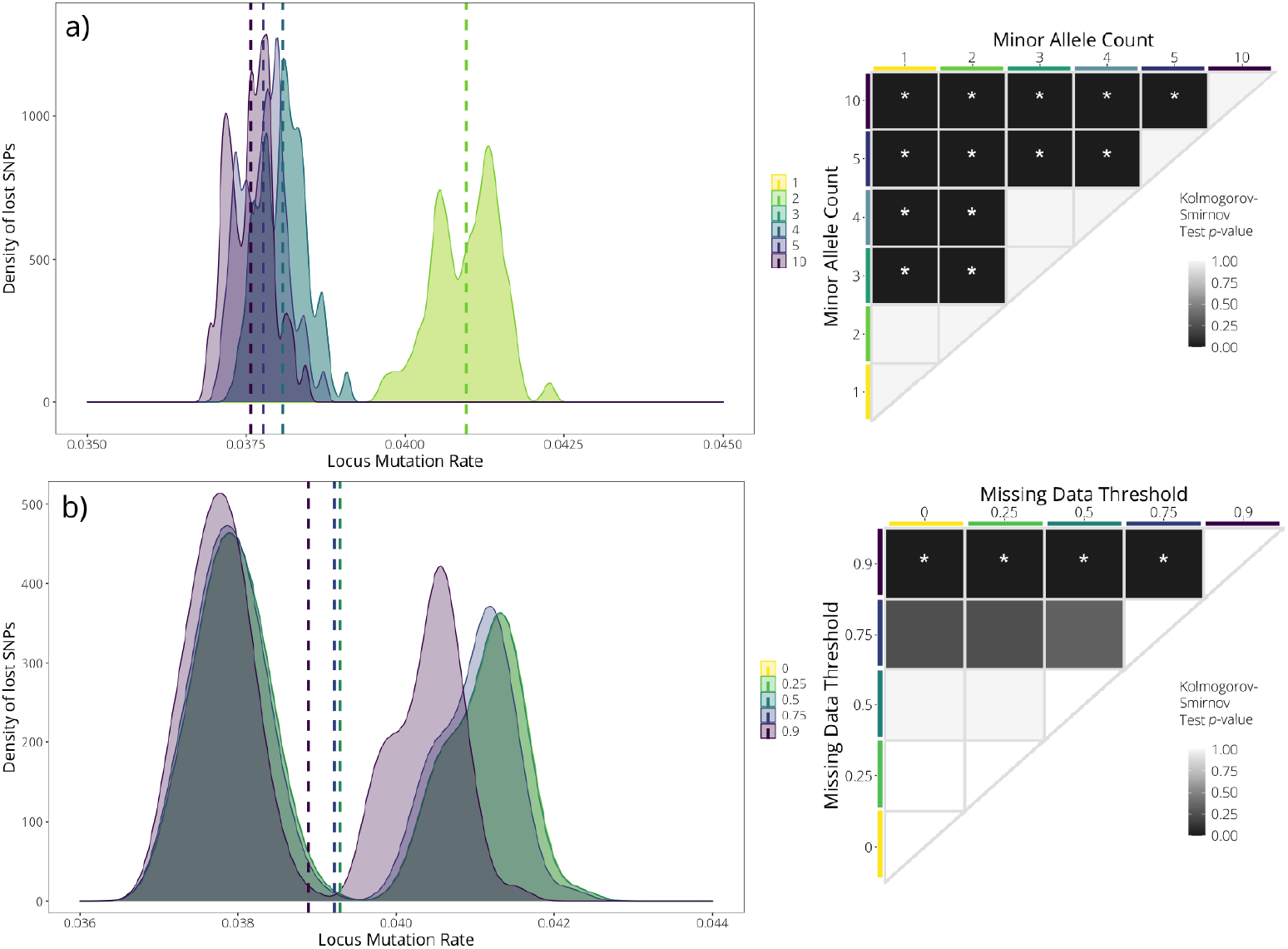
In our simulations, SNPs that were lost first were biased towards those on loci with higher mutation rates for both minor allele count (MAC; a) and missing data (b) filters. Mean values for distributions are indicated with dashed lines, with colors corresponding to minor allele or missing data threshold. Righthand panels show significance from Kolmogorov-Smirnov tests on the distribution of values for each MAC and missing data threshold, with asterisks denoting significant differences (*p <*0.01) among distributions. Note that some distributions are completely overlapping with one another, and thus not visible (i.e., MAC=1 and MAC=2; MAC=3 and MAC=4; miss=0 and miss=0.25). Mutation rates were calculated as the number of variable sites simulated per 5000bp locus.

We also found that the SNPs lost due to MAC filters were those that provided support for nodes toward the tips of the tree (Fig. 7a). We found that the lost nodes had fewer descendants (i.e., were closer to the tips of the tree) than expected based on a null distribution, and the lost nodes were also those that had lower nodal bootstrap support originally (Fig. 7b). As the MAC threshold increased (i.e., became more stringent), more nodes toward the tips of the tree were then lost, which lowered the mean number of node descendants for lost nodes (Fig. 7a). When examining the bootstrap support for nodes that were lost, we find that nodes lost at low MAC cut-offs had lower bootstrap support than those retained, and increasing the MAC cut-off resulted in losing nodes with progressively higher bootstrap support. There were no clear patterns in the number of node descendants for nodes lost across different missing data thresholds (Fig. 7c): the mean number of node descendants of lost nodes remained constantly lower than that of retained nodes independent of missing data threshold. In contrast, the mean bootstrap support of lost nodes decreased with increasing missing data thresholds (Fig. 7d), indicating that low-support nodes are being filtered out at high missing data thresholds, but those lost at lower thresholds are not necessarily the ones with the lowest bootstrap support.

**Fig. 7.**
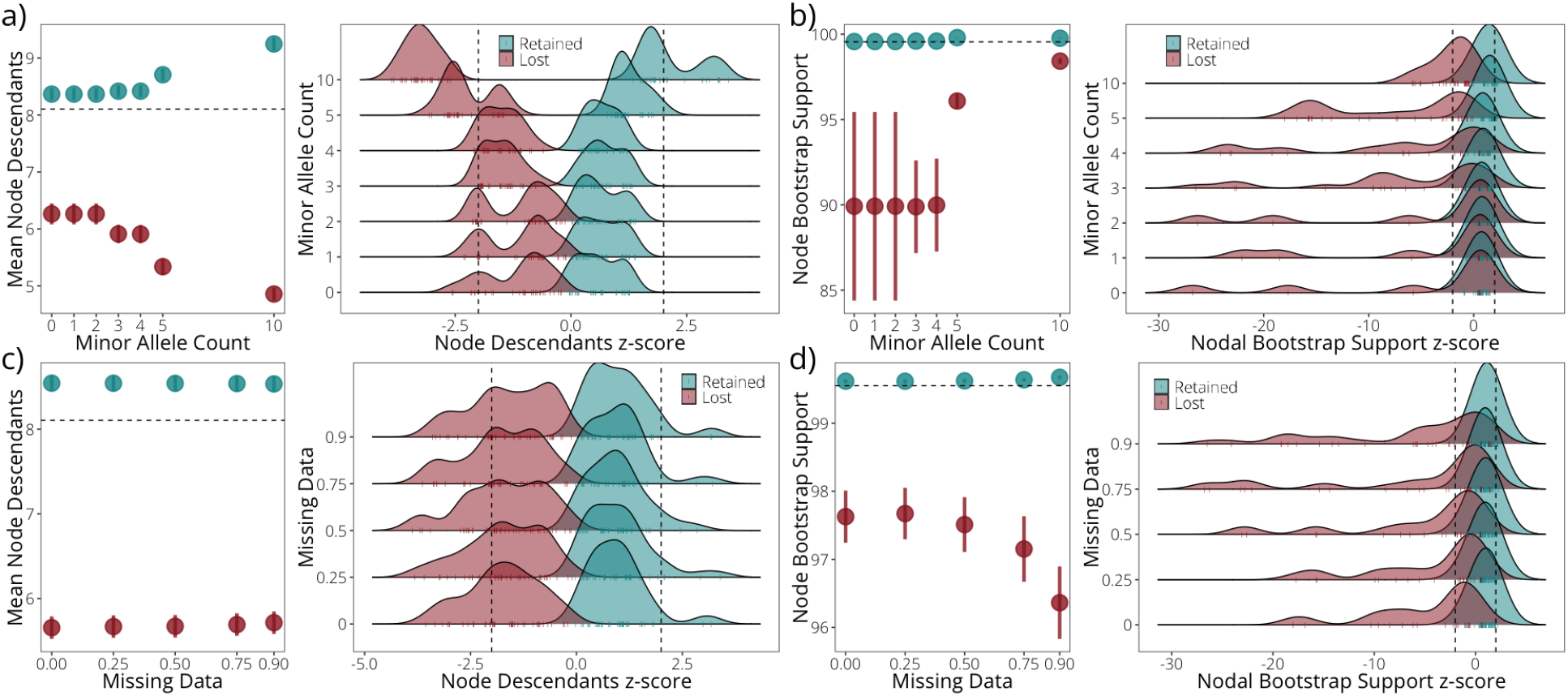
Nodes that were lost from trees with stringent minor allele count (MAC) and missing data filters tended to be those with fewer node descendants. Nodes lost with MAC and missing data filtering also had lower bootstrap support than retained nodes. a) Mean number of node descendants for nodes lost (red) and retained (blue) at each minor allele count threshold, as well as the *z*-scores for the deviation of these means from a null distribution for nodes randomly drawn from the species tree. b) Nodes with lower bootstrap support were generally lost first at low MAC thresholds. Plots show the mean bootstrap support values for those nodes that are lost at each minor allele count threshold, as well as the distribution of *z*-scores for the deviation of these values from a null distribution of bootstrap support values. c) Mean number of node descendants for nodes lost at each missing data cutoff, demonstrating a generally lower number of node descendants for lost nodes but no clear trend as missing data cutoffs become more stringent. d) Mean bootstrap support values for nodes lost at each missing data cutoff, demonstrating a reduction in mean bootstrap support for lost nodes as the missing data threshold becomes more stringent. Left plots for each panel show means and standard errors, as calculated using the mean se function in ggplot2. Horizontal dashed lines indicate the overall mean bootstrap support and node descendants across all nodes, respectively. In right panels, dashed lines indicate *| z |* = 1.96.

In contrast to the patterns observed across MAC thresholds, the distribution of mutation rates did not differ overall between sites retained in data sets aligned to the internal versus external reference genomes. Additionally, the nodes lost when using an internal versus external reference genome were not biased toward those found in a certain part of the trees.

### Empirical analyses

#### General overview of empirical data sets

The ten replicates of each of our two empirical clades (drawing 51 random individuals each replicate) resulted in 2,800 data sets with an average of 82,978 SNPs (tropheines; range 311–550,214) and 730,239 SNPs (*Lates*; range 1,147–16,018,003). The average distance of ingroup taxa to the reference genome for the 40-species tropheine clade’s distant reference (*Oreochromis niloticus*) was *d_xy_*=0.517, and for the close reference genome (*Pundamilia nyererei*) was *d_xy_*=0.161 (Fig. S7). For the 4-species *Lates* clade, the average reference genome distances were *d_xy_*=0.602 (outgroup reference, *Lates calcarifer*) and *d_xy_*=0.0669 (ingroup reference, *L. mariae*; Fig. S7). As with our simulated data, the number of SNPs retained was highly inversely correlated with MAC cutoff (Pearson’s *r*(1292) = -0.141, *p* = 3.5*e −* 07) and with missing data threshold (Pearson’s *r*(1292) = -0.237, *p <* 2.2*e −* 16). The number of SNPs retained was also correlated with overall tree height for the `RAxML` trees (Pearson’s *r*(1292) = 0.384, *p <* 2.2*e −* 16). Ingroup tree height was correlated with ingroup Colless and Sackin imbalance, although the correlation was negative in the tropheines and positive in the *Lates* clade (tropheines: Pearson’s *r*(593) = -0.275, *p* = 8.287*e −* 12; *Lates*: Pearson’s *r*(698) = 0.114, *p* = 0.0024). In contrast to the simulations, tree height was not correlated with ingroup *γ* in either empirical clade (tropheines: Pearson’s *r*(593) = 0.006, *p* = 0.883; *Lates*: Pearson’s *r*(698) = -0.012, *p* = 0.746).

#### Tropheine analyses

For the two empirical data sets and the phylogenies inferred using `RAxML`, we modeled ingroup tree height, ingroup Colless imbalance (*I_c_*), and *γ* as a function of minor allele count, missing data threshold (miss), and reference genome identity (ref), as well as the interaction between reference genome choice and each of minor allele count and missing data thresholds. In contrast to the simulation results, the reference genome was treated as a binary choice (ingroup/close reference versus outgroup/distant reference), with the more distant genome coded as 0 and the close genome coded as 1. In our phylogenetic inferences for the tropheine data, our models explained the largest proportion of the variation in the data for *γ* (pseudo-*R*^2^=0.800), less variation for imbalance (pseudo-*R*^2^=0.337), and tree height (pseudo-*R*^2^=0.432). We found that MAC threshold was an important predictor of ingroup tree height, imbalance, and *γ*, and missing data threshold was an important predictor of ingroup imbalance and *γ* (Fig. 8a). As the MAC cutoff became more stringent, the inferred ingroup tree height increased and *γ* increased to become more positive (Fig. 8b). Imbalance decreased as MAC increased from MAC=0 to MAC=3, and then the imbalance increased again between MAC=5 and MAC=10 (Fig. 8b). With increasingly stringent missing data thresholds, ingroup tree height increased and then decreased, ingroup imbalance decreased, and ingroup *γ* decreased (Fig. 8b).

**Fig. 8.**
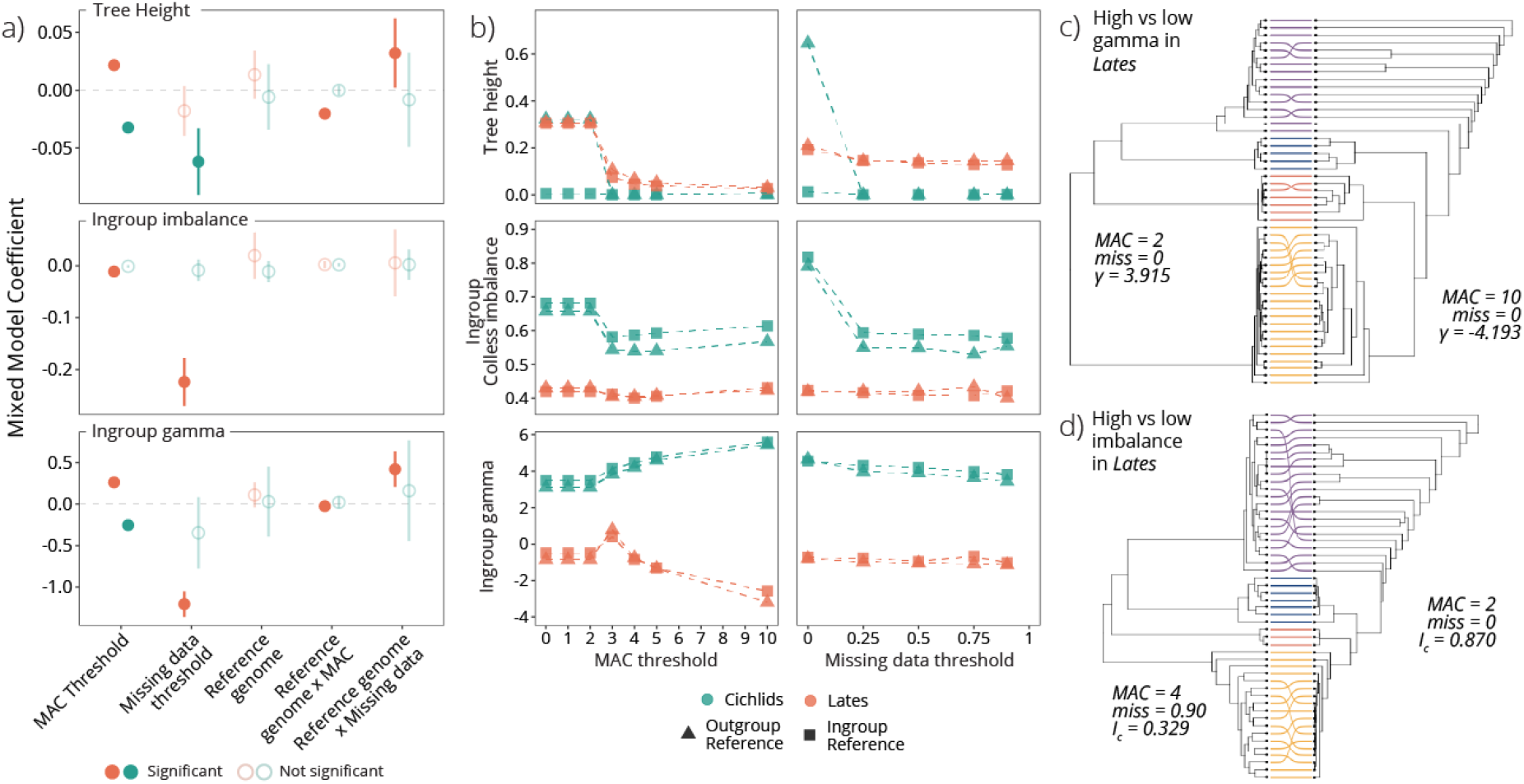
In the empirical data sets, missing data, minor allele count (MAC), and the choice of reference genome all interacted to affect the inferred tree height, imbalance, and *γ* statistics. Plots in (a) show model coefficients from linear mixed models for empirical data sets, with MAC threshold, missing data threshold, reference genome choice, the interaction between missing data threshold and reference genome choice, and the interaction between MAC threshold and reference genome choice as explanatory variables, and tree height, imbalance, and gamma (*γ*) in the `RAxML`-inferred phylogenies as response variables. Results are shown for tropheine (orange) and *Lates* (teal) empirical data sets, with univariate responses across MAC and missing data thresholds shown in (b) for the three response variables. Example trees are shown from the *Lates* data demonstrating the differences between trees at the two extremes for (c) the *γ* statistic, and (d) Colless’ imbalance statistic. In each pair of trees, datasets differ only in the filtering parameters used, as indicated. In (c) and (d), colors indicate the species identity of each individual, with lines matching up the same individual between the left and righthand trees. Note that in both cases, one of the species (in purple) becomes non-monophyletic in the tree on the right, which leads to a large change in tree statistics.

We additionally found an influence of reference genome on ingroup tree height and *γ*, via interactions between the reference genome and our filtering parameters (Fig. 8a). In the tropheine topologies, using the outgroup reference genome tended to increase tree height compared to the ingroup reference genome, particularly at high MAC thresholds. Using the ingroup reference genome also resulted in trees that were more imbalanced, while the outgroup reference genome produced trees that had higher *γ* statistics than trees constructed using the ingroup reference genome. At low MAC thresholds, trees using the ingroup reference were more tip-heavy than those using the outgroup reference, but had similar *γ* values to trees using the outgroup reference at higher MAC thresholds (Fig. 8b). Similarly, ingroup reference trees had similar *γ* statistics to outgroup reference trees when all missing data was allowed (miss = 0), but then had higher *γ* statistics than the outgroup trees at higher missing data thresholds (Fig. 8b).

In analyses of tree space based on pairwise RF distances between tropheine phylogenies inferred in `RAxML`, trees generally clustered together primarily based on missing data, with additional influence of reference genome and MAC (Fig. 4). The first two PCoA axes were generally correlated with missing data (100% of replicates on axis 1) and reference genome choice (80% on axis 2), and a few iterations also had strong correlations between these first two axes and MAC (70% on axis 1 and 30% on axis 2; Fig. 4b). The first two PCoA axes combined for `RAxML`-inferred tropheine phylogenies explained an average of 91.8% of the variation in the data.

The linear mixed model for Colless imbalance in the `ASTRAL` tropheine trees explained just under half of the variation in the data (pseudo-*R*^2^ = 0.448), and indicated that imbalance differences between the trees were driven predominantly by the missing data cutoff and the interaction between the reference genome choice and missing data cutoff (Table S5; Fig. S6), with more stringent missing data cutoffs leading to less imbalanced trees (Fig. S27). Supporting this, the first PCoA axis for tropheine trees inferred in `ASTRAL` was generally significantly correlated with the missing data cutoff (70% of iterations; Fig. S21a), while minor allele count was associated with the first axis in four replicate data sets and the second axis in four of the ten replicate data sets. The first two PCoA axes combined for `ASTRAL`-inferred tropheine phylogenies explained an average of 72.2% of the variation in the data.

#### Lates analyses

In our analyses of the *Lates* topologies, we found that our models explained the largest amount of variation in the data for imbalance (pseudo-*R*^2^ = 0.823), less variation for height (pseudo-*R*^2^ = 0.567), and the smallest amount of variation for *γ* (pseudo-*R*^2^ = 0.261). The MAC threshold was again an important predictor of ingroup tree height and ingroup *γ* (Fig. 8a). However, the relationship between MAC and both tree height and *γ* was in the opposite direction to the tropheines and the simulated data: increased stringency in the MAC filter reduced ingroup tree height and reduced ingroup *γ* values, except for a brief increase in *γ* at MAC=3 (Fig. 8b). Similar to the tropheines, imbalance decreased slightly from 0 ⩽ MAC ⩽ 2 to 3 ⩽ MAC ⩽ 5, and then increased again at MAC=10 (Fig. 8b).

Missing data was also an important predictor of ingroup height in the *Lates* data, such that increasingly stringent missing data filters resulted in shorter trees (Fig. 8b). The reference genome choice, as well as the interaction between reference genome and our minor allele and missing data filters, was not an important predictor in any of our models with the *Lates* data (Fig. 8a; Table S3), although differences among trees aligned to different reference genomes can be seen at some filtering thresholds (Fig. 8b).

*Lates* trees generally clustered together in PCoA tree space primarily based on reference genome on the first PCoA axis and missing data on the second PCoA axis (Fig. 4b; S21b). The MAC filter only correlated with one of the two main PCoA axes in one of the ten *Lates* replicate data sets. The first two PCoA axes combined for `RAxML`-inferred *Lates* phylogenies explained an average of 42.2% of the variation in the data.

Filtering parameter choices tended to have a greater influence on the characteristics of the *Lates* trees inferred using `ASTRAL` than those inferred using `RAxML`. The MAC threshold, missing data cutoff, and reference genome identity all were identified as significant predictors in our model for ingroup imbalance (Table S5; Fig. S6b). In PCoA space, the *Lates* trees inferred using `ASTRAL` tended to cluster based on missing data cutoff on the first PCoA axis and MAC threshold on the second, while reference genome was correlated with the second PCoA axis in only two of the ten replicate data sets (Fig. S21a). The first two PCoA axes combined for `ASTRAL`-inferred *Lates* phylogenies explained an average of 43.2% of the variation in the data.

## Discussion

Single nucleotide polymorphism (SNP) data sets are used extensively in population genomic and phylogenomic studies, but best practices in the bioinformatic choices made while aligning and filtering these data remain unclear for both reduced-representation and whole genome sequencing data sets. There is no generally accepted analysis pipeline for these data, and it is unclear whether a “one size fits all” approach is warranted. Our results from a combination of simulated and empirical analyses demonstrate how decisions made by researchers during the assembly and processing of genomic data, as well as characteristics of the true divergence history among taxa, influence the final data set and the inferred phylogenetic and diversification history. We found that the value chosen for minor allele filters interacts with the choice of reference genome and total tree height (and thus ILS) to bias phylogenetic inferences. This suggests that removing rare variants by retaining only sites that have a minor allele count above a certain threshold reduces phylogenetic accuracy, particularly toward the tips of a tree. In contrast, missing data filters can increase the topological accuracy of the inferred phylogenetic tree, although this desirable behavior may be countered by an undesirable truncation of the mutation spectrum. We also examined the effect these parameter choices have on tree balance and shape. In our simulations, we found that increasing the minor allele threshold (and thereby removing rare variants) generally reduces the accuracy of tree shape and balance statistics: more stringent filtering results in increased topological distances, increased imbalances, and increased *γ* statistics of inferred trees compared to the true tree. Our results complement and extend those from other recent studies—which focused mostly on single filtering and alignment parameters of interest (e.g., Linck and Battey, 2019; Huang and Knowles, 2016; Prasad et al., 2021; Günther and Nettelblad, 2019; Shafer et al., 2017; Molloy and Warnow, 2018; Lanier et al., 2014)—by demonstrating that the choice of reference genome can also bias phylogenetic analyses, as well as exacerbate or modify the effects of individual filtering parameters. Furthermore, a number of these effects are independent of the overall size of the final data sets, suggesting that it is the nature of the remaining data—not merely the volume—that biases phylogenetic inference.

### Stringent minor allele filters bias topological inference and diversification statistics

Most variation in natural populations is rare, and rare variants are those that contain the most information about differentiation between individuals or populations (Slatkin, 1985; Nelson et al., 2012); thus it is perhaps unsurprising that we found a clear pattern that stringent minor allele filters bias inferred topologies. This was true in both our full data set, and in the subsampled data sets that were used to control for data set size. Stringently filtered trees tended to be more imbalanced and more tip-heavy (i.e., had a less negative *γ*) than the true species trees. The topologies closest to that of the true tree and with imbalance closest to that of the true tree were at minor allele count (MAC) 3–4. This suggests that excluding sites with very low-frequency alleles (*<*3 copies across all samples in the data set, corresponding to a minor allele frequency of 1–3%) leads to more accurate results in phylogenetic analyses, likely by removing SNPs that represent sequencing or alignment errors. However, higher thresholds remove true phylogenetic signal from the data set, reducing data quality and biasing inferences toward trees that are more pectinate than the true phylogenetic history. This suggests that using minor allele frequency filters of 5-10%—common thresholds in empirical studies—may be unknowningly biasing evolutionary inferences.

In contrast to the patterns observed with imbalance and topological accuracy, our inferred topologies had ingroup *γ* statistics closest to that of the true trees at the lowest MAC thresholds (MAC=0–2) and became much more biased at MAC=3–4. Our simulated trees were created using a pure birth process, which produced *γ* statistics that follow a standard normal distribution (with a mean *γ ≈* 0 and sd=1). The magnitude of change between phylogenies filtered with MAC=0 and MAC=10 was great enough to change *γ* from significantly negative (*γ* = -5.73) to significantly positive (*γ* = 7.84; Fig. S8–S10), and the mean absolute change in *γ* was 3.28 for low ILS trees, 8.55 for medium ILS trees, and 8.02 for high ILS trees. This suggests that the MAC filter alone could dramatically change inferences of diversification rates through time and, importantly, lead to incorrect macroevolutionary inferences at high MAC thresholds.

These findings for imbalance and *γ* statistics follow from theoretical expectations that recent splits in the phylogeny have less shared variation supporting them (i.e., more low-frequency SNPs or rare variants), and thus are disproportionately lost when using a MAC filter (Linck and Battey, 2019). This is supported by our findings that the majority of nodes lost due to the minor allele filter were those with few node descendants (Fig. 7a) and therefore less hierarchical redundancy (sensu Eaton et al., 2017). The SNPs lost at lower MAC thresholds also tended to be those with higher mutation rates (Fig. 6). At 0 ⩽ MAC ⩽ 2, we thus gained topological accuracy but also shortened branch lengths near the tips of the trees, shifting the tree’s center of gravity tipward and increasing *γ* estimates. This process may have also driven the increase in tree imbalance: the loss of more recently evolved SNPs removes lineage-specific variation from nested clades, which then may lead to more pectinate topologies. Because of where low-frequency SNPs are found in the tree, these biases we observe in imbalance and *γ* may be particularly problematic in trees with rapid recent divergence.

Some MAC-related bias was more apparent in our subsampled data sets than our full simulations, suggesting an effect of the overall data set size on the general trends in our parameters of interest described above. The most clear of these was the shape of the topological accuracy response across MAC thresholds: the full data sets showed a slight increase in topological accuracy from MAC=1–2 to MAC=3–4 (Fig. 3b), while the subsampled data sets showed a much more dramatic increase in accuracy at MAC=3–4 (Fig. 5b). This suggests that the deviation from the true tree when MAC ⩽ 2 in the subsampled data sets may have been due to having fewer SNPs (i.e., less data) than the full data sets, and therefore the signal from any “erroneous” SNPs (e.g., those that result from sequencing errors) was stronger and more likely to bias the inferred tree. Thus, filtering out these true sequencing or alignment errors did improve topological accuracy. In this way, our results suggest that using a MAC filter is more important in data sets with fewer overall SNPs, although this relationship will need to be explored further in future studies.

These results together suggest that trade-offs exist between reducing SNP errors, improving topological accuracy, and improving diversification statistic accuracy. The general consistency observed between results for the full data sets and results for the subsampled data sets suggests that the numerical loss of SNPs when using more stringent filters was not the only factor driving these patterns, but rather that there are biases in the characteristics of the SNPs being lost. This also suggests that caution should be used in evaluating diversification statistics in trees inferred using heavily filtered data. Our results echo others’ findings that filtering based on minor allele count or frequency can bias population genetic inferences, especially for events in the recent past (Linck and Battey, 2019; Shafer et al., 2017; Boitard et al., 2016), and provides reason for using caution when interpreting macroevolutionary patterns from SNP-based phylogenomic datasets.

Some of the patterns in our empirical data mirror our observations from the simulations. The MAC filter also had a strong influence on the inferred topology, tree height, and *γ* for our empirical phylogenies, although the patterns differed between the tropheine and *Lates* data sets. The change in *γ* with MAC was less evident in our empirical data sets than simulated data sets; however, ingroup *γ* decreased in the *Lates* phylogenies, transitioning from a non-significant *γ* estimate to significant and negative *γ* at more stringent MAC thresholds (Fig. 8b, S15). Thus, macroevolutionary inferences in this clade would differ widely at different MAC thresholds (as demonstrated in Fig. 8c). In contrast, estimates of *γ* increased slightly with increasing MAC filters in the tropheines, although it remained significantly positive among all MAC thresholds and thus would not change macroevolutionary inferences (Fig. 8b, S14). Whereas imbalance for the simulated data sets increased across all MAC thresholds, imbalance actually decreased for both empirical data sets from MAC=0 to MAC=3, and then increased again above MAC=5 (Fig. 8b). We do not know the true value for these data sets and cannot know whether the values we estimated deviate from the true statistic. However, the consistency of the observed inflection point at mid-range MAC thresholds may suggest that these data sets are also improved in this range. In examining the trees with extreme *γ* and imbalance statistics for the empirical data sets, it appears that trees with the highest imbalance involve non-monophyly of groups expected to be monophyletic (Fig. 8d; S4), suggesting that these extreme values are not representative of the true evolutionary relationships. Similarly, the trees with extremely low *γ* in the *Lates* data sets involve non-monophyly of one of the species (Fig. 8c), suggesting again that extreme values represent bias derived from topological inaccuracy.

In both the subsampled data sets and the empirical data sets, there were differences between the responses at MAC*<*3 and MAC*>*4. In the subsampled data sets, this was a general minimum in divergence from the true tree, while the truth is unknown for the empirical data sets. That this threshold would be an inflection point is not surprising: given that these are diploid data sets, SNPs filtered at MAC=3 but not MAC=2 must be shared by at least two tip taxa. Consequently, this is the first MAC filter that must remove shared variation, rather than alleles found in only a single taxon, and thus it is expected to have a broader influence on tree topology and tree metrics.

### Missing data filters affect diversification statistics

In contrast to the patterns with minor allele filters in the simulations, stringent missing data filters (i.e., those that removed sites with lower amounts of missing data) were associated with an improvement in the accuracy of imbalance and *γ* statistics, and had little effect on topological accuracy. The subsampled data sets demonstrated a clear pattern that more stringent missing data filters resulted in imbalance and *γ* statistics closer to the true values (Fig. 5a). Our full data sets trended the same direction, although missing data was not a statistically significant predictor in our models (Fig. 3a).

Missing data had a stronger influence on imbalance and *γ* estimates in our empirical data sets than in our simulated data sets. While we do not know the truth for the empirical data sets, we found a negative relationship between these two statistics and missing data for both the *Lates* and the tropheines (Fig. 8). This likely reflects aspects of our empirical data that are not captured in our simulations, such as biases in which sites are most likely to have high levels of missing data (e.g., allelic dropout due to restriction site mutations, which has been incorporated in other simulation studies that find a strong bias due to missing data; Huang and Knowles, 2016; Eaton et al., 2017). Our simulated data also had low overall rates of missing data: an average of only 0.51% of sites in the unfiltered simulated data had *>*50% missing data (mean per-site missing data = 2.21%). In contrast, our empirical data sets had an average of 97.0% of sites with *>*50% missing data (mean per-site missing data = 94.3%). This discrepancy led to fewer sites being lost due to missing data filters in the simulated data sets, which then would lead to less variation between phylogenies inferred at different missing data thresholds.

Despite the low amount of missing data present in our simulated data sets, we did find that filtering for missing data improved the accuracy of our imbalance and *γ* estimates without much reduction in topological accuracy. Diversification statistic inferences were most accurate for simulated data at our most stringent threshold for missing data (miss=0.9; Fig. S23,S25). This suggests that the reductions in imbalance and *γ* statistics in both of the empirical data sets (Fig. 8b) may indicate more accurate inferences as well. However, the difference in these statistics between miss=0.25 and miss=0.90 was not great enough to change macroevolutionary inferences based on *γ* or imbalance statistics for either clade. In addition, we found that filtering sites for missing data biased the mutation spectrum of the retained loci by disproportionately removing SNPs on loci with the highest mutation rates in the simulation data sets (Fig. 6). Thus, the small possible improvement in imbalance and *γ* accuracy may not outweigh the previously demonstrated drawbacks to filtering stringently for missing data in real-world data sets (e.g., Huang and Knowles, 2016; Eaton et al., 2017; Chan et al., 2020).

### Reference genome choice can bias topological inference and tree imbalance

In addition to minor allele and missing data filters influencing the accuracy of inferred trees, our results suggest that reference genome choice can strongly influence phylogenetic insights gained from short read data. The difference between trees aligned to an ingroup versus an outgroup reference genome was most evident in principal coordinate analyses of Robinson-Foulds distances among all trees that came from the same simulated species tree, where the choice of reference genome was correlated with the first or second PCoA axis in the majority of analyses for both `RAxML`- and `ASTRAL`-inferred trees (Fig. 4b; S18). Similarly, reference genome choice was correlated with the first or second PCoA axis for empirical trees as well, although the effect was stronger in `RAxML` trees than those inferred using `ASTRAL`. However, in PCoA tree space, there is no clear pattern for whether the data sets aligned to the ingroup or outgroup reference genome produced trees closer to the true tree in tree space (among all tree heights, 16.7% of true trees were closer to trees from the ingroup reference, 16.7% of true trees were closer to those from the ougroup reference, and 66.7% were equally close to both ingroup and outgroup reference trees). Our models, on the other hand, suggested that the distance to the reference genome did not have a strong influence on topological accuracy on its own in our simulated data sets. This seeming discrepancy likely arose because absolute Robinson-Foulds distance to the true tree (used in our modeling) describes the one-dimensional magnitude of topological differences, but not *how* a tree is different from the true tree or whether it deviates in the same way that other trees deviate from the true tree. In contrast, PCoAs effectively provide a two-dimensional map of tree space, thus picking up more detail about how similar the inferred trees are to one another and allowing inferences about which trees differ from the true tree in similar ways. In doing so, the PCoA results suggest that reference genome choice may be even more important to the inferred topology than our modeling results are able to detect.

Increasing the average distance of the reference genome to the ingroup taxa increased imbalance in the inferred topologies, although the influence of reference genome on its own was not a significant predictor of imbalance in any of our models. These patterns may have arisen due to biases in the different characteristics of loci retained when using ingroup versus outgroup reference genomes: the former are more likely to retain low-frequency variation within the ingroup and the latter are more likely to retain conserved regions with low mutation rates (i.e., sites where the ingroup taxa are more similar to the reference genome; Ros-Freixedes et al., 2018; Brandt et al., 2015). The loss of low-frequency variants within the taxa of interest when using an outgroup reference genome may lead to a lack of support for nested clades within the phylogeny, which then leads to more imbalanced and pectinate topologies. In addition, genotype calling accuracy for heterozygotes decreases with increased distance to the reference genome (Duchen and Salamin, 2021), and it is possible that this influences our inferred phylogenies as well.

Intriguingly, our tropheine and *Lates* empirical data trended in the opposite direction, with the ingroup reference genome producing trees that were more imbalanced than those using the outgroup reference genome for the `RAxML` trees, while `ASTRAL`-inferred trees tended to be more imbalanced overall (Fig. S27), and more imbalanced when using the outgroup reference than the ingroup reference (Fig. S27). There are several possible explanations for the discrepancies between the simulated and empirical data in the `RAxML`-inferred phylogenies. First, we do not know the true level of imbalance for the evolutionary history for these empirical clades, and thus it is possible that higher imbalance in the tropheines and the *Lates* are less biased imbalance estimates, in which case these results would agree with our simulated data. Second, it is possible that the different responses are related in differences in distances to the reference genomes used or related to differences in taxon sampling. For example, the tropheine clade’s closer reference (*Pundamilia nyererei*) is not actually nested within the clade, but rather is a more closely related outgroup than the “outgroup” reference (*Oreochromis niloticus*). Thus, the mean distance to the close reference genome (Fig. S7) was greater in the tropheine clade (*d_xy_*=0.161) than for the *Lates* ingroup reference (*d_xy_*=0.0669) or the simulation ingroup references (*d_xy_*=0.039). In addition, the 40-species clade of tropheines is likely to have more low-frequency variation (i.e., rare variants) captured than the 4-species clade of *Lates*—and both of these sampling structures differ from our simulations, where we simulated each tip as an individual taxon. An extensive body of work already exists on biases introduced via taxon sampling choices (e.g., Zwickl and Hillis, 2002; Heath et al., 2008b; Hohna et al., 2011; Cusimano and Renner, 2010; Brock et al., 2011), and our results indicate that the choice of reference genome may interact with taxon sampling schemes to bias phylogenetic inferences and diversification statistics. We did find that there is a smaller difference in the mean number of SNPs retained between data sets aligned to the close versus distant reference genomes for the tropheine clade (difference in means = 9,552 SNPs, with more SNPs retained with the close reference; Fig. S29) than in the *Lates* clade (difference in means = 57,643 SNPs, with more SNPs retained with the outgroup reference). These differences in number of SNPs may indicate differences in the characteristics of SNPs retained as well. While we did not detect differences in mutation spectra retained in outgroup versus ingroup data sets, these patterns warrant further investigation.

Finally, it is also possible that reference genome quality—not just distance—systematically affects phylogenetic inferences, such that more complete reference genomes retain more SNPs or that more fragmented genomes tend to retain more conserved variation. The distant reference genomes used in our empirical analyses here were both chromosome-level assemblies (*L. calcarifer*, 24 chromosomes; *O. niloticus*, 22 chromosomes), while the close reference assemblies are not yet assembled into complete chromosomes (*L. mariae*, 1,154 scaffolds; *P. nyererei*, 7,236 scaffolds). This pattern—having a distant, well-assembled genome and a closer, more fragmented genome available—is common when making the choice between a close versus distantly related reference genome. Thus some of the effect of reference genome distance that we observe in our empirical analyses (and perhaps some of the differences between our empirical and simulation results) may be linked to assembly completeness. While genome quality does not appear to bias population genomic demographic reconstruction methods (Patton et al., 2019), it remains unclear whether phylogenetic inferences are similarly unaffected – although this is not something that we were able to test explicitly with our analyses.

If multiple reference genome choices are available, one way to potentially mitigate the bias introduced due to reference genome choice may be to map reads to multiple reference genomes. This approach is recommended by Bertels et al. (2014), who found that the choice of reference taxon affected the percentage of incorrect topologies in their simulations, but demonstrated that this bias was reduced by replicating analyses over multiple reference genomes. Our results support this recommendation, as mapping to multiple reference genomes allows an exploration of how the topology changes when the data are aligned to one reference versus another. Another promising possibility to reduce the biases demonstrated here may be using multiple references to create a “pseudogenome” for the species of interest (Huang et al., 2014), as demonstrated in Sarver et al. (2017). By generating a pseudogenome based on multiple possible references or by incorporating variation from multiple individuals into a reference assembly (such as with a pangenome; Eizenga et al., 2020), reference-based mapping biases may be reduced, particularly when there is great variation among focal species in their phylogenetic distance to the possible reference genomes. Both of these options are increasingly possible due to the acceleration in genome sequencing for non-model taxa (Hotaling et al., 2021; Rhie et al., 2021; Formenti et al., 2022). However, not every clade of interest will have multiple reasonable reference genome options. In this case, our results can be used to inform filtering practices: stringent filtering thresholds (particularly for minor allele count) will bias the results more when using a distant reference genome than when using an ingroup (or closely related) reference genome.

### Magnitude of bioinformatic biases depend on interactions and the true divergence history

No bioinformatic choice is made in isolation, and our results demonstrate that the magnitude of bias introduced by each decision depends both on the true evolutionary history of the clade of interest and on interactions among bioinformatic choices. Generally, our longest `RAxML`-inferred trees were the most resistant to filtering-related deviations in topology and imbalance (Fig. 3b,5b), suggesting that trees with low ILS are the least prone to bias arising from bioinformatic choices. In these longer trees, larger internode distances mean that there are more redundant SNPs supporting each split in the tree and more time for lineage sorting to occur among ancestral polymorphisms, and this redundancy may allow for correct inferences despite biased patterns to the loss of SNPs. The low ILS trees, however, were also generally further from the true *γ* statistic than medium or high ILS trees, despite also having more consistent *γ* statistics across MAC thresholds (Fig. 3b, 5b). The deviation in *γ* at low MAC thresholds for low ILS trees may be due to proportionally more SNPs occurring—and also being lost—along the tip branches in the low ILS trees, but more work is needed to interrogate this effect.

The greatest difference observed between `RAxML`- and `ASTRAL`-inferred trees in our simulations was in how the low ILS data sets performed at high MAC thresholds. While high and medium ILS trees showed similar patterns across both MAC and missing data thresholds, `ASTRAL` inferred much more inaccurate trees at high MAC thresholds for the low ILS trees, as measured by all three topological distance measures (Fig. S26a–f). For imbalance measures, `ASTRAL`-inferred trees were closer to the true tree imbalance at high MAC thresholds than their `RAxML`-inferred counterparts across all three tree heights (Fig. S26g–j). While other results were largely concordant among inference methods, these differences suggest that the choice of inference method can additionally interact with the clade history, alignment choices, and filtering choices, to influence evolutionary interpretations. It is possible that the poorer performance of `ASTRAL` at high MAC thresholds is due to low numbers of SNPs per locus leading to loss of signal among gene trees, which then leads to poor inference of species trees. Thus, despite species tree methods being ILS-aware and generally performing better than concatenation methods for data sets with high ILS, our results suggest that they are not a solution to mitigating all of the biases found in our study.

Our two empirical data sets, which had different amounts of gene tree incongruence, different imbalances, and different diversification histories, demonstrate how these bioinformatic filters can interact with a clade’s evolutionary history to influence phylogenetic inference. Overall, we saw much more dramatic responses in tree characteristics in the *Lates* trees (particularly tree height and *γ*; Fig. 8b) than in the tropheine trees across minor allele thresholds, while missing data had a greater effect on the tropheine trees than the *Lates* (particularly for tree height and *γ*; Fig. 8b). While we do not know the “true” statistics for the empirical analyses, we found that *γ*, imbalance, and overall tree height all showed an inflection point around MAC=3–4, as was the case with our simulated data. However, the shape of these statistics’ responses to MAC across this inflection point differed between the tropheine, *Lates*, and simulated trees (Fig. 3b,5b,8b). In the *Lates* analyses, the highest *γ* values were at moderate minor allele and moderate missing data thresholds. In the tropheine data set, on the other hand, *γ* increased slightly with increasing minor allele thresholds and decreased with increasing missing data thresholds.

These interactions suggest that the best filtering scheme will differ for data and clades with different characteristics (e.g., the rapidity of diversification within the clade, the amount of incomplete lineage sorting, the level of missing data or sequencing errors within the data set, and the overall size of the SNP data set). With smaller data sets and data sets with higher levels of ILS, moderate minor allele filters (i.e., MAC=3–4) likely improve topological inferences, but higher minor allele filters (i.e., MAC *>*5) will then reduce the accuracy of inference. However, in trees with low levels of ILS, little topological accuracy is gained with shifting from low (0 ⩽ MAC ⩽ 2) to moderate (MAC=3–4) minor allele filters. When using an outgroup reference, stringent MAC filters will have a stronger effect in reducing the accuracy of inferences, while those same stringent MAC filters may result in more accurate trees when using an ingroup reference.

It is also worth noting that our simulations only included trees simulated under an equal-rates Yule model (with equal speciation rates and zero extinction across all lineages), resulting in balanced topologies with a true *γ ≈* 0. In clades with true divergence histories that are more pectinate or have a higher or lower center of gravity, it is possible that differences in the distribution of SNPs across the tree may cause response patterns to differ. Similarly, imbalanced taxon sampling across the topology (e.g., clustered sampling) may also affect the way that a data set responds to bioinformatic choices, and thus it will be important to investigate these patterns across a variety of different tree shapes to understand our ability to generalize these results. While we did not explicitly test uneven taxon sampling here, an extensive body of work already exists on biases introduced via taxon sampling choices (e.g., Zwickl and Hillis, 2002; Heath et al., 2008b; Hohna et al., 2011; Cusimano and Renner, 2010; Brock et al., 2011), and our results indicate that the choice of reference genome may interact with taxon sampling schemes to bias phylogenetic inferences and diversification statistics. It is also possible that linkage among SNPs—or filtering to reduce linkage—may affect phylogenetic inferences in empirical datasets, although testing this effect was beyond the scope of our simulations.

### Conclusions and Recommendations

Our results suggest that stringent bioinformatic filters tend to reduce the accuracy of phylogenetic analyses and should therefore be used with caution. However, best practices for filtering will depend on the specifics of the divergence history of a clade and the reference genome(s) available for short read alignment. In particular, the rapidity of diversification within a clade is predictive of the sensitivity of phylogenetic analyses to biases introduced by minor allele and missing data filters, such that trees with greater gene tree incongruence require these filters to improve inferences, but also show the most bias compared to the true topologies and true diversification statistics as filter stringency increases. While the patterns differed slightly depending on the phylogenetic inference method used, the biases observed are not completely mitigated by using ILS-aware species tree methods in place of concatenation-based inferences, and in fact may be exacerbated when using species tree methods for clades with low gene tree incongruence. In addition, we find that data aligned to more distant reference genomes are more sensitive to stringent filtering parameters. We further demonstrate that minor allele filters greater than *∼*4% reduce topological accuracy in our 51-taxon phylogenies and increase both imbalance and *γ* statistics, suggesting that they are removing important signal above this threshold rather than only removing sequencing or alignment errors. Thus, our analyses support other recommendations suggesting that filtering for a minor allele count of 3 allows for removing true sequencing errors without removing true signal in the data in most situations (e.g., Rochette et al., 2019; Rivera-Colón and Catchen, 2022; O’Leary et al., 2018). The use of minor allele filters of 5% or higher is common in many empirical studies; these high thresholds may be inadvertently biasing downstream results in these studies.

Intriguingly, we also demonstrate that the best filtering scheme for producing an accurate topology and imbalance statistic may differ from the optimal scheme for accurate *γ* inference. While it may seem counter-intuitive, topology, imbalance, and *γ* can all vary independently (Fig. S30-S32), and thus the optimal filtering scheme likely differs depending on the intended downstream inferences. Although these general patterns are clear from our simulation results, we recommend that researchers still replicate their own analyses over a series of bioinformatic parameter combinations to ensure that results are in agreement across a range of bioinformatic choices. While our results make clear that reference genome choice and bioinformatic filters can bias phylogenetic inferences and downstream macroevolutionary interpretations, the increasing availability of reference-quality genomes makes it possible to implement multiple alignment and filtering schemes to assess the effect that each choice has for the research question of interest. We encourage careful assessment of these choices in empirical data sets.

## Funding

This work was supported by a National Science Foundation grant to C.E.W. (grant number DEB-1556963); by a Graduate Student Research Award from the Society of Systematic Biologists to J.A.R.; by the Aven Nelson Fellowship in Systematic Botany from the University of Wyoming to J.A.R.; by a University of Wyoming NASA Space Grant Research Fellowship from the Wyoming NASA Space Grant Consortium to J.A.R. (NASA Grant number NNX15AI08H); by financial support provided by Robert B. Berry and the University of Wyoming to C.D.B.; and by the University of Wyoming INBRE Data Science Core, which is supported by an Institutional Development Award (IDeA) from the National Institute of General Medical Sciences of the National Institutes of Health (grant number 2P20GM103432).

## Acknowledgements

The authors would like to thank E.J. McTavish for the use of and help with the TreeToReads pipeline. We also thank C.A. Buerkle for insightful discussions about the role of minor alleles and rare variants in evolutionary biology. We also thank members of the Wagner Lab at the University of Wyoming for many discussions related to this work that shaped the direction of this research, and two anonymous reviewers and the associate editor for their thoughtful and insightful comments that greatly improved this manuscript. This work was made possible through the use of the University of Wyoming Advanced Research Computing Center’s Mt. Moran (http://n2t.net/ark:/85786/m4159c), Teton (https://doi.org/10.15786/M2FY47), and Beartooth (https://doi.org/10.15786/m2fy47) computing environments. The findings of this paper are solely the responsibility of the authors and do not necessarily represent the official views of NIH, NSF, or other funding agencies.

## Data Availability Statement

Supplementary material, including data files and online-only appendices, can be found in the Dryad data repository (https://doi.org/10.5061/dryad.djh9w0w2g). Simulated data, simulation scripts, and analysis scripts are available from the Zenodo Repository: http://dx.doi.org/10.5281/zenodo.5940691. Scripts used in simulations and analyses are also available on GitHub https://github.com/jessicarick/refbias_scripts.

## Appendix

### Supplementary Material

#### Supplemental methods

##### Reference genome for simulations

To create a reference genome to use as a base for our `TreeToReads` simulations, we started with the *Lates calcarifer* reference genome (Vij et al., 2016), as this was the most complete of the four reference genomes used in our empirical analyses. We randomly sampled 5,000bp windows without replacement using a custom bash script. For each random segment, we replaced any ambiguous bases (N) with a random base (A, C, G, or T), with equal weighting given to each option.

##### Genotyping-by-sequencing for Lake Tanganyika tropheine cichlids

We sampled individuals in the cichlid tribe Tropheini in the Kigoma region of Tanzania in 2002, 2005, 2007, 2012, 2015, 2016, 2017, and 2018. All sampling was conducted under Institutional Animal Care and Use Committee (IACUC) approved protocols. Briefly, we collected individuals using gill nets while snorkeling in the rocky littoral zone at a 1–10 m depth. We took a fin clip from each individual for genetic analyses, which was stored in either DMSO-EDTA or 95% ethanol prior to DNA extraction. We retained all individuals as vouchers, which are deposited at the Cornell University Museum of Vertebrates (2002, 2005, and 2007 individuals) and the University of Wyoming Museum of Vertebrates (2015, 2016, 2017, and 2018 individuals).

We extracted DNA from the fin clips using DNeasy Blood & Tissue kits (Qiagen, Inc.) and prepared genomic libraries for high-throughput DNA sequencing following the GBS protocol described in Parchman et al. (2012). Briefly, we fragmented DNA using EcoRI and MseI restriction enzymes and then ligated a unique 8 – 10 base pair (bp) barcode to each individual’s DNA. We used polymerase chain reaction (PCR) to amplify the restriction/ligation products (two independent replicates per individual) and then combined the PCR products to create the final libraries. Prior to sequencing, the libraries were size-selected for 250 – 400 bp fragments using BluePippin (Sage Science). Sequencing was completed on Illumina HiSeq 2500 and 4000 platforms (100 bp single-end) at the University of Texas Genome Sequencing and Analysis Facility (Austin, Texas, USA) and the University of Oregon Genomics and Cell Characterization Core Facility (Eugene, Oregon, USA). The individuals in this project were included in libraries containing samples of other cichlid species as part of a larger sequencing effort. Each library contained *∼*100 individuals and was sequenced in its own lane.

With the raw sequence data, we first assigned reads to individuals and subsequently removed the barcode sequences using a custom `perl` script. We then aligned the reads to the *Pundamilia nyererei* and *Oreochromis niloticus* reference genomes (Brawand et al., 2015) using `bwa` v0.7.17 (v0.7.17 Li and Durbin, 2009) with default settings to produce `bam` alignment files, which were used for variant calling in downstream analyses.

#### Supplemental tables and figures

**Table S1.** Model coefficients from linear mixed models, including parameter estimates (Est), minimum 95% confidence interval (Min CI), upper 95% confidence interval (Max CI), and significance at *α* = 0.05 (* indicates *p <*0.05). Parameters are listed for models for all `RAxML` trees, as well as the three ILS levels individually. Results are summarized graphically in Fig. 3). Predictor variables in each model are average distance from the ingroup taxa to the reference genome (avg dxy), minor allele count threshold (maf), missing data threshold (missing), and the interaction among these terms, with simulation number as a random effect.

| Variable | All Trees |  |  |  | Low ILS |  |  |  | Medium ILS |  |  |  | High ILS |  |  |  |
| --- | --- | --- | --- | --- | --- | --- | --- | --- | --- | --- | --- | --- | --- | --- | --- | --- |
|  | Est | Min CI | Max CI | Sig | Est | Min CI | Max CI | Sig | Est | Min CI | Max CI | Sig | Est | Min CI | Max CI | Sig |
| **Standardized ingroup gamma** |  |  |  |  |  |  |  |  |  |  |  |  |  |  |  |  |
| avg_dxy | 1.463 | 0.217 | 2.711 | * | 0.268 | -0.647 | 1.182 |  | 0.203 | -1.169 | 1.578 |  | 0.030 | -1.238 | 1.300 |  |
| maf | 0.815 | 0.760 | 0.870 | * | 0.611 | 0.566 | 0.656 | * | 0.937 | 0.880 | 0.995 | * | 0.851 | 0.799 | 0.903 | * |
| missing | -0.371 | -0.888 | 0.146 |  | -0.282 | -0.704 | 0.140 |  | -0.451 | -0.987 | 0.086 |  | -0.378 | -0.867 | 0.111 |  |
| avg_dxy:maf | -0.089 | -0.280 | 0.102 |  | -0.033 | -0.173 | 0.108 |  | -0.013 | -0.223 | 0.197 |  | -0.013 | -0.208 | 0.181 |  |
| avg_dxy:missing | -0.005 | -1.800 | 1.791 |  | -0.170 | -1.486 | 1.146 |  | 0.006 | -1.969 | 1.980 |  | 0.172 | -1.653 | 1.996 |  |
| **Robinson-Foulds distance to true tree** |  |  |  |  |  |  |  |  |  |  |  |  |  |  |  |  |
| avg_dxy | -0.015 | -0.035 | 0.005 |  | 0.005 | -0.017 | 0.026 |  | 0.003 | -0.024 | 0.029 |  | -0.025 | -0.061 | 0.012 |  |
| maf | 0.013 | 0.012 | 0.014 | * | 0.016 | 0.015 | 0.017 | * | 0.012 | 0.011 | 0.013 | * | 0.012 | 0.010 | 0.013 | * |
| missing | 0.002 | -0.007 | 0.010 |  | 0.001 | -0.009 | 0.010 |  | 0.002 | -0.009 | 0.012 |  | 0.001 | -0.013 | 0.015 |  |
| avg_dxy:maf | 0.003 | 0.000 | 0.006 | * | 0.000 | -0.003 | 0.003 |  | 0.002 | -0.002 | 0.006 |  | 0.007 | 0.002 | 0.013 | * |
| avg_dxy:missing | -0.006 | -0.035 | 0.023 |  | -0.012 | -0.043 | 0.018 |  | -0.012 | -0.050 | 0.027 |  | 0.014 | -0.039 | 0.066 |  |
| **Standardized ingroup Colless imbalance** |  |  |  |  |  |  |  |  |  |  |  |  |  |  |  |  |
| avg_dxy | 0.019 | 0.004 | 0.034 | * | 0.006 | -0.008 | 0.021 |  | 0.010 | -0.019 | 0.038 |  | 0.032 | 0.005 | 0.059 | * |
| maf | 0.010 | 0.009 | 0.010 | * | 0.007 | 0.006 | 0.008 | * | 0.009 | 0.008 | 0.011 | * | 0.012 | 0.011 | 0.013 | * |
| missing | -0.004 | -0.011 | 0.002 |  | -0.001 | -0.008 | 0.006 |  | -0.005 | -0.016 | 0.006 |  | -0.007 | -0.018 | 0.003 |  |
| avg_dxy:maf | -0.002 | -0.005 | 0.000 | * | 0.000 | -0.002 | 0.002 |  | -0.001 | -0.005 | 0.004 |  | -0.005 | -0.009 | -0.001 | * |
| avg_dxy:missing | -0.001 | -0.023 | 0.020 |  | -0.006 | -0.028 | 0.015 |  | 0.002 | -0.039 | 0.043 |  | 0.000 | -0.039 | 0.039 |  |

**Table S2.** Model coefficients from linear mixed models for subsampled data sets, including parameter estimates (Est), minimum 95% confidence interval (Min CI), upper 95% confidence interval (Max CI), and significance at *α* = 0.05 (* indicates *p <*0.05). Parameters are listed for models for all trees, as well as the three ILS levels individually. Results are summarized graphically in Fig. 5). Predictor variables in each model are average distance from the ingroup taxa to the reference genome (avg dxy), minor allele count threshold (maf), missing data threshold (missing), and the interaction among these terms, with simulation number as a random effect.

| Variable | All Trees |  |  |  | Low ILS |  |  |  | Medium ILS |  |  |  | High ILS |  |  |  |
| --- | --- | --- | --- | --- | --- | --- | --- | --- | --- | --- | --- | --- | --- | --- | --- | --- |
|  | Est | Min CI | Max CI | Sig | Est | Min CI | Max CI | Sig | Est | Min CI | Max CI | Sig | Est | Min CI | Max CI | Sig |
| **Standardized ingroup gamma** |  |  |  |  |  |  |  |  |  |  |  |  |  |  |  |  |
| avg.dxy | 0.729 | 0.236 | 1.222 | * | 0.029 | -0.435 | 0.493 |  | 0.313 | -0.425 | 1.051 |  | -0.074 | -0.699 | 0.552 |  |
| maf | 0.755 | 0.739 | 0.771 | * | 0.254 | 0.238 | 0.271 | * | 0.931 | 0.908 | 0.953 | * | 0.862 | 0.843 | 0.881 | * |
| missing | -0.449 | -0.645 | -0.253 | * | -0.034 | -0.238 | 0.171 |  | -0.590 | -0.872 | -0.309 | * | -0.544 | -0.780 | -0.308 | * |
| avg.dxy:maf | -0.127 | -0.183 | -0.071 | * | 0.021 | -0.032 | 0.073 |  | -0.075 | -0.158 | 0.009 |  | 0.024 | -0.047 | 0.095 |  |
| avg.dxy:missing | 0.115 | -0.579 | 0.809 |  | -0.020 | -0.673 | 0.632 |  | 0.036 | -1.001 | 1.074 |  | 0.044 | -0.836 | 0.924 |  |
| **Robinson-Foulds distance to true tree** |  |  |  |  |  |  |  |  |  |  |  |  |  |  |  |  |
| avg.dxy | -0.014 | -0.030 | 0.001 |  | -0.012 | -0.030 | 0.007 |  | 0.016 | -0.006 | 0.039 |  | -0.005 | -0.031 | 0.020 |  |
| maf | 0.006 | 0.006 | 0.007 | * | 0.015 | 0.014 | 0.016 | * | 0.005 | 0.005 | 0.006 | * | 0.002 | 0.002 | 0.003 | * |
| missing | -0.005 | -0.011 | 0.002 |  | -0.003 | -0.012 | 0.005 |  | -0.001 | -0.010 | 0.007 |  | -0.009 | -0.018 | 0.001 |  |
| avg.dxy:maf | 0.003 | 0.001 | 0.005 | * | 0.001 | -0.001 | 0.003 |  | 0.000 | -0.002 | 0.003 |  | 0.002 | -0.001 | 0.005 |  |
| avg.dxy:missing | 0.003 | -0.019 | 0.025 |  | 0.005 | -0.022 | 0.031 |  | -0.011 | -0.043 | 0.020 |  | 0.016 | -0.021 | 0.052 |  |
| **Standardized ingroup Colless imbalance** |  |  |  |  |  |  |  |  |  |  |  |  |  |  |  |  |
| avg.dxy | -0.001 | -0.011 | 0.009 |  | 0.007 | -0.005 | 0.020 |  | -0.010 | -0.028 | 0.008 |  | -0.001 | -0.019 | 0.016 |  |
| maf | 0.009 | 0.009 | 0.010 | * | 0.009 | 0.009 | 0.009 | * | 0.010 | 0.010 | 0.011 | * | 0.009 | 0.008 | 0.009 | * |
| missing | -0.005 | -0.009 | -0.001 | * | 0.002 | -0.004 | 0.007 |  | -0.007 | -0.014 | 0.000 |  | -0.008 | -0.015 | -0.002 | * |
| avg.dxy:maf | 0.002 | 0.001 | 0.003 | * | 0.000 | -0.001 | 0.002 |  | 0.004 | 0.002 | 0.006 | * | 0.001 | -0.001 | 0.003 |  |
| avg.dxy:missing | -0.005 | -0.019 | 0.010 |  | -0.012 | -0.030 | 0.006 |  | -0.008 | -0.034 | 0.017 |  | 0.002 | -0.023 | 0.027 |  |

**Table S3.** Model coefficients from linear mixed models for the empirical Tropheine and *Lates* data sets, including parameter estimates (Est), minimum 95% confidence interval (Min CI), upper 95% confidence interval (Max CI), and significance at *α* = 0.05. Parameters are listed for models for all trees, as well as the three ILS levels individually. Results are summarized graphically in Fig. 8). Predictor variables in each model are reference genome choice (ref), minor allele count threshold (mac), missing data threshold (missing), and the interaction among these terms, with iteration number as a random effect. Reference genome choice (ref) was coded as 0=outgroup reference and 1=ingroup reference.

| Variable | Tropheine Topologies |  |  |  | Lates Topologies |  |  |  |
| --- | --- | --- | --- | --- | --- | --- | --- | --- |
|  | Est | Min CI | Max CI | Sig | Est | Min CI | Max CI | Sig |
| **Ingroup gamma** |  |  |  |  |  |  |  |  |
| ref | 0.111 | -0.037 | 0.259 |  | 0.030 | -0.390 | 0.450 |  |
| mac | 0.262 | 0.246 | 0.278 | * | -0.254 | -0.299 | -0.208 | * |
| missing | -1.204 | -1.357 | -1.051 | * | -0.345 | -0.773 | 0.082 |  |
| ref:mac | -0.025 | -0.048 | -0.002 | * | 0.021 | -0.043 | 0.086 |  |
| ref:missing | 0.420 | 0.207 | 0.634 | * | 0.161 | -0.444 | 0.766 |  |
| **Ingroup tree height** |  |  |  |  |  |  |  |  |
| ref | 0.013 | -0.007 | 0.034 |  | -0.006 | -0.034 | 0.022 |  |
| mac | 0.022 | 0.019 | 0.024 | * | -0.032 | -0.035 | -0.029 | * |
| missing | -0.018 | -0.039 | 0.004 |  | -0.062 | -0.091 | -0.033 | * |
| ref:mac | -0.020 | -0.024 | -0.017 | * | 0.000 | -0.005 | 0.004 |  |
| ref:missing | 0.032 | 0.002 | 0.062 | * | -0.008 | -0.049 | 0.032 |  |
| **Ingroup Colless imbalance** |  |  |  |  |  |  |  |  |
| ref | 0.020 | -0.025 | 0.064 |  | -0.011 | -0.031 | 0.009 |  |
| mac | -0.011 | -0.016 | -0.006 | * | -0.001 | -0.003 | 0.001 |  |
| missing | -0.224 | -0.270 | -0.178 | * | -0.009 | -0.029 | 0.012 |  |
| ref:mac | 0.002 | -0.005 | 0.009 |  | 0.002 | -0.001 | 0.005 |  |
| ref:missing | 0.006 | -0.059 | 0.070 |  | 0.002 | -0.027 | 0.031 |  |

**Table S4.** Model coefficients from linear mixed models for `ASTRAL` trees, including parameter estimates (Est), minimum 95% confidence interval (Min CI), upper 95% confidence interval (Max CI), and significance at *α* = 0.05 (* indicates *p <*0.05). Parameters are listed for models for all trees, as well as the three ILS levels individually. Predictor variables in each model are average distance from the ingroup taxa to the reference genome (avg dxy), minor allele count threshold (mac), missing data threshold (missing), and the interaction among these terms, with simulation number as a random effect.

| Variable | All Trees |  |  |  | Low ILS |  |  |  | Medium ILS |  |  |  | High ILS |  |  |  |
| --- | --- | --- | --- | --- | --- | --- | --- | --- | --- | --- | --- | --- | --- | --- | --- | --- |
|  | Est | Min CI | Max CI | Sig | Est | Min CI | Max CI | Sig | Est | Min CI | Max CI | Sig | Est | Min CI | Max CI | Sig |
| **Robinson-Foulds distance to true tree** |  |  |  |  |  |  |  |  |  |  |  |  |  |  |  |  |
| avg_dxy | -0.007 | -0.022 | 0.009 |  | -0.011 | -0.036 | 0.015 |  | -0.001 | -0.022 | 0.020 |  | 0.001 | -0.018 | 0.019 |  |
| maf | 0.000 | -0.001 | 0.000 |  | 0.000 | -0.001 | 0.002 |  | 0.000 | -0.001 | 0.001 |  | -0.001 | -0.002 | 0.000 | * |
| missing | -0.008 | -0.014 | -0.002 | * | -0.015 | -0.027 | -0.003 | * | -0.006 | -0.015 | 0.002 |  | -0.002 | -0.009 | 0.005 |  |
| avg_dxy:maf | 0.001 | -0.002 | 0.003 |  | 0.000 | -0.004 | 0.003 |  | -0.001 | -0.004 | 0.002 |  | 0.004 | 0.001 | 0.007 | * |
| avg_dxy:missing | -0.002 | -0.024 | 0.020 |  | 0.018 | -0.019 | 0.054 |  | 0.000 | -0.031 | 0.030 |  | -0.031 | -0.058 | -0.005 | * |
| **Standardized ingroup Colless imbalance** |  |  |  |  |  |  |  |  |  |  |  |  |  |  |  |  |
| avg_dxy | -0.018 | -0.042 | 0.006 |  | -0.021 | -0.054 | 0.012 |  | 0.003 | -0.028 | 0.035 |  | -0.014 | -0.046 | 0.019 |  |
| maf | 0.019 | 0.018 | 0.020 | * | 0.028 | 0.027 | 0.030 | * | 0.016 | 0.015 | 0.018 | * | 0.014 | 0.013 | 0.016 | * |
| missing | 0.013 | 0.003 | 0.023 | * | 0.016 | 0.001 | 0.031 | * | 0.006 | -0.006 | 0.019 |  | 0.019 | 0.007 | 0.032 | * |
| avg_dxy:maf | 0.007 | 0.003 | 0.011 | * | 0.002 | -0.003 | 0.007 |  | 0.002 | -0.003 | 0.007 |  | 0.009 | 0.005 | 0.014 | * |
| avg_dxy:missing | 0.000 | -0.035 | 0.035 |  | 0.010 | -0.037 | 0.058 |  | -0.004 | -0.049 | 0.041 |  | -0.017 | -0.064 | 0.030 |  |

**Table S5.** Model coefficients from linear mixed models for `ASTRAL` trees for the empirical datasets, including parameter estimates (Est), minimum 95% confidence interval (Min CI), upper 95% confidence interval (Max CI), and significance at *α* = 0.05 (* indicates *p <*0.05). Parameters are listed for models for the Tropheine trees and *Lates* trees. Predictor variables in each model are reference genome choice (ref), minor allele count threshold (mac), missing data threshold (missing), and the interaction among these terms, with iteration number as a random effect. Reference genome choice (ref) was coded as 0=outgroup reference and 1=ingroup reference.

| Variable | Tropheine Topologies |  |  |  | Lates Topologies |  |  |  |
| --- | --- | --- | --- | --- | --- | --- | --- | --- |
|  | Est | Min CI | Max CI | Sig | Est | Min CI | Max CI | Sig |
| **Ingroup Colless imbalance** |  |  |  |  |  |  |  |  |
| intINT | -0.025 | -0.054 | 0.004 |  | -0.051 | -0.077 | -0.026 | * |
| maf | 0.001 | -0.002 | 0.005 |  | 0.015 | 0.012 | 0.017 | * |
| missing | -0.127 | -0.157 | -0.098 | * | -0.048 | -0.074 | -0.022 | * |
| intINT:maf | -0.002 | -0.006 | 0.003 |  | 0.004 | 0.000 | 0.007 |  |
| intINT:missing | -0.007 | -0.049 | 0.035 |  | -0.047 | -0.084 | -0.009 | * |

**Fig. S1.**
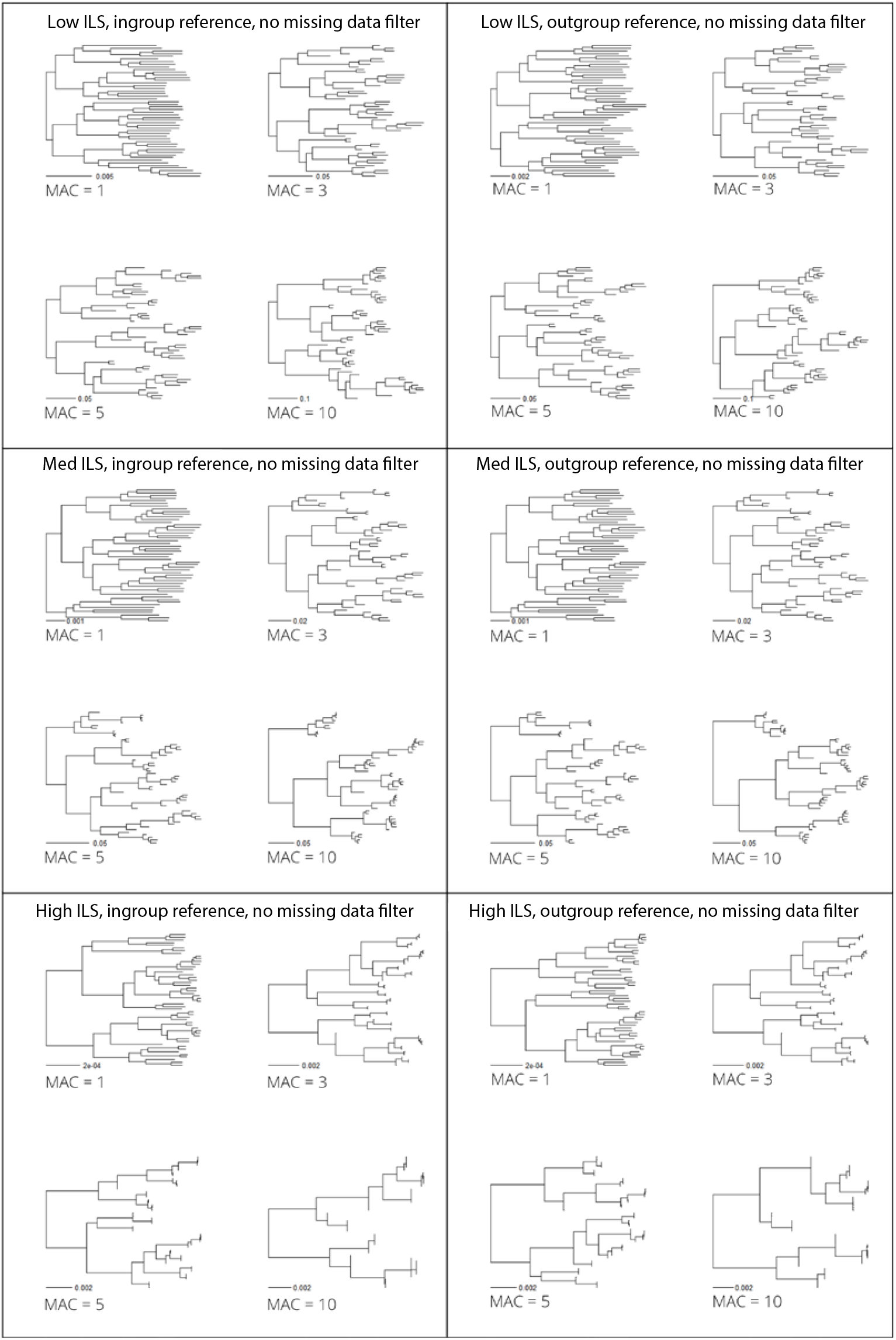
Examples of inferred trees for a single simulation for high, medium, and low ILS species trees, where data were aligned to an ingroup (left) versus outgroup (right) reference genome. Trees are shown for no missing data cutoff and a minimum minor allele count of 1, 3, 5, and 10, to demonstrate the changes in the trees that are summarized through the *γ* and imbalance statistics.

**Fig. S2.**
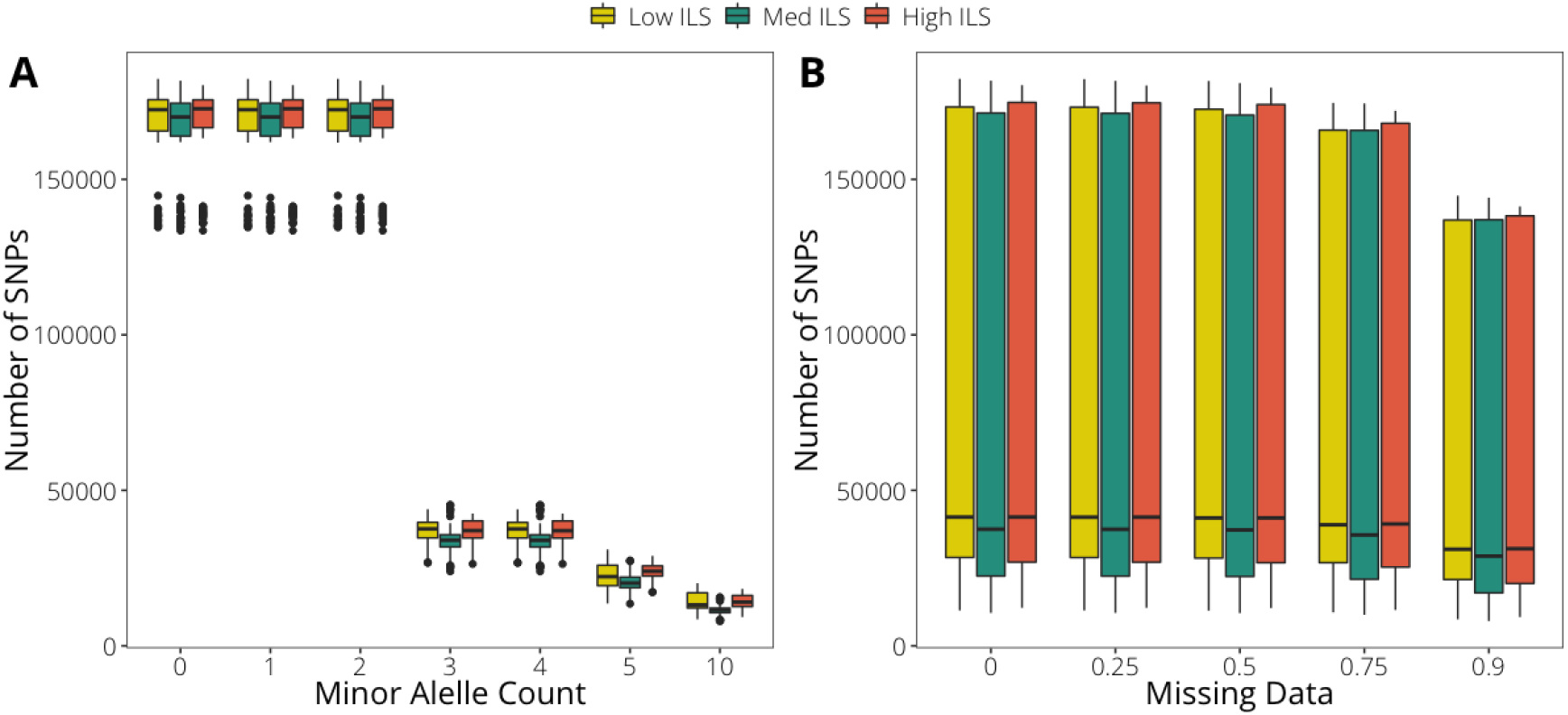
Relationship within the full simulation data sets between (A) minor allele count threshold and the number of sites retained in a data set, and (B) missing data threshold and the number of sites retained in a data set, with colors indicating the tree height.

**Fig. S3.**
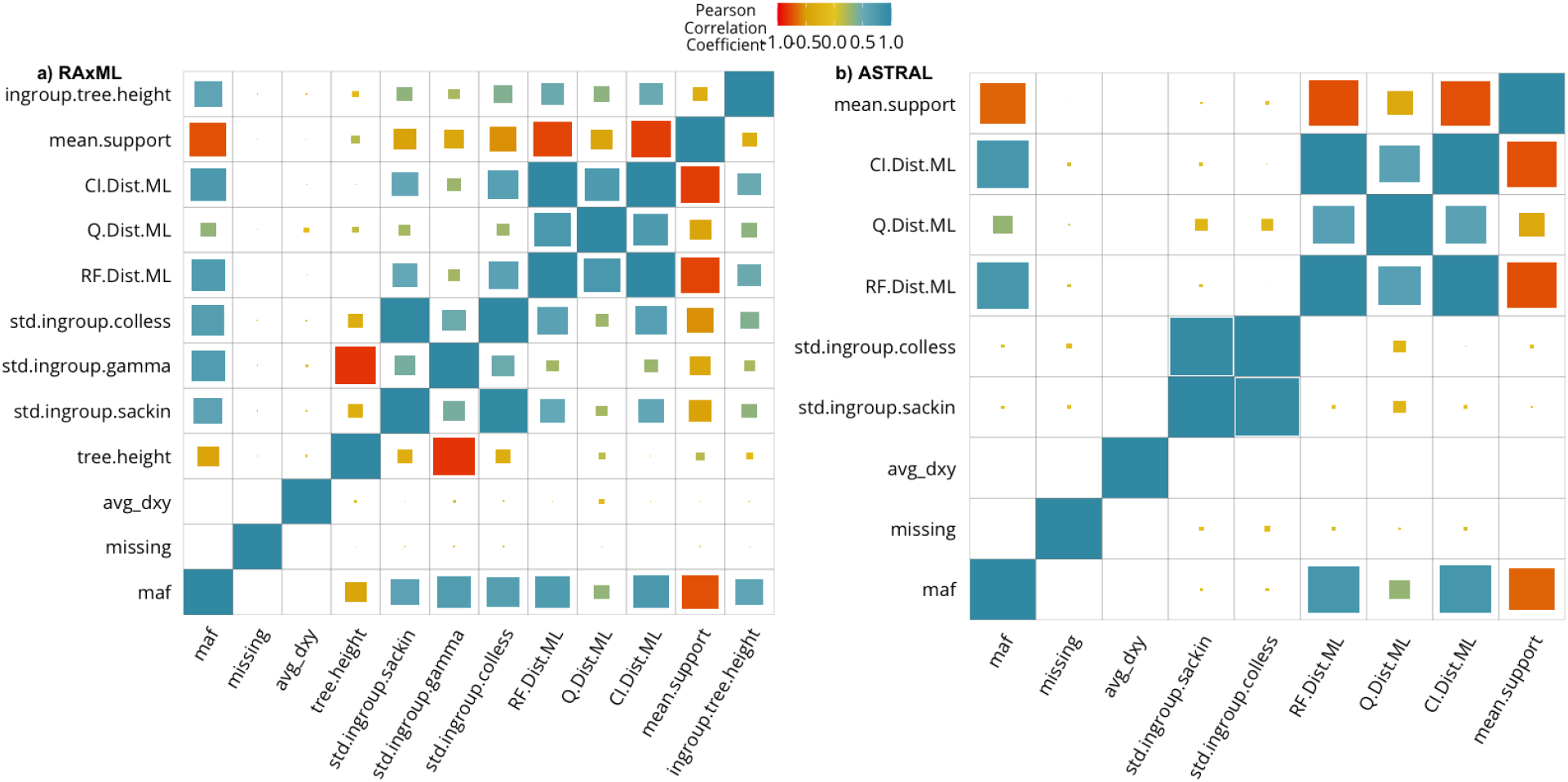
Correlation matrix between output tree characteristics for the full simulation data sets, for phylogenies inferred in `RAxML` (a) and `ASTRAL` (b). Color and size correspond to the magnitude of the Pearson’s correlation between the two parameters. Variables are overall tree height (tree.height), standardized Sackin imbalance of the ingroup (std.ingroup.sackin), standardized *γ* of the ingroup (std.ingroup.gamma), standardized Colless imbalance of the ingroup (std.ingroup.colless), Robinson Foulds distance to the true tree (RF.Dist.ML), quartet distance to the true tree (Q.Dist.ML), mean bootstrap support of all nodes in the tree (mean.support), height of the ingroup (ingroup.tree.height), and clustering information distance to the true tree (CI.Dist.ML). Statistics requiring branch lengths were not calculated for `ASTRAL`-inferred trees.

**Fig. S4.**
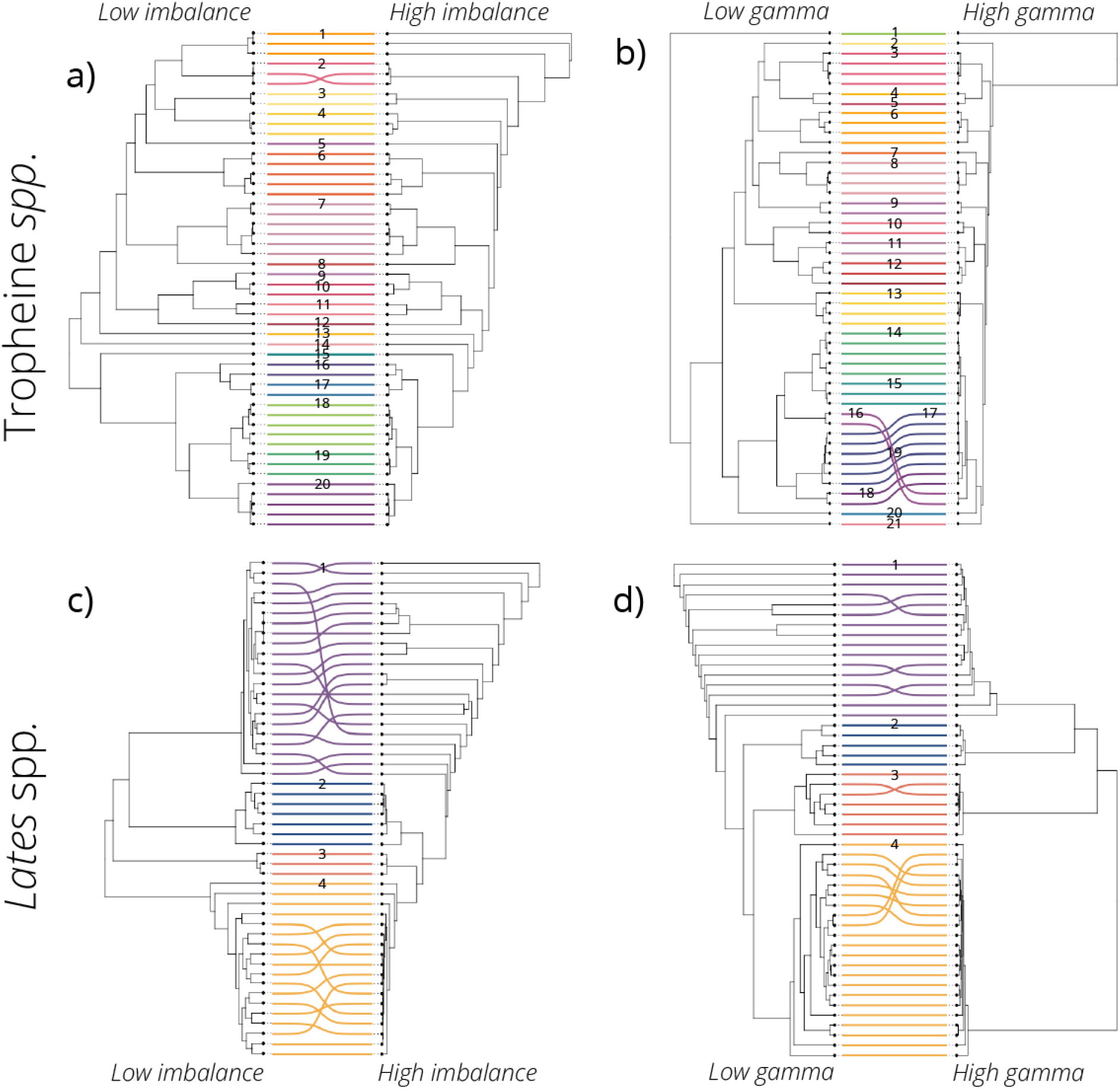
Examples of inferred trees for tropheine (top) and *Lates* datasets, demonstrating extreme values for Colless’ imbalance (left) and gamma (right). In each comparison, the two paired trees have the same individuals and differ only in filtering parameters. Each individual is connected to itself by a line colored according to species identity. For the tropheine phylogenies in (a), species identities are 1: *Tropheus annectens*, 2: *Tropheus kaiser*, 3: *Tropheus sp.*“kirschfleck”, 4: *Tropheus sp.* “crescentic”, 5: *Tropheus brichardi*, 6: *Tropheus duboisi*, 7: *Pundamilia nyererei*, 8: *Lobochilotes labiatus*, 9: *Petrochromis sp.* “green”, 10: *Petrochromis moshi*, 11: *Petrochromis sp.* “kazumbe”, 12: *Petrochromis* cf. *polyodon*, 13: *Petrochromis fasciolatus*, 14: *Petrochromis orthognathus*, 15: *Petrochromis famula*, 16: *Simochromis diagramma*, 17: *Limnotilapia dardennii*, 18: *Ctenochromis horei*, 19: *Pseudosimochromis marginatus*, 20: *Pseudosimochromis babaulti*. For the tropheine phylogenies in (b), species are 1: *Xenotilapia sima*, 2: *Limnotilapia dardennii*, 3: *Pseudosimochromis marginatus*, 4: *Pseudosimochromis babaulti*, 5: *Pseudosimochromis margaritae*, 6: *Petrochromis famula*, 7: *Petrochromis fasciolatus*, 8: *Petrochromis orthognathus*, 9: *Tropheus kaiser*, 10: *Tropheus sp.* “kirschfleck”, 11: *Lobochilotes labiatus*, 12: *Petrochromis moshi*, 13: *Petrochromis sp.* “kazumbe”, 14: *Petrochromis* cf. *polyodon*, 15: *Simochromis diagramma*, 16: *Tropheus annectens*, 17: *Tropheus sp.* “crescentic”, 18: *Tropheus brichardi*, 19: *Tropheus duboisi*, 20: *Pundamilia nyererei*. In both (c) and (d), the *Lates* species identities are 1: *L. mariae*, 2: *L. angustifrons*, 3: *L. microlepis*, 4: *L. stappersii*.

**Fig. S5.**
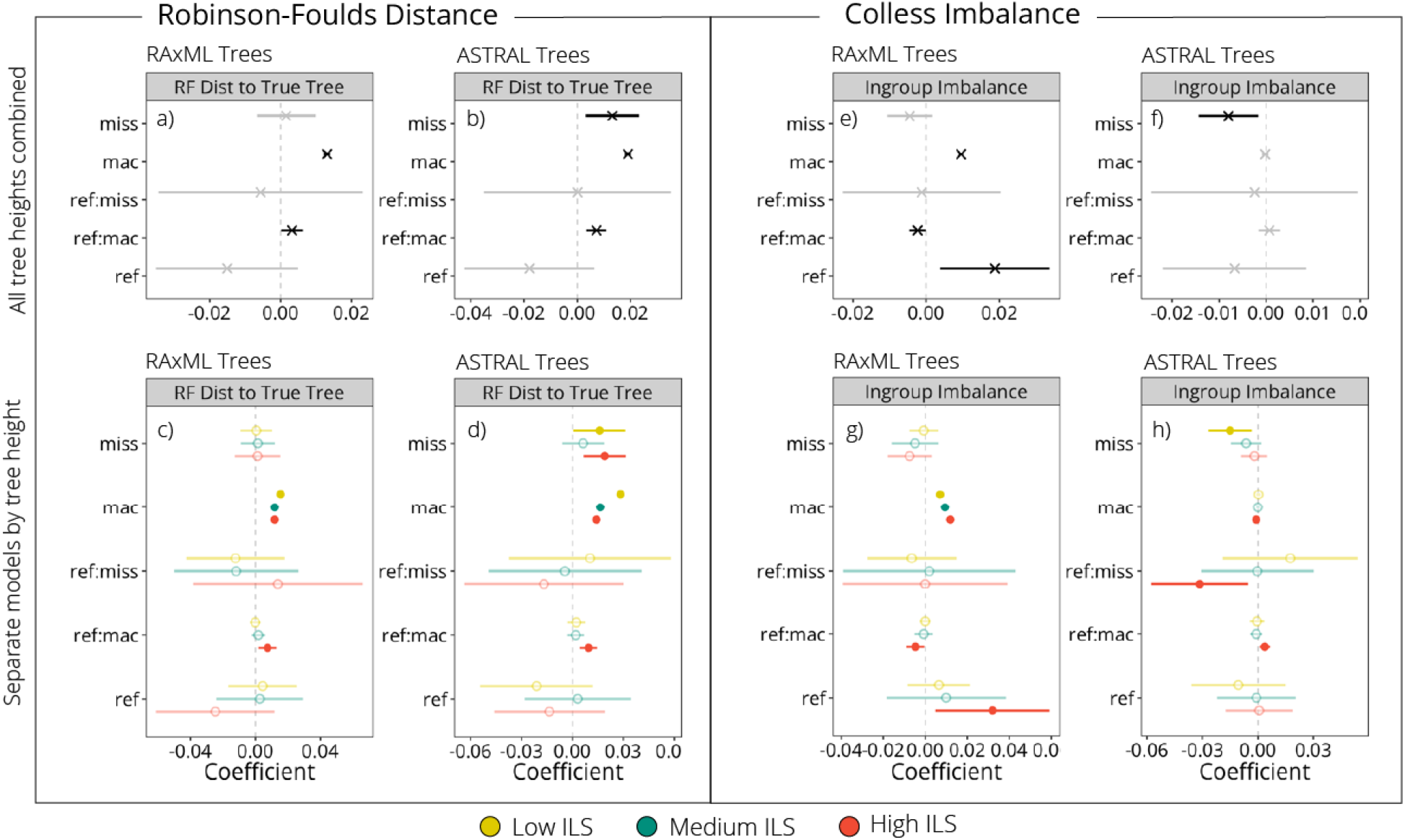
Minor allele count (MAC) threshold had the greatest effect on the Robinson-Foulds distance (RF Dist) from an output tree to the true tree for both `RAxML` and `ASTRAL` output trees, with some interaction between minor allele count threshold and distance to the reference genome and tree height (i.e., amount of incomplete lineage sorting in the true tree). For predicting tree imbalance, MAC threshold was the strongest predictor for `RAxML` trees, while imbalance for `ASTRAL` trees was more dependent on the missing data cutoff. Plots in both (a)–(d) show coefficient estimates and 95% confidence intervals for each predictor variable from linear mixed models with Robinson-Foulds distance to the true tree as the response variable, while (e)–(h) are coefficient estimates and 95% confidence intervals for each predictor variable from linear mixed models with Colless’ imbalance statistic as the response variable. Filled in and darkened symbols indicate significant positive or negative relationships, while faded symbols are predictors with confidence intervals overlapping zero. Plots in (a),(b), (e), and (f) are the results for all tree types combined, while those in (c), (d), (g), and (h) were modeled individually for each of the three simulated ILS levels. For model variables, mac = minor allele count threshold, avg dxy = mean distance from ingroup taxa to the reference genome, miss = maximum missing data allowed, and colons show interaction terms between the given variables.

**Fig. S6.**
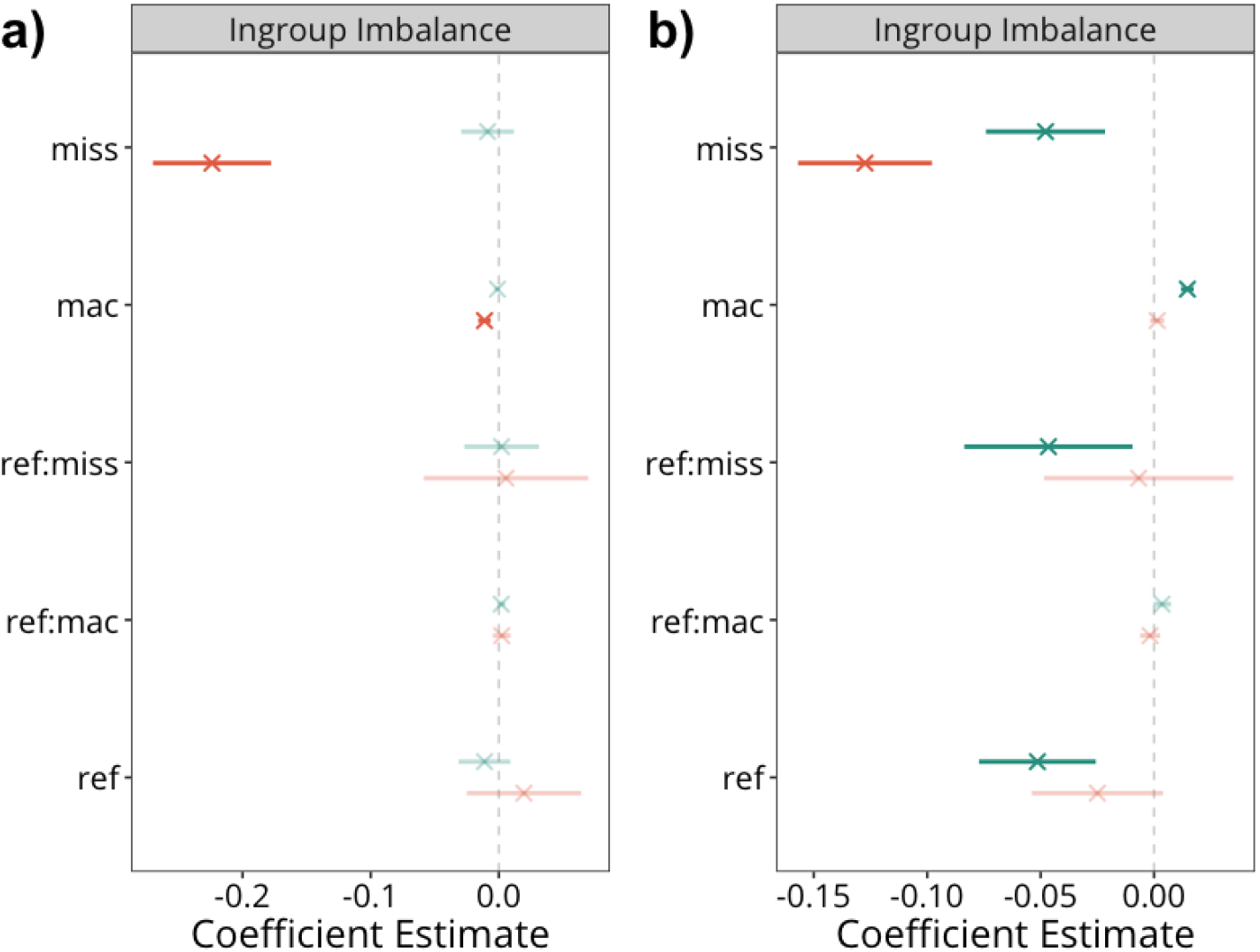
A comparison of model coefficients for trees inferred using (a) `RAxML` and (b) `ASTRAL` for the Tropheine (orange) and *Lates* (teal) datasets. For the Tropheine dataset, missing data continued to be the strongest predictor of imbalance for the `ASTRAL` trees, minor allele count (MAC) threshold was a stronger predictor of imbalance, and the interaction between the reference genome and the missing data cutoff was more important in the `ASTRAL` trees than those inferred using `RAxML`. For the *Lates* dataset, missing data and MAC thresholds were both stronger predictors of imbalance in the `ASTRAL` trees, and the reference genome also had a stronger effect. Plots show coefficient estimates and 95% confidence intervals for each predictor variable from linear mixed models with the Colless’ imbalance statistic of the ingroup as the response variable, with missing data cutoff (miss), minor allele count (mac), reference genome (ref), the interaction between missing data and reference genome choice (int:miss), and the interaction between minor allele count and reference genome choice (int:mac) as explanatory variables. Filled in and darkened symbols indicate significant positive or negative relationships, while faded symbols are predictors with confidence intervals overlapping zero.

**Fig. S7.**
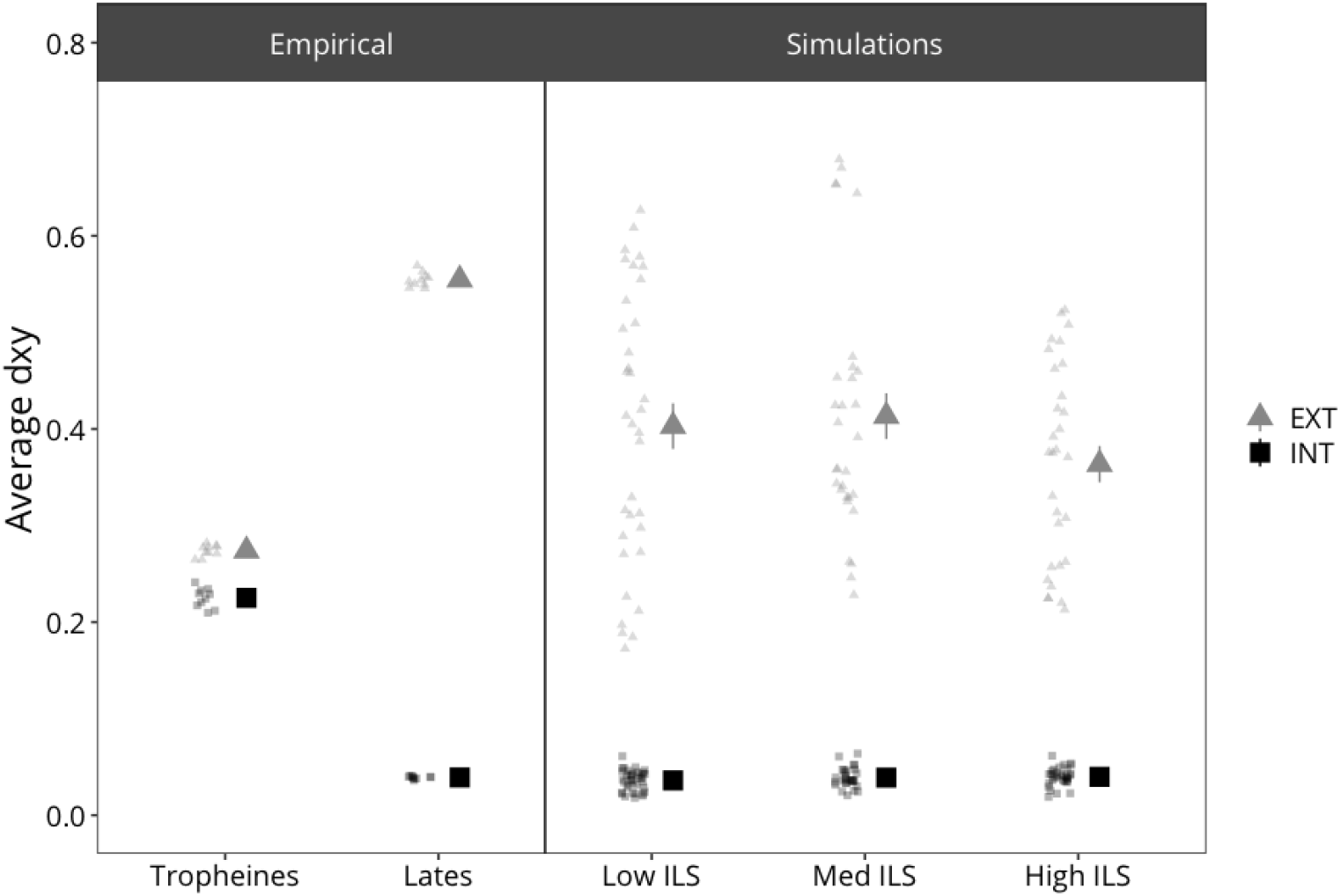
A comparison of the average distance of ingroup taxa to the reference genome for the empirical (left) and simulated (right) datasets. Each light-colored point represents one replicate dataset, and larger dark points show the mean and standard error for each category.

**Fig. S8.**
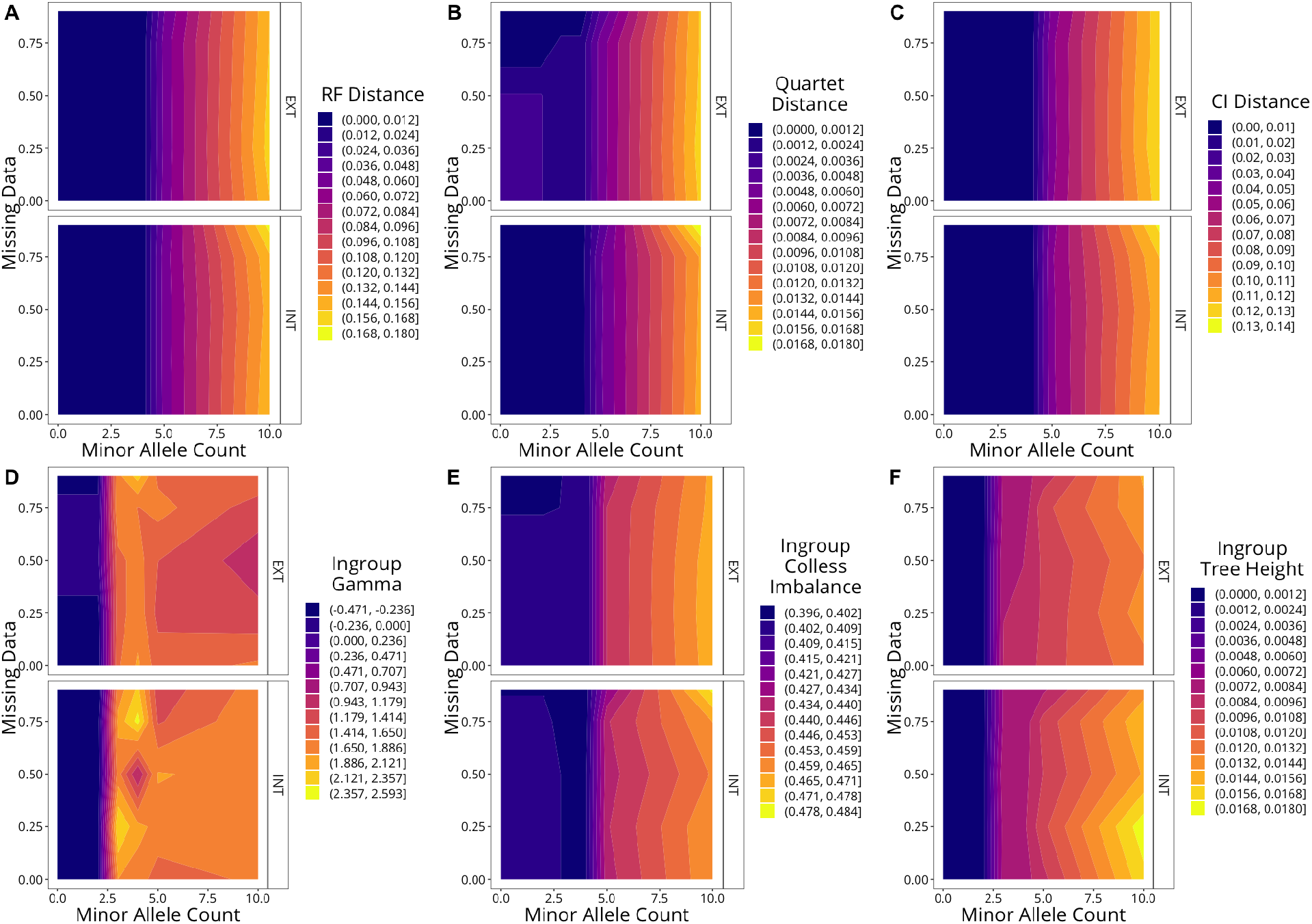
Heatmaps for the **low ILS trees** indicating how each of our tree descriptive parameters varies with minor allele count and missing data thresholds, with darker colors indicating lower values and lighter colors indicating higher values. Three of the parameters, Robinson-Foulds (RF) distance, quartet distance, and clustering information (CI) distance, are measures of topological dissimilarity compared to the true simulated tree, and lower values indicate topologies closer to the true tree. The gamma and Colless imbalance statistics are measures of tree center of gravity and tree shape, with means for the true trees around *I_c_*=0.4 and *γ*=0.

**Fig. S9.**
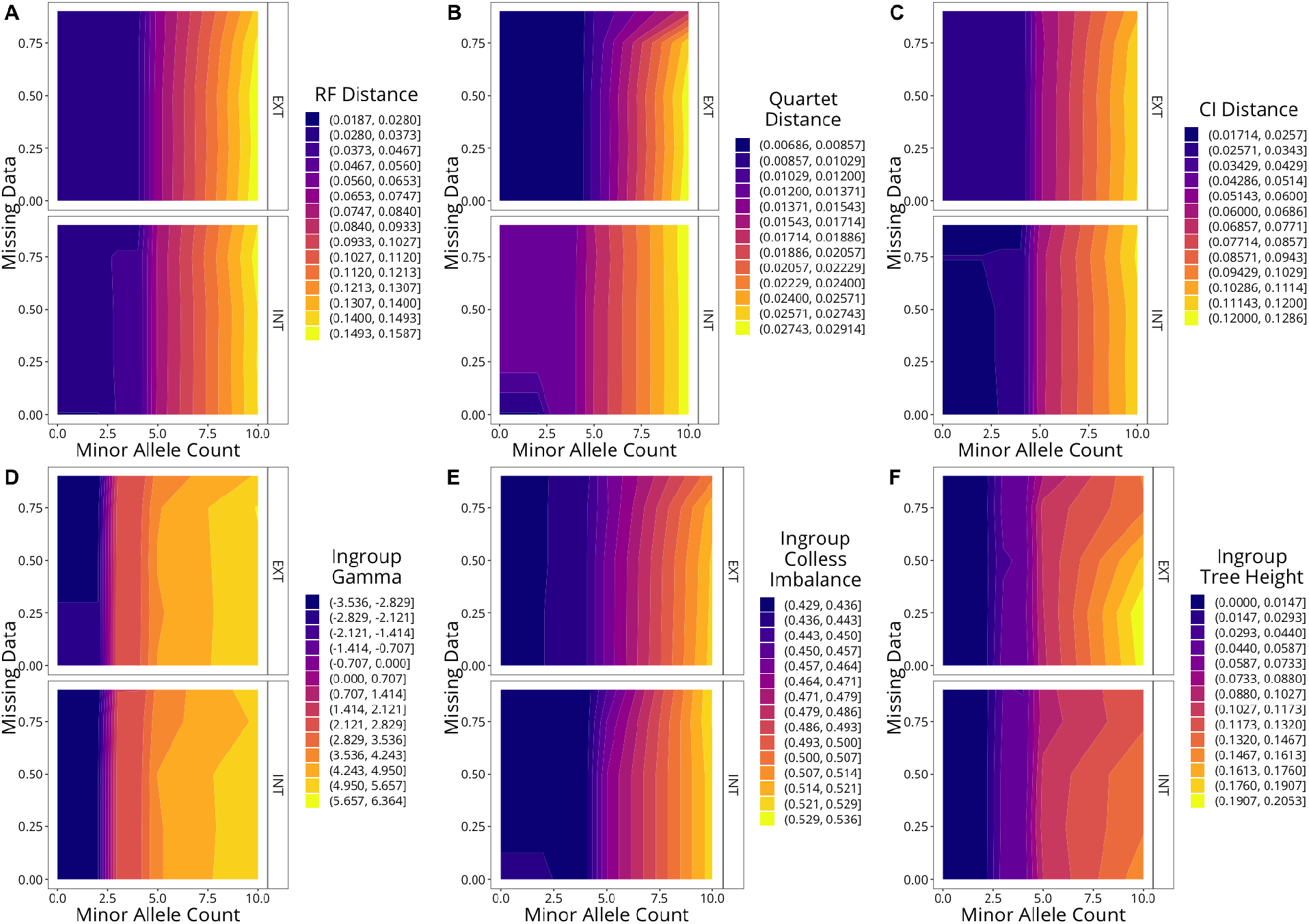
Heatmaps for the **medium ILS trees**indicating how each of our tree descriptive parameters varies in two-dimensional space with minor allele count and missing data thresholds, with darker colors indicating lower values and lighter colors indicating higher values. Three of the parameters, Robinson-Foulds (RF) distance, quartet distance, and clustering information (CI) distance, are measures of topological dissimilarity compared to the true simulated tree, and lower values indicate topologies closer to the true tree. The gamma and Colless imbalance statistics are measures of tree center of gravity and tree shape, with means for the true trees around *I_c_*=0.4 and *γ*=0.

**Fig. S10.**
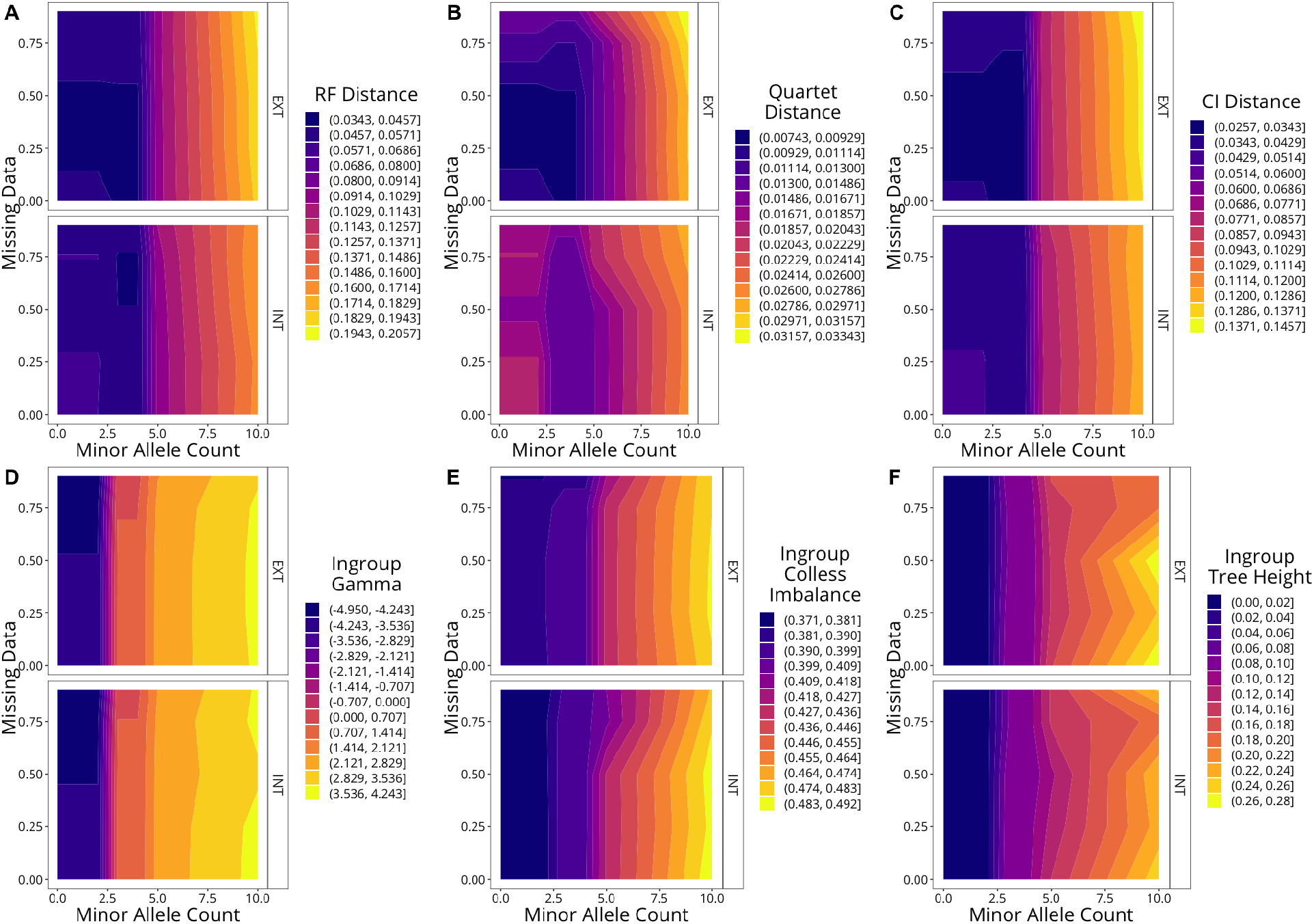
Heatmaps for the **high ILS trees**indicating how each of our tree descriptive parameters varies in two-dimensional space with minor allele count and missing data thresholds, with darker colors indicating lower values and lighter colors indicating higher values. Three of the parameters, Robinson-Foulds (RF) distance, quartet distance, and clustering information (CI) distance, are measures of topological dissimilarity compared to the true simulated tree, and lower values indicate topologies closer to the true tree. The gamma and Colless imbalance statistics are measures of tree center of gravity and tree shape, with means for the true trees around *I_c_*=0.4 and *γ*=0.

**Fig. S11.**
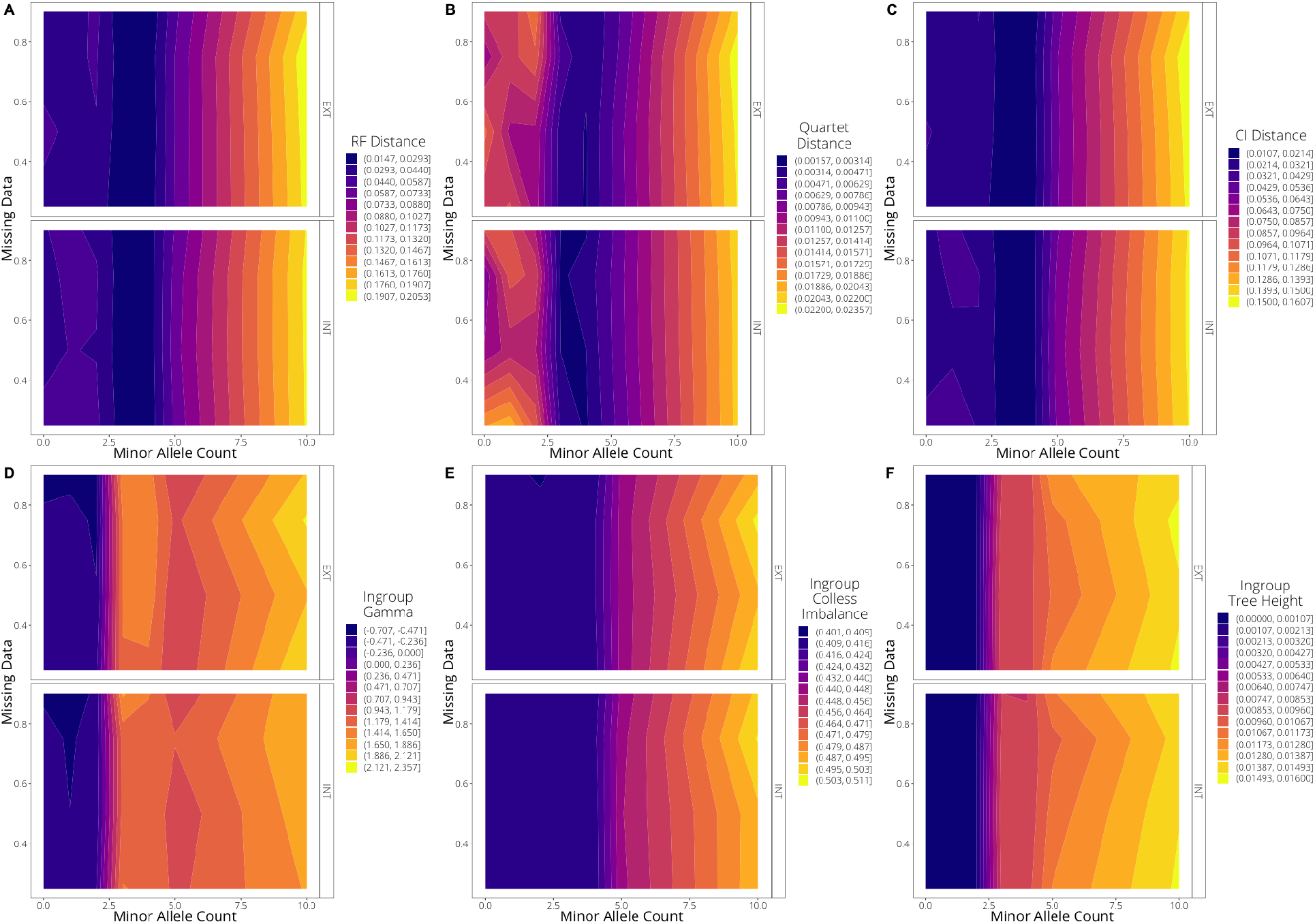
Heatmaps for the **subsampled low ILS trees**indicating how each of our tree descriptive parameters varies with minor allele count and missing data thresholds, with darker colors indicating lower values and lighter colors indicating higher values. Three of the parameters, Robinson-Foulds (RF) distance, quartet distance, and clustering information (CI) distance, are measures of topological dissimilarity compared to the true simulated tree, and lower values indicate topologies closer to the true tree. The gamma and Colless imbalance statistics are measures of tree center of gravity and tree shape, with means for the true trees around *I_c_*=0.4 and *γ*=0.

**Fig. S12.**
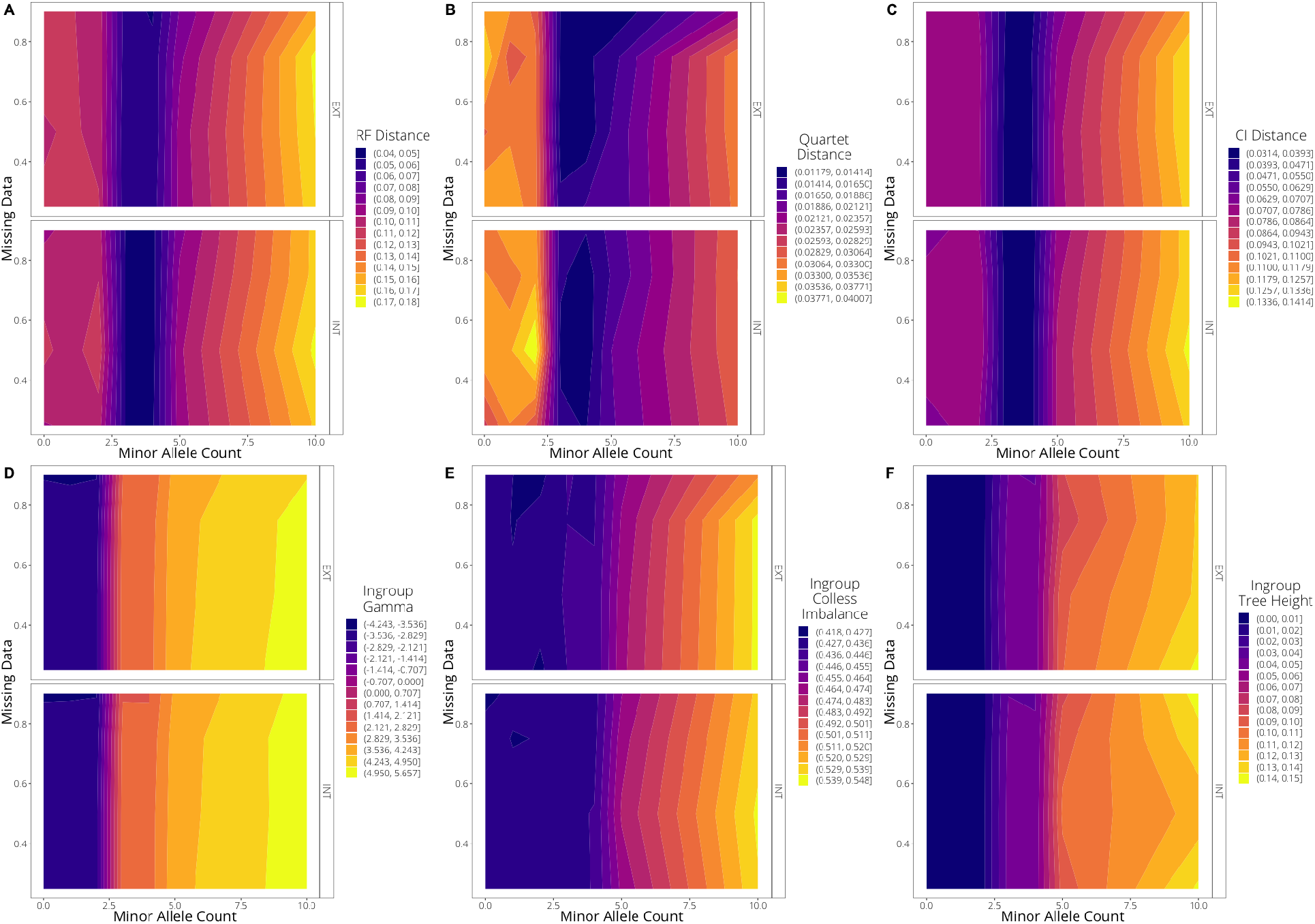
Heatmaps for the **subsampled medium ILS trees** indicating how each of our tree descriptive parameters varies in two-dimensional space with minor allele count and missing data thresholds, with darker colors indicating lower values and lighter colors indicating higher values. Three of the parameters, Robinson-Foulds (RF) distance, quartet distance, and clustering information (CI) distance, are measures of topological dissimilarity compared to the true simulated tree, and lower values indicate topologies closer to the true tree. The gamma and Colless imbalance statistics are measures of tree center of gravity and tree shape, with means for the true trees around *I_c_*=0.4 and *γ*=0.

**Fig. S13.**
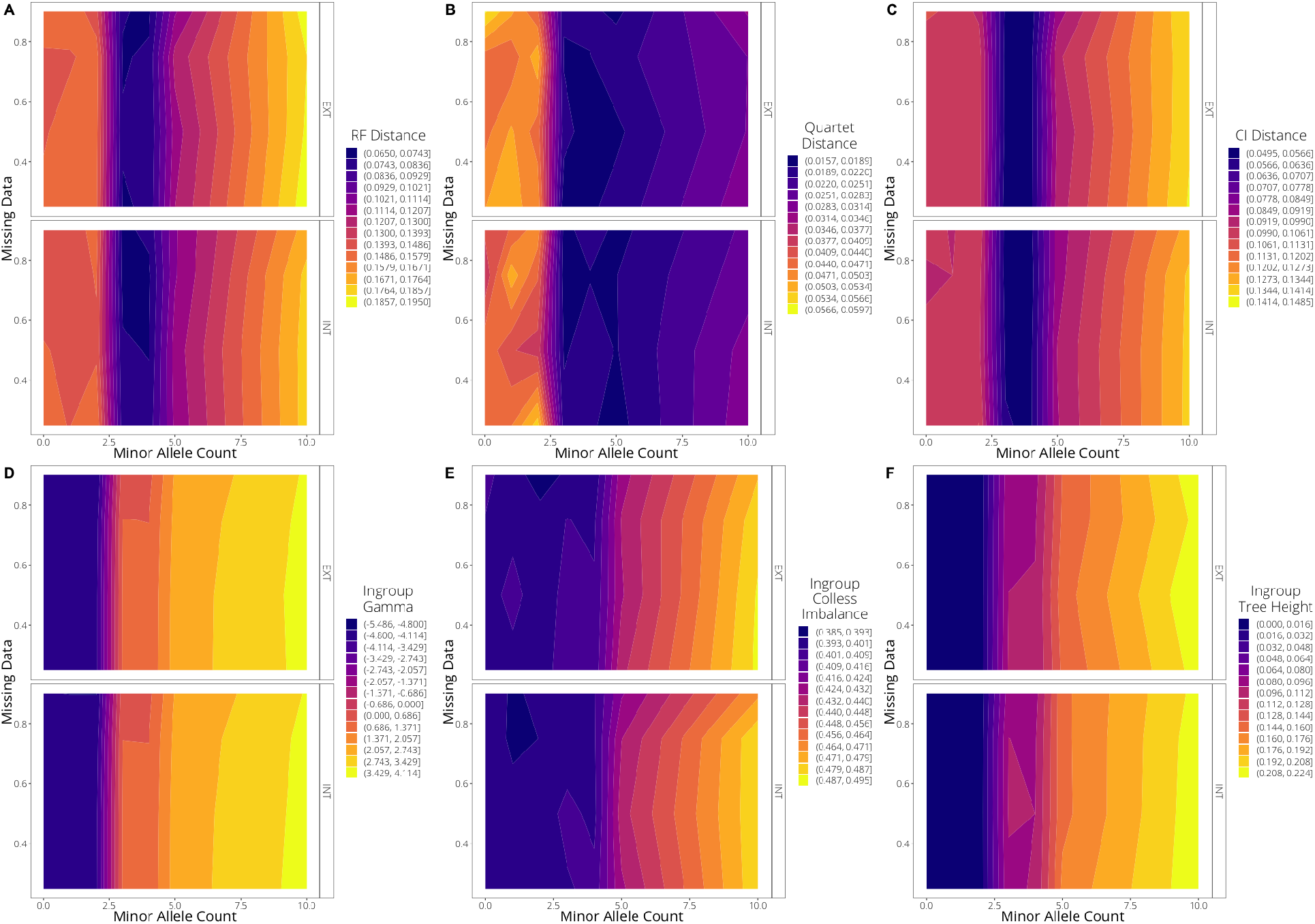
Heatmaps for the **subsampled high ILS trees** indicating how each of our tree descriptive parameters varies in two-dimensional space with minor allele count and missing data thresholds, with darker colors indicating lower values and lighter colors indicating higher values. Three of the parameters, Robinson-Foulds (RF) distance, quartet distance, and clustering information (CI) distance, are measures of topological dissimilarity compared to the true simulated tree, and lower values indicate topologies closer to the true tree. The gamma and Colless imbalance statistics are measures of tree center of gravity and tree shape, with means for the true trees around *I_c_*=0.4 and *γ*=0.

**Fig. S14.**
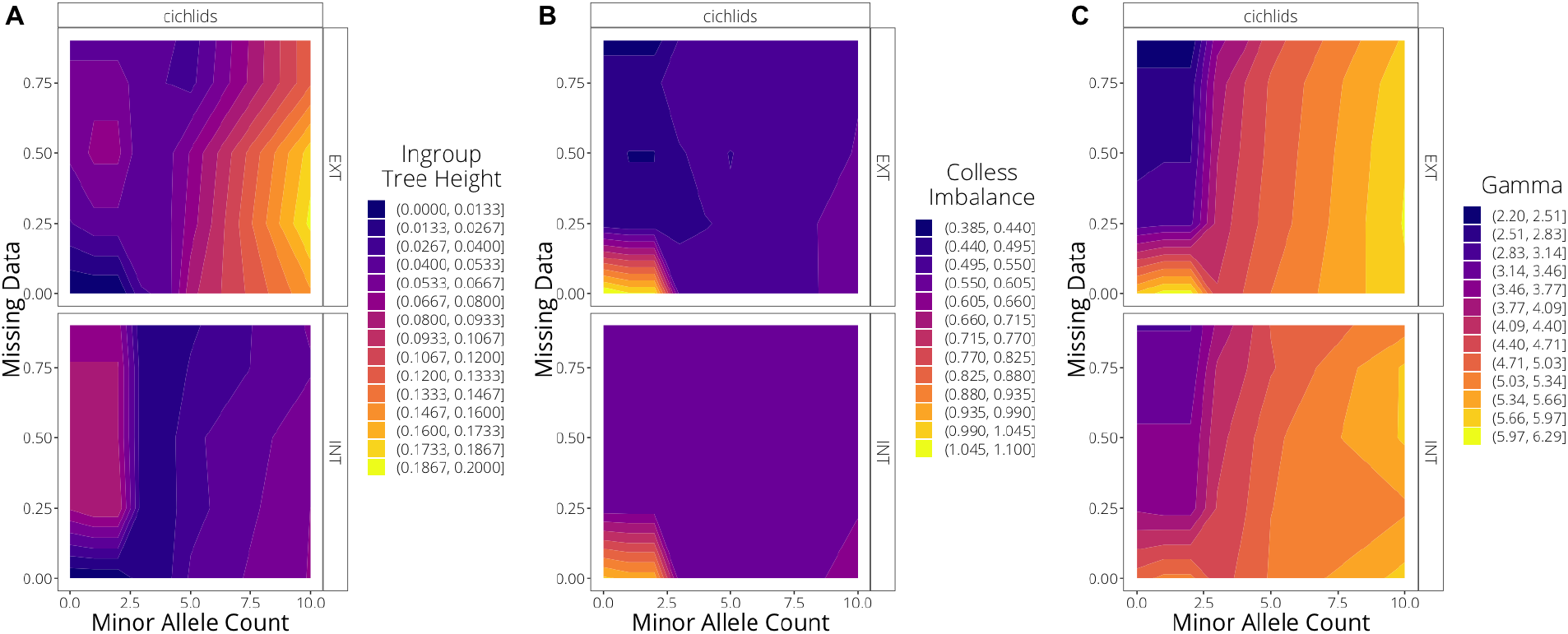
Heatmaps for the **empirical Tropheine trees** indicating how each of our tree descriptive parameters varies in two-dimensional space with minor allele count and missing data thresholds, with darker colors indicating lower values and lighter colors indicating higher values. The gamma and Colless imbalance statistics are measures of tree center of gravity and tree shape, respectively.

**Fig. S15.**
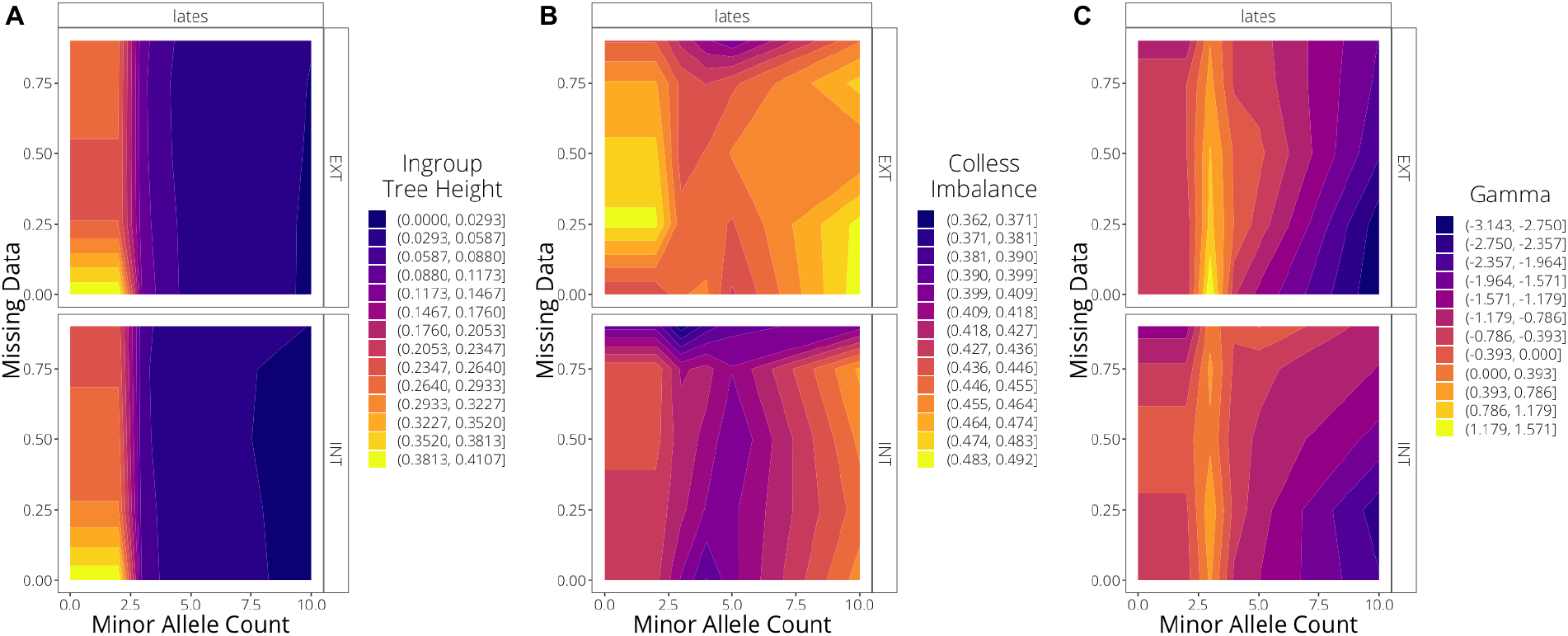
Heatmaps for the **empirical *Lates* trees** indicating how each of our tree descriptive parameters varies with minor allele count and missing data thresholds, with darker colors indicating lower values and lighter colors indicating higher values. The gamma and Colless imbalance statistics are measures of tree center of gravity and tree shape, respectively.

**Fig. S16.**
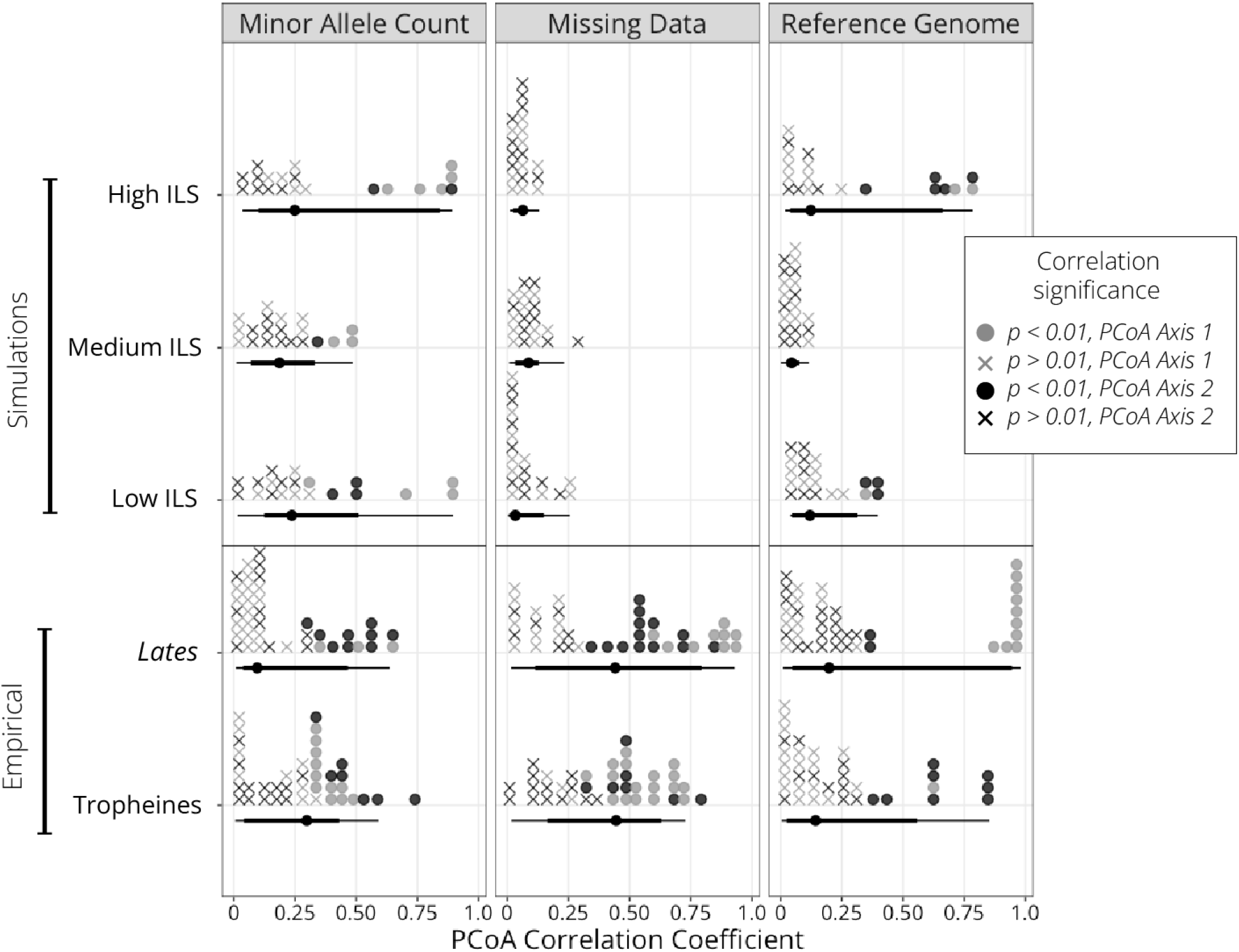
Principal coordinate analyses (PCoAs) based on pairwise Robinson-Foulds distances between `ASTRAL`-inferred trees demonstrate the relationship between simulated and true trees in two dimensional tree space. Plot shows correlation coefficients between each of the three bioinformatic choices and the first two PCoA axes for all simulated and empirical iterations, demonstrating the relative importance of minor allele count in the simulated data and missing data in the empirical data sets, while reference genome was important for all. Dotplots are colored by correlation significance (*p*-value *<*0.01 in solid circles, *p*-value ⩾ 0.01 shown with x’s), assessed using Pearson’s correlation coefficients. Lines below dots show median, 95% quantiles (thick lines), and full data range (thin lines) for each group of analyses.

**Fig. S17.**
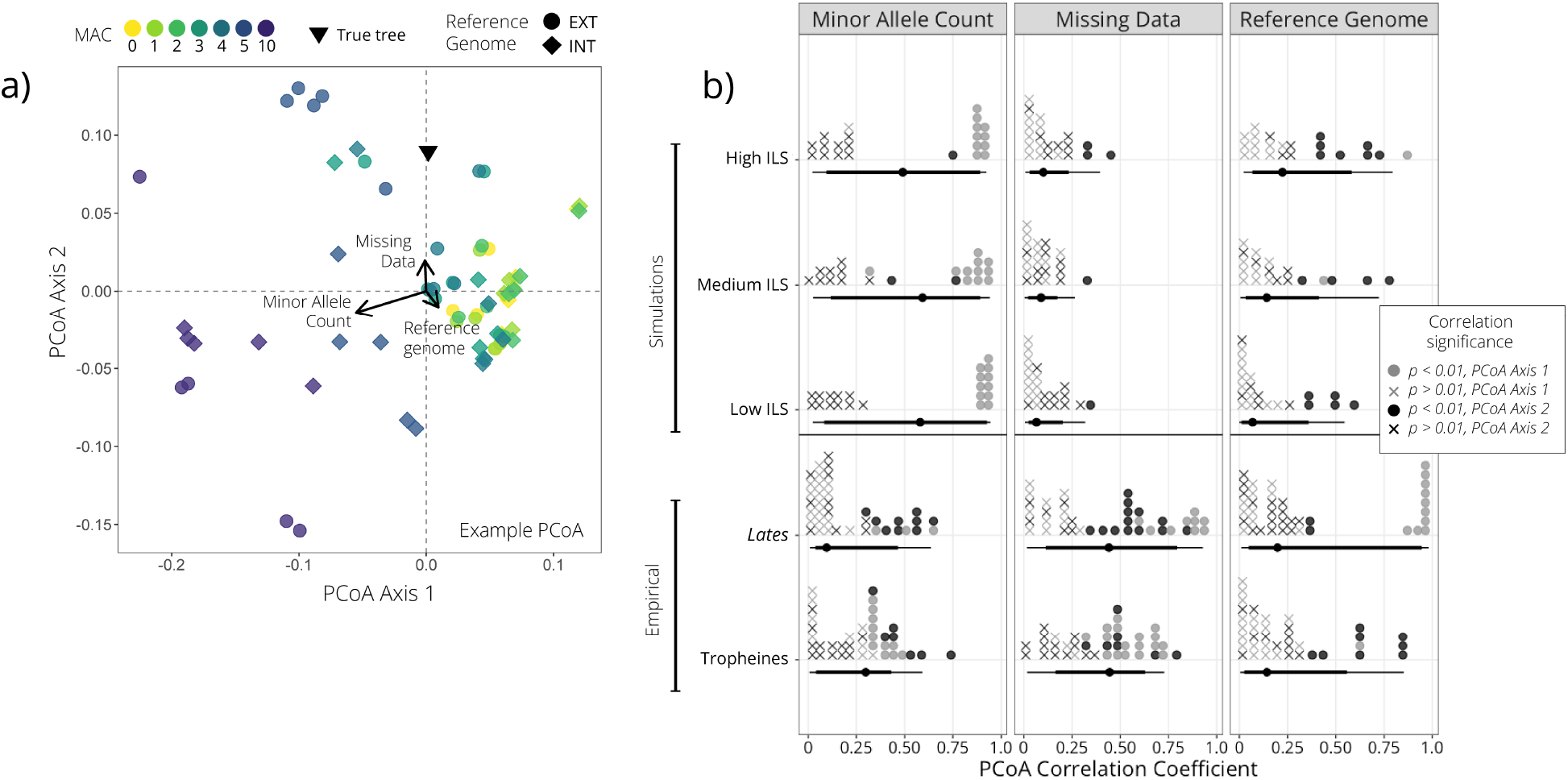
Principal coordinate analyses (PCoAs) based on pairwise Robinson-Foulds distances between `ASTRAL`-inferred trees demonstrate the relationship between simulated and true trees in two dimensional tree space. (a) Example PCoA plot of trees from one iteration of simulated data (high ILS, simulation 18), with the true tree indicated and correlations between PCoA axes and each of reference genome choice, minor allele count (MAC), and missing data threshold visualized via arrows. (b) Summary of correlations between each of the three bioinformatic choices and the first two PCoA axes for all simulated and empirical iterations, demonstrating the relative importance of minor allele count in the simulated data and missing data in the empirical data sets, while reference genome was important for all. Dotplots show the distribution of values, colored by correlation significance (p-value *<*0.01 in solid circles, p-value *>*0.01 shown with x’s), assessed using Pearson’s correlation coefficients. Lines below dots show median, 95% quantiles (thick lines), and full data range (thin lines) for each group of analyses.

**Fig. S18.**
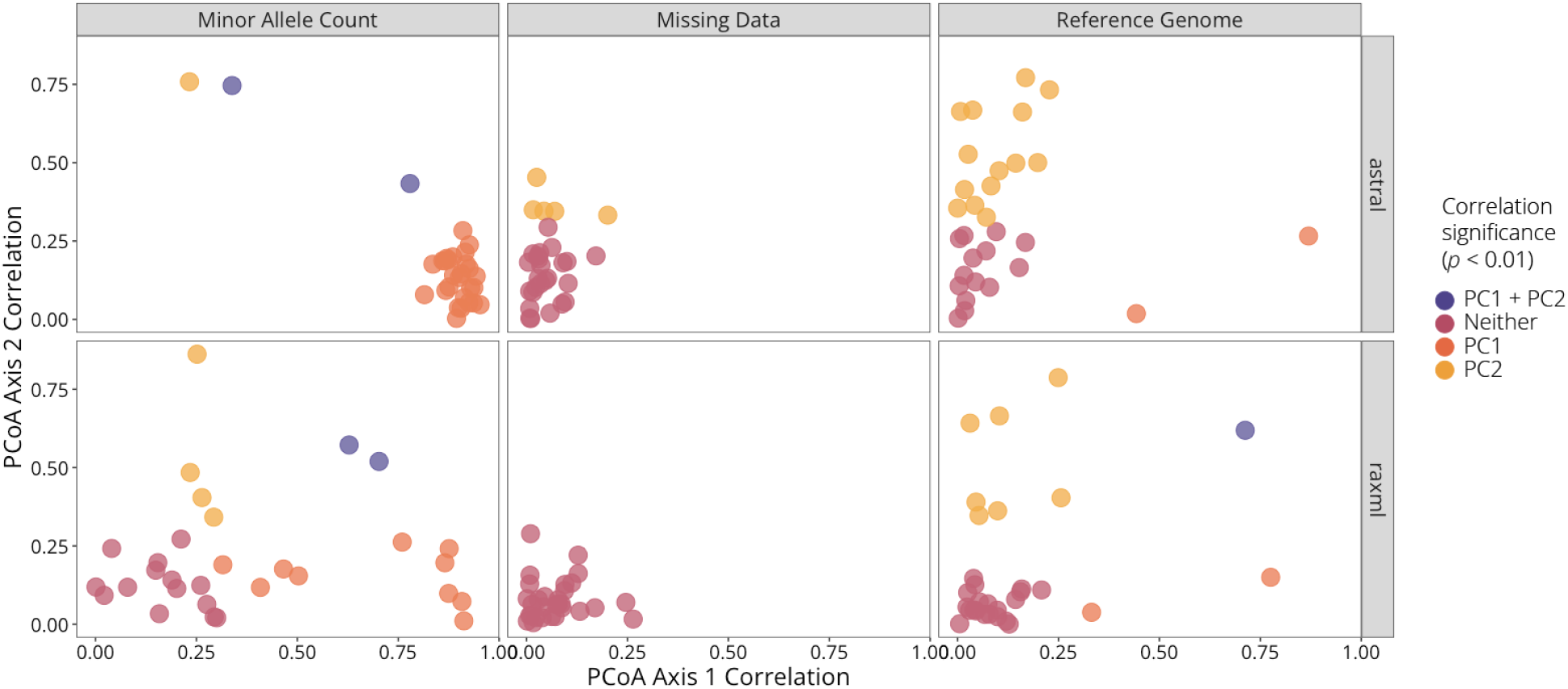
Comparison of PCoA loadings between `RAxML` and `ASTRAL` trees, based on all pairwise Robinson-Foulds distances between trees inferred with the given method. For each parameter, the correlation coefficient between the parameter and a tree’s location in PCoA space for each dataset is plotted, with points colored by significance of the correlation (orange, PC1 *p <* 0.01; yellow, PC2 *p <* 0.01; purple, PC1 & PC2 both *p <* 0.01; pink, both *p >*= 0.01). Using both phylogenetic inference methods, minor allele count and reference genome were generally strongly correlated with the first two principal component axes.

**Fig. S19.**
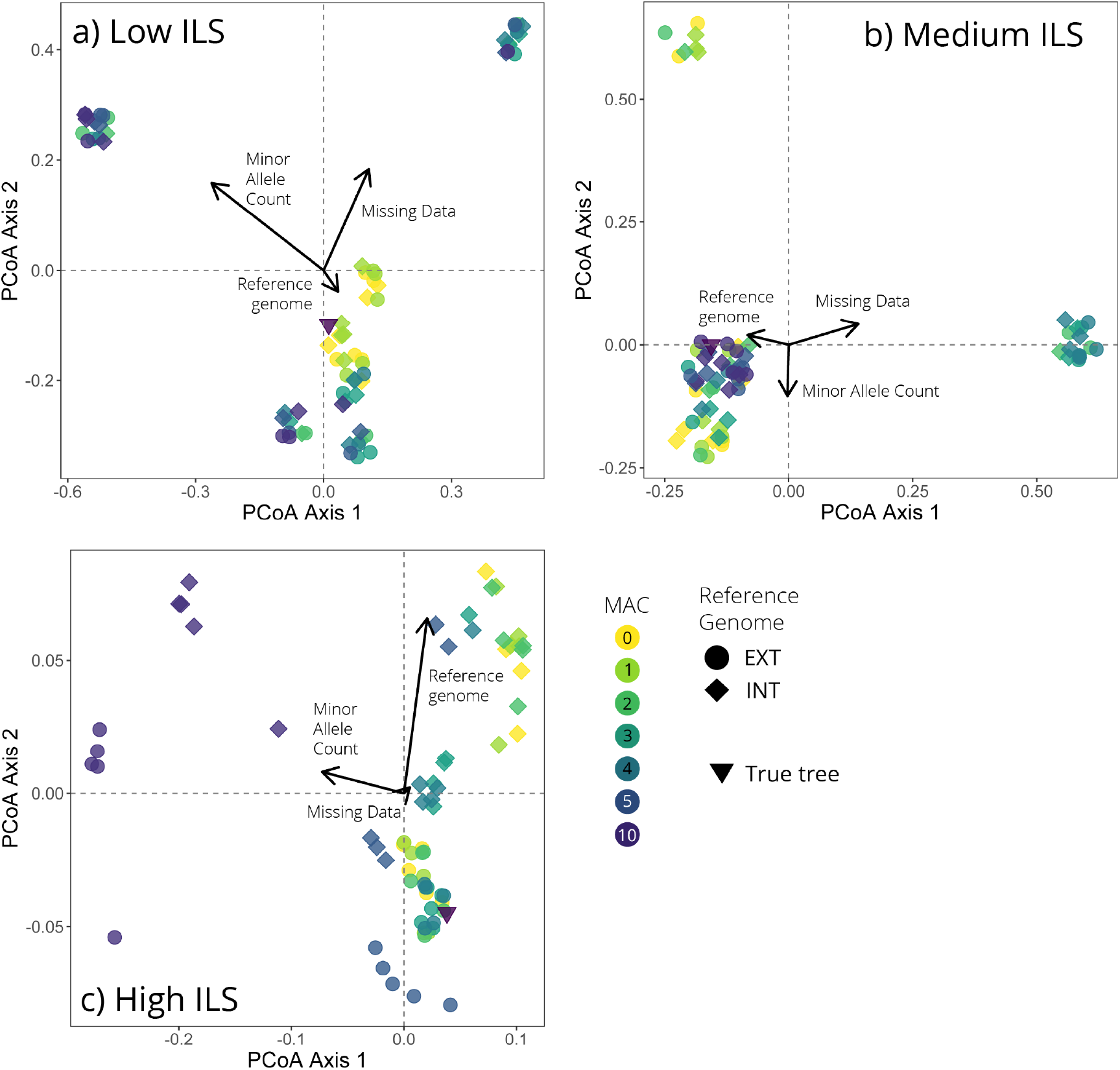
Example plots of principal coordinates analysis (PCoA) visualizing distance between trees in tree space for (A) low ILS, (B), medium ILS, and (C) high ILS trees. For each plot, the true, simulated species tree is shown as a black triangle, while the other points are colored according to their minor allele count threshold and shape indicates the reference genome choice (circle = EXT/outgroup reference; diamond = INT/ingroup reference). Arrows on the plots show correlations of each of the three bioinformatic choices of interest with the first two PCoA axes, with the length of the arrow corresponding to the magnitude of correlation with the given bioinformatic choice. Note that points have been jittered slightly to make it clear when multiple trees overlap in tree space.

**Fig. S20.**
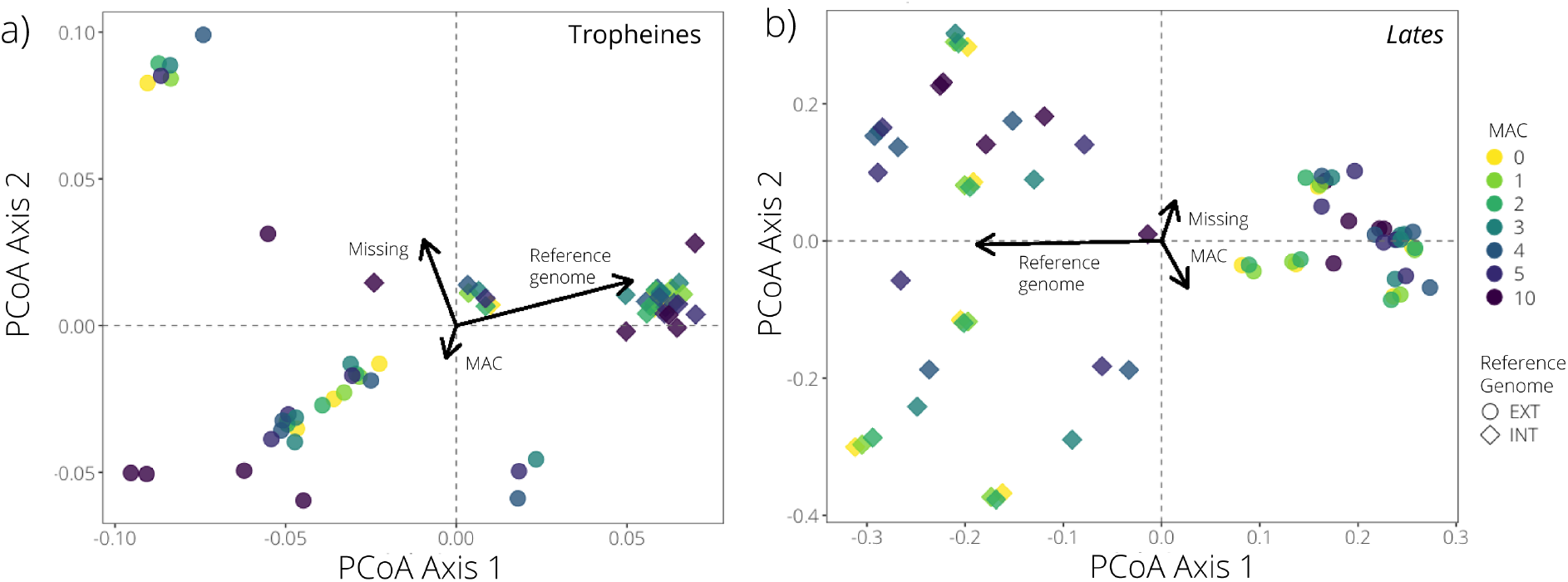
Example plots of principal coordinates analysis (PCoA) visualizing distance between trees in tree space for (a) Tropheine and (b) *Lates* trees. For each plot, the points are colored according to their minor allele count threshold and shape indicates the reference genome choice (circle = EXT/outgroup reference; diamond = INT/ingroup reference). Arrows on the plots show correlations of each of the three bioinformatic choices of interest with the first two PCoA axes, with the length of the arrow corresponding to the magnitude of correlation with the given bioinformatic choice.

**Fig. S21.**
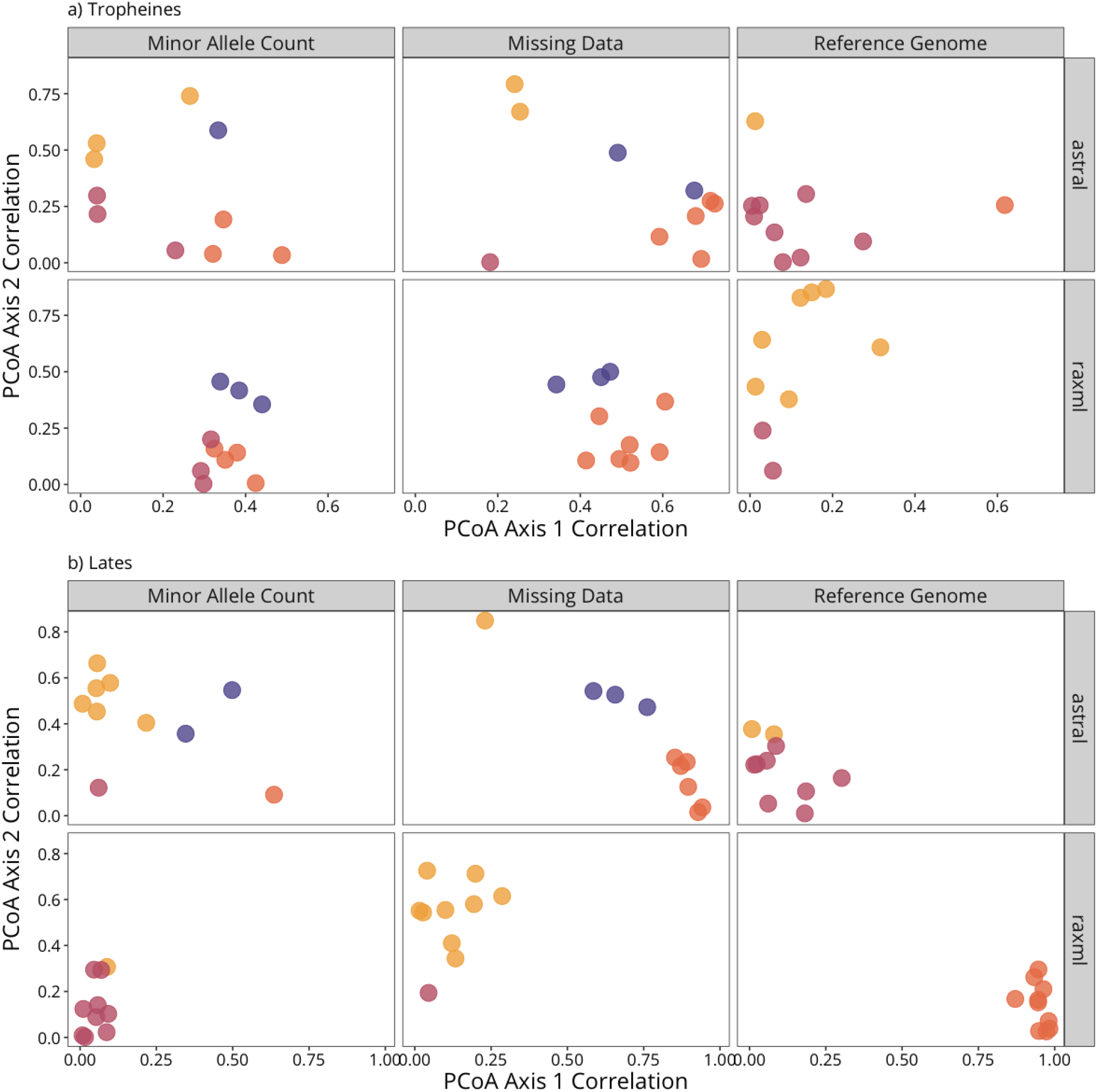
Comparison of PCoA axis correlations between `RAxML` and `ASTRAL` trees for the empirical Tropheine (a) and *Lates* (b) datasets, based on all pairwise Robinson-Foulds distances between trees. For each parameter, the correlation coefficient between the parameter values and each tree’s location in PCoA space for each dataset is plotted (i.e., each point is one of the ten replicate datasets), with points colored by significance of the correlation (orange, PC1 *p <* 0.01; yellow, PC2 *p <* 0.01; purple, PC1 and PC2 both *p <* 0.01; pink, both *p* ⩾ 0.01).

**Fig. S22.**
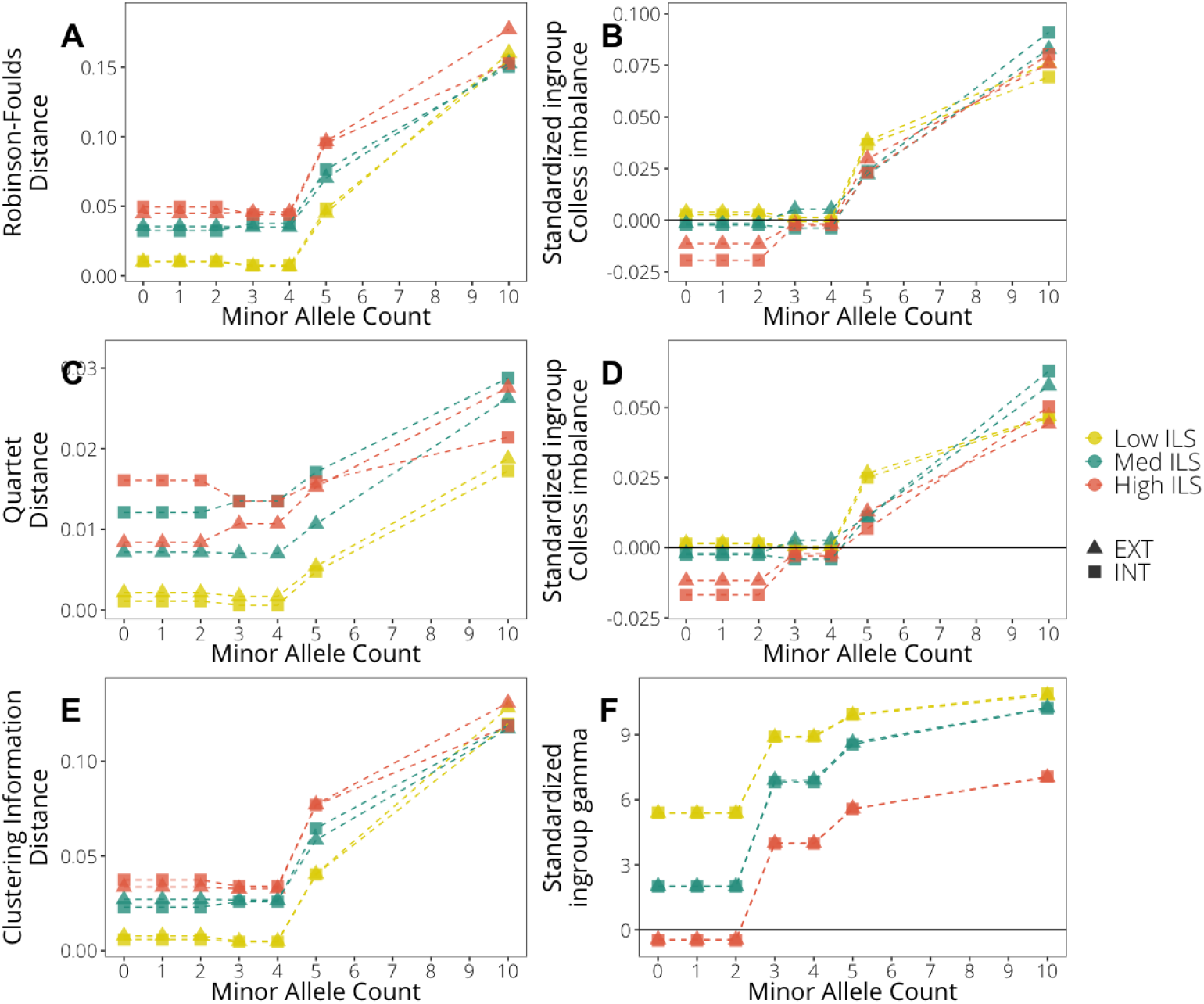
Patterns of variation for the full simulation data sets in the three metrics of distance to the true tree, two metrics of tree imbalance, and tree center of gravity (*γ*) across minor allele counts, with shapes indicating the reference used (EXT, outgroup reference; INT, ingroup reference) and colors indicating the level of ILS. For topological accuracy metrics (Robinson Foulds, quartet, and clustering information distances to the true tree), higher values indicate topologies more biased away from the true tree, and all have been normalized to be bounded by 0 ⩽ distance ⩽ 1. Measures of Colless imbalance, Sackin imbalance, and *γ* have been standardized so that a value of 0 is equal to the measure in the true tree.

**Fig. S23.**
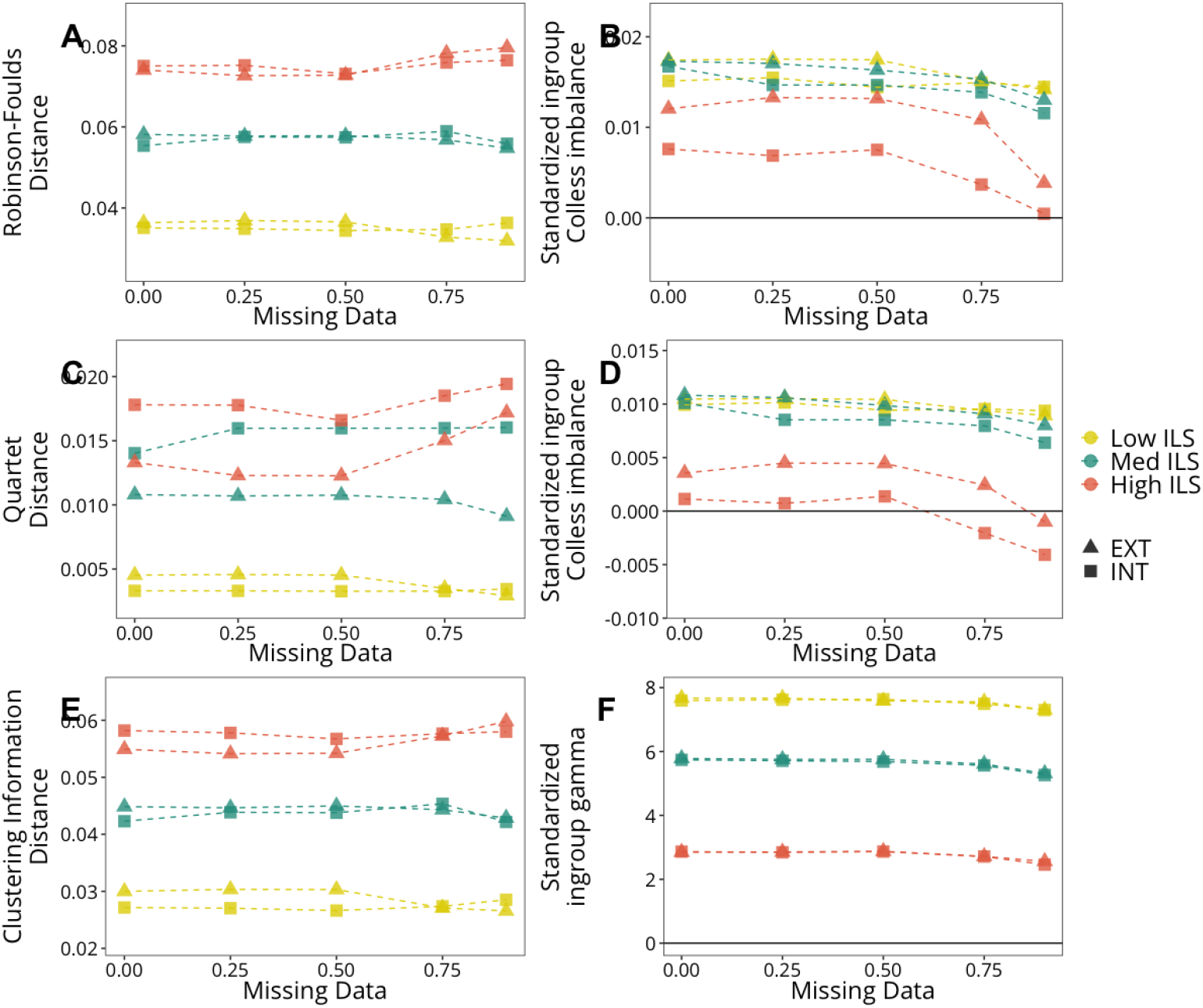
Patterns of variation for the full simulation data sets in the three metrics of distance to the true tree, two metrics of tree imbalance, and tree center of gravity (*γ*) across missing data thresholds, with shapes indicating the reference used (EXT, outgroup reference; INT, ingroup reference) and colors indicating the level of ILS. For topological accuracy metrics (Robinson Foulds, quartet, and clustering information distances to the true tree), higher values indicate topologies more biased away from the true tree, and all have been normalized to be bounded by 0 ⩽ distance ⩽ 1. Measures of Colless imbalance, Sackin imbalance, and *γ* have been standardized so that a value of 0 is equal to the measure in the true tree.

**Fig. S24.**
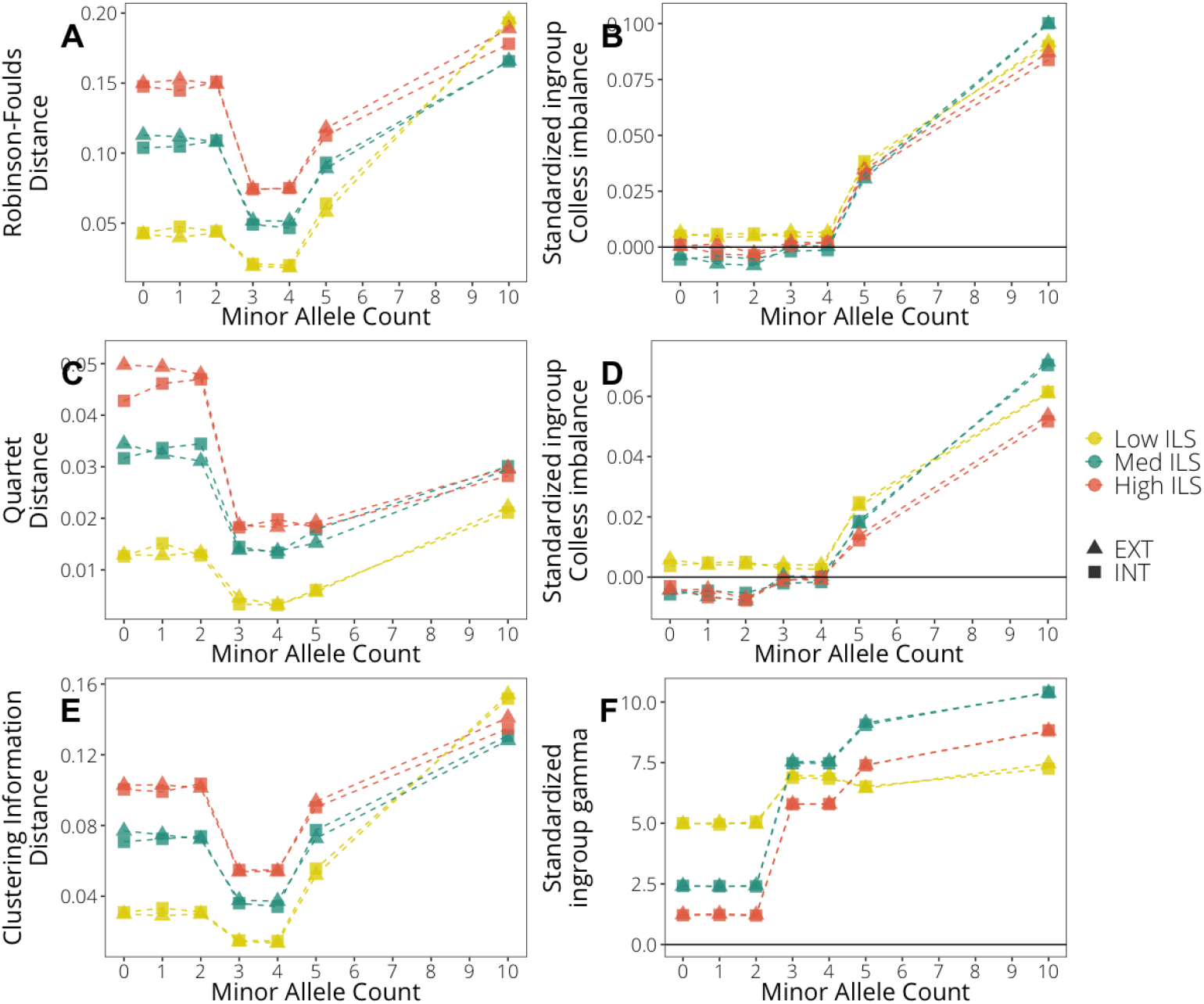
Patterns of variation for the subsampled simulation data sets in the three metrics of distance to the true tree, two metrics of tree imbalance, and tree center of gravity (*γ*) across minor allele counts, with shapes indicating the reference used (EXT, outgroup reference; INT, ingroup reference) and colors indicating the level of ILS. For topological accuracy metrics (Robinson Foulds, quartet, and clustering information distances to the true tree), higher values indicate topologies more biased away from the true tree, and all have been normalized to be bounded by 0 ⩽ distance ⩽ 1. Measures of Colless imbalance, Sackin imbalance, and *γ* have been standardized so that a value of 0 is equal to the measure in the true tree.

**Fig. S25.**
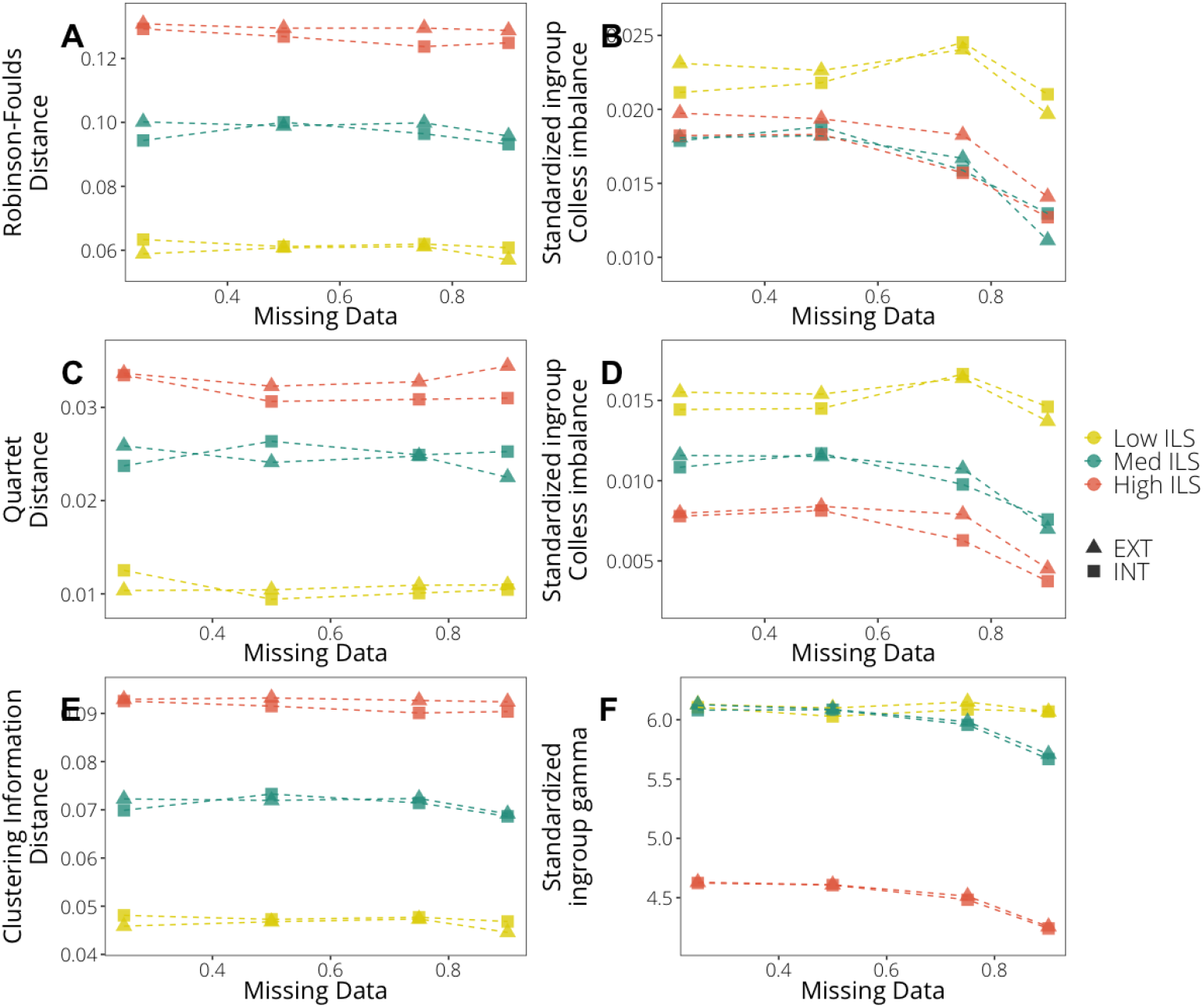
Patterns of variation for the subsampled simulation data sets in the three metrics of distance to the true tree, two metrics of tree imbalance, and tree center of gravity (*γ*) across missing data thresholds, with shapes indicating the reference used (EXT, outgroup reference; INT, ingroup reference) and colors indicating the level of ILS. For topological accuracy metrics (Robinson Foulds, quartet, and clustering information distances to the true tree), higher values indicate topologies more biased away from the true tree, and all have been normalized to be bounded by 0 ⩽ distance ⩽ 1. Measures of Colless imbalance, Sackin imbalance, and *γ* have been standardized so that a value of 0 is equal to the measure in the true tree.

**Fig. S26.**
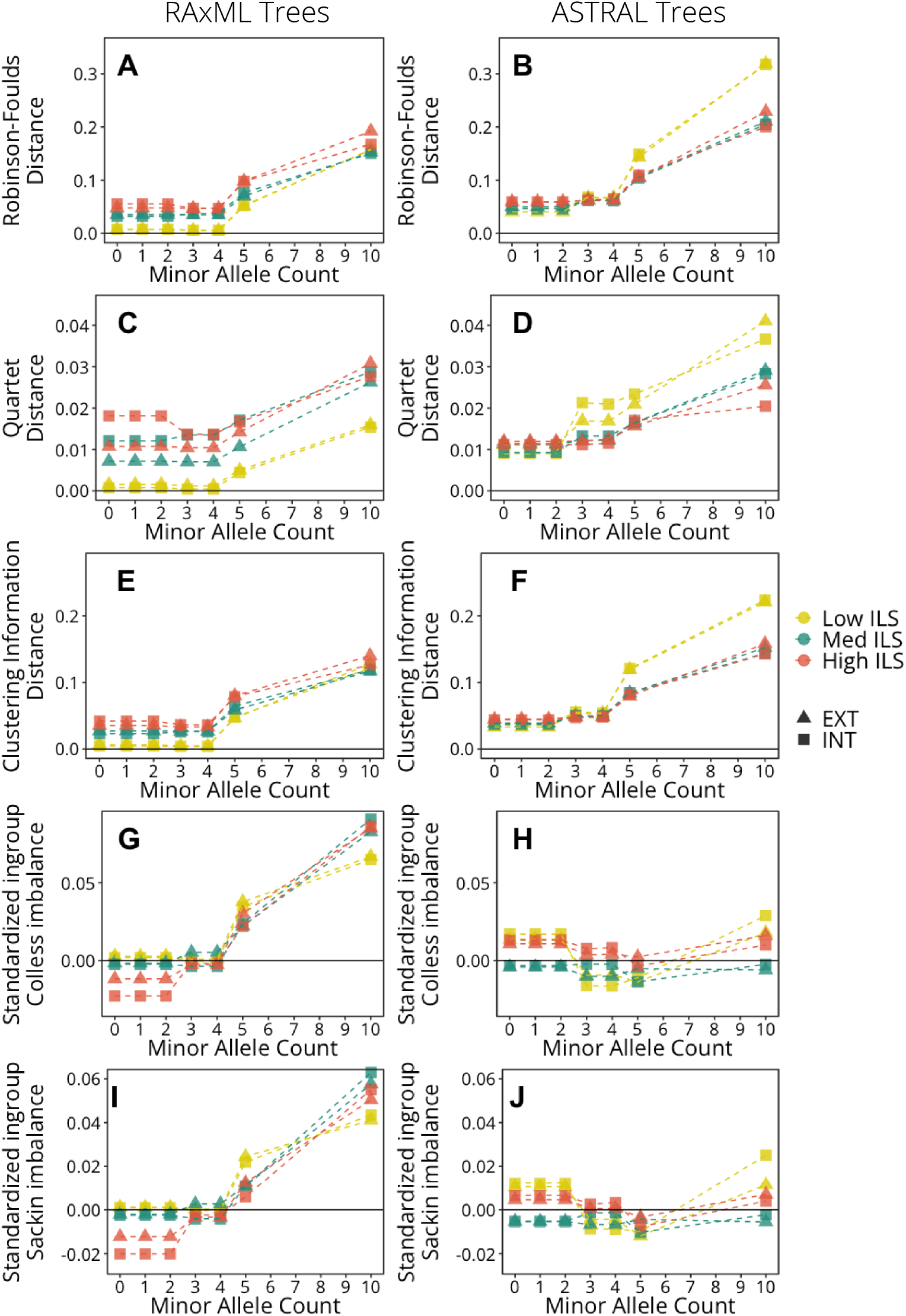
Patterns of variation for the full simulation data sets for `RAxML` (left column) and `ASTRAL` (right column) trees in the three metrics of distance to the true tree and two metrics of tree imbalance across minor allele counts. Shapes indicate the reference that reads were aligned to (EXT, outgroup reference; INT, ingroup reference) and colors indicate the level of ILS in the true tree. For topological accuracy metrics (Robinson Foulds, quartet, and clustering information distances to the true tree), higher values indicate topologies more biased away from the true tree, and all have been normalized to be bounded by 0 ⩽ distance ⩽ 1. Measures of Colless imbalance and Sackin imbalance have been standardized so that a value of 0 is equal to the measure in the true tree. In all plots, black lines indicate the value for the true tree.

**Fig. S27.**
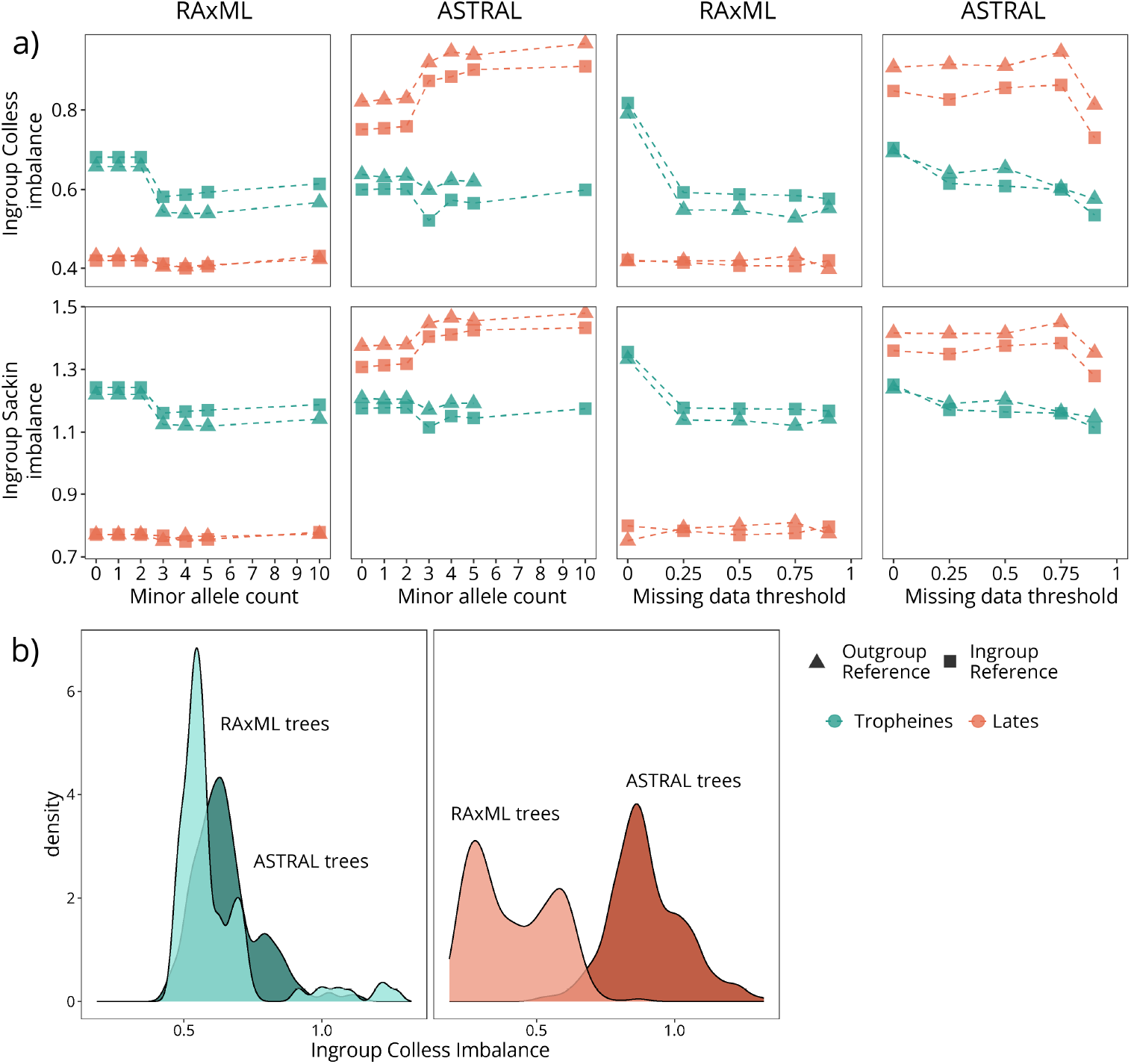
Patterns of variation for the empirical data sets for `RAxML` (left column of each pair) and `ASTRAL` (right column of each pair) trees in the two metrics of tree imbalance across minor allele counts (left) and missing data (right). Shapes indicate the reference that reads were aligned to and colors indicate the data set. In (b), distributions of imbalance statistics are shown for each empirical data set comparing `RAxML` (light colors) to `ASTRAL` (dark colors) trees, demonstrating that trees inferred in `ASTRAL` are generally more imbalanced.

**Fig. S28.**
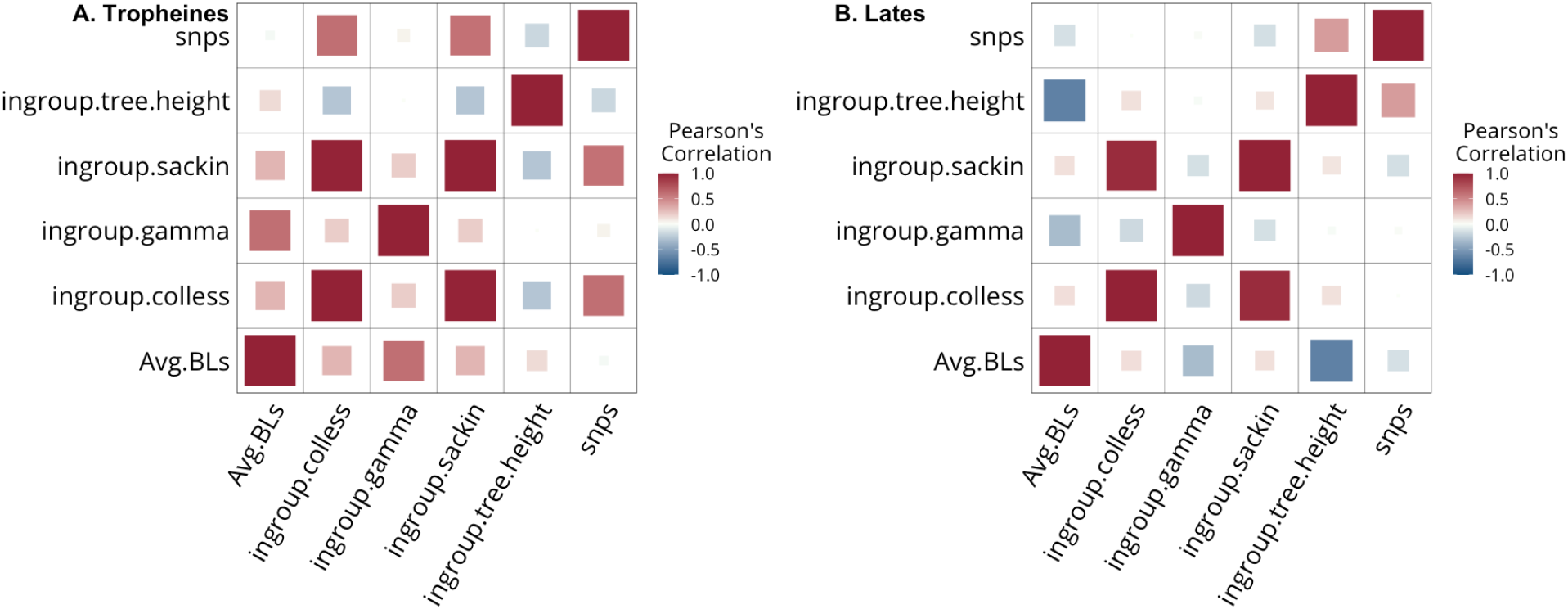
Correlation matrix between output tree characteristics for the (A) Tropheine and (B) *Lates* empirical data sets. Color and size correspond to the magnitude of the correlation. Variables are ingroup tree height (ingroup.tree.height), Sackin imbalance of the ingroup (ingroup.sackin), *γ* of the ingroup (ingroup.gamma), Colless imbalance of the ingroup (ingroup.colless), and average branch lengths (Avg.BLs). White squares indicate pairs with non-significant Pearson’s correlation coefficients at *α* = 0.05.

**Fig. S29.**
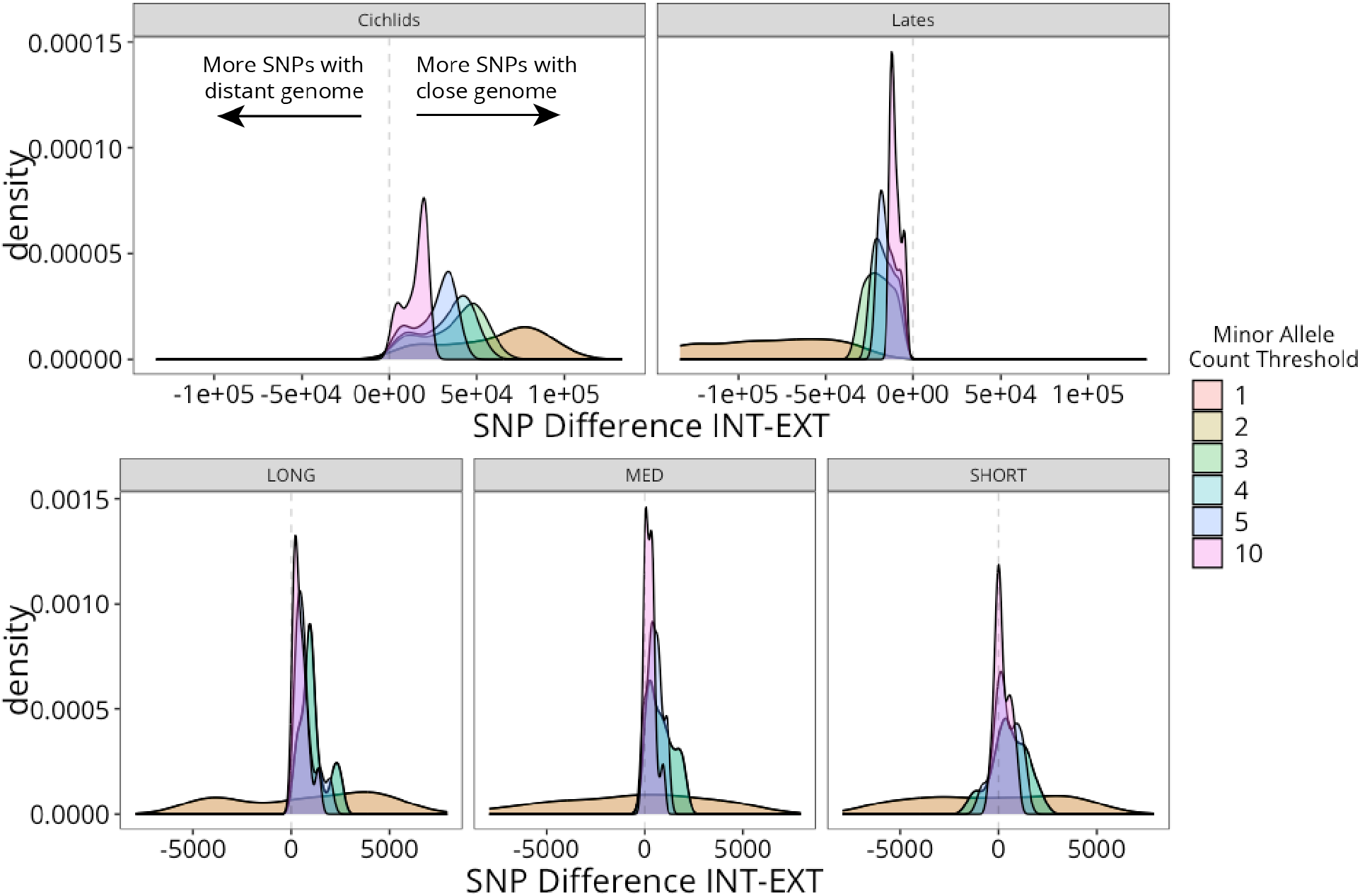
A comparison of data set size (i.e., number of SNPs) between data sets differing only in the reference genome identity. For each plot, density plots are shown for the difference between number of SNPs with the close reference genome (”INT”) and the number of SNPs with the distant reference genome (”EXT”); data sets are colored by the minor allele count threshold used in filtering. For all three ILS levels in the simulations and the tropheine clade, data sets aligned to the close reference genome generally had more SNPs, while the opposite was true for the *Lates* data sets. At low MAC thresholds in the simulations, there was less of a clear trend in which data sets had more SNPs.

**Fig. S30.**
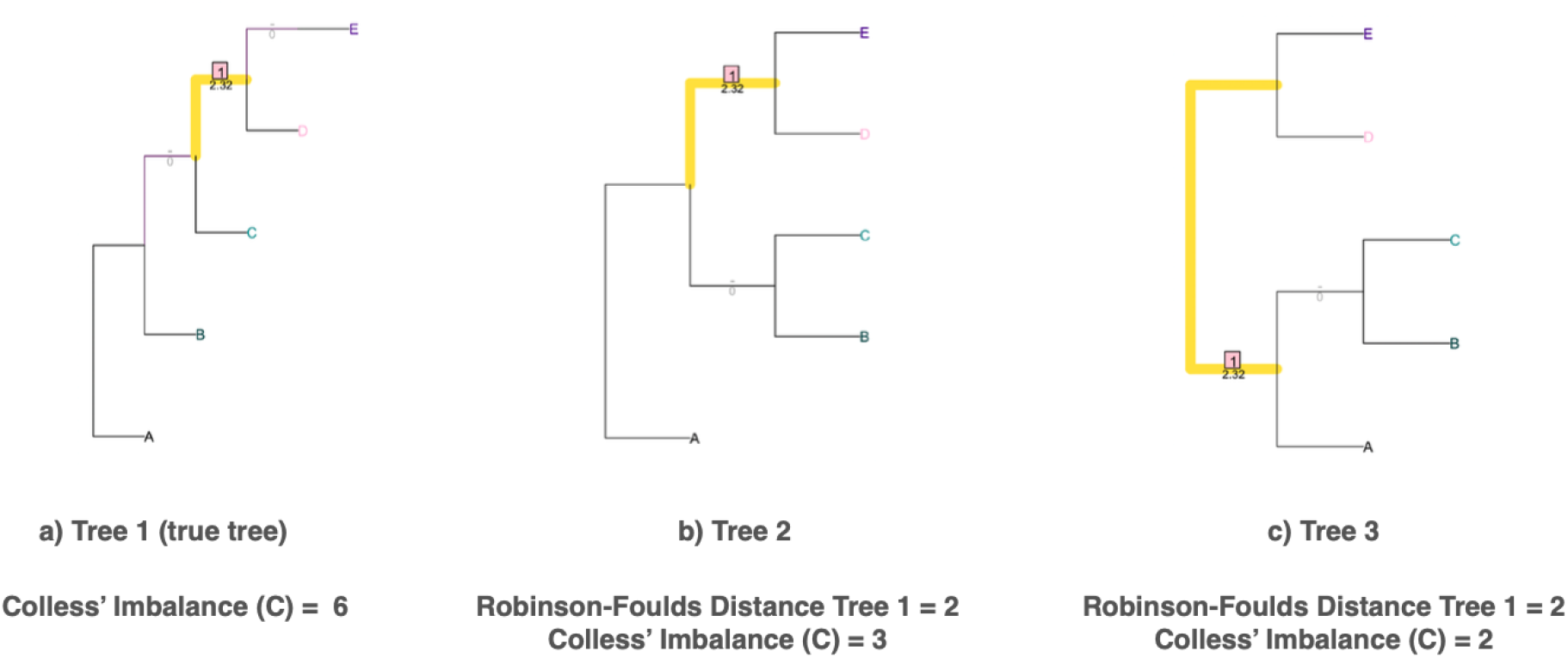
The imbalance of a topology and its distance with respect to the true topology can vary independently. a) The ‘true’ topology used to compare Trees 2 (b) and 3 (c) when calculating Robinson-Foulds (RF) distances. Highlighted (yellow) branches indicate shared splits between the focal tree and Tree 1 (i.e., the split isolating clade DE). For Tree 2, the split CB is unique compared to Tree 1 while the split CDE in Tree 1 is unique compared to Tree 2, resulting in RF distance = 1 + 1 = 2. Similarly, for Tree 3 the split CB is unique compared to Tree 1, which again has the unique split CDE resulting in RF distance = 1 + 1 = 2. Note that since RF distances are computed excluding the root edge the most basal splits are excluded. The pectinate topology of Tree 1 has the highest Colless imbalance score (C) of (4 *−* 1) + (3 *−* 1) + (2 *−* 1) = 6. Tree 2 has a C = (4 *−* 1) = 3 and Tree 3 has a C = (3 *−* 2) + (2 *−* 1) = 2.

**Fig. S31.**
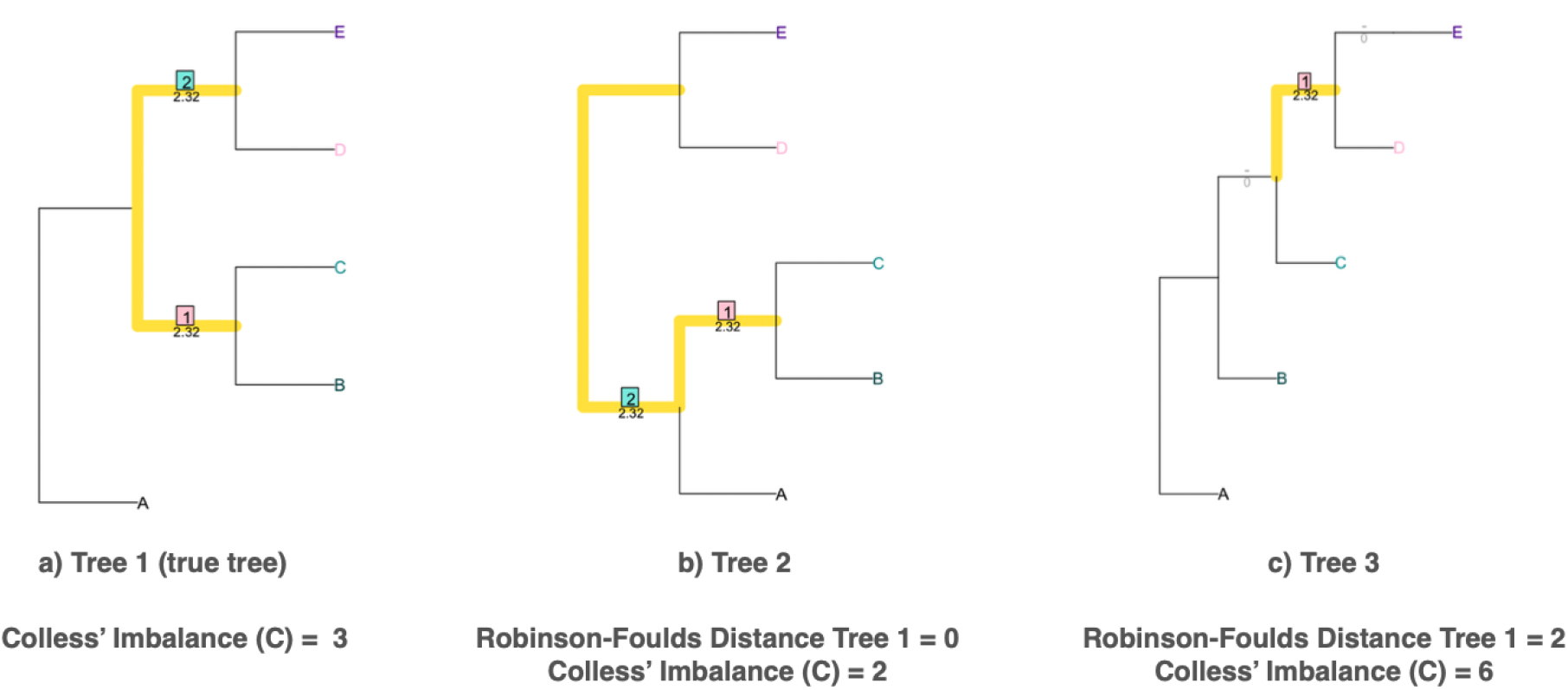
The imbalance of a topology and its distance with respect to the true topology can vary independently. As before (Fig. S30), highlighted (yellow) branches indicate shared splits between the focal tree and Tree 1 (i.e., the split isolating clade DE). For Tree 2, no splits are unique compared to Tree 1 (both CB and DE are shared and ABC in Tree 2 involves the root edge, and is thus ignored), resulting in RF distance = 0 + 0 = 0. For Tree 3 the split CDE is unique compared to Tree 1, which has the unique split CB resulting in RF distance = 1 + 1 = 2. The pectinate topology of Tree 3 has the highest Colless imbalance score (C) of (4 *−* 1) + (3 *−* 1) + (2 *−* 1) = 6. Tree 1 has a C = (4 *−* 1) = 3 and Tree 2 has a C = (3 *−* 2) + (2 *−* 1) = 2.

**Fig. S32.**
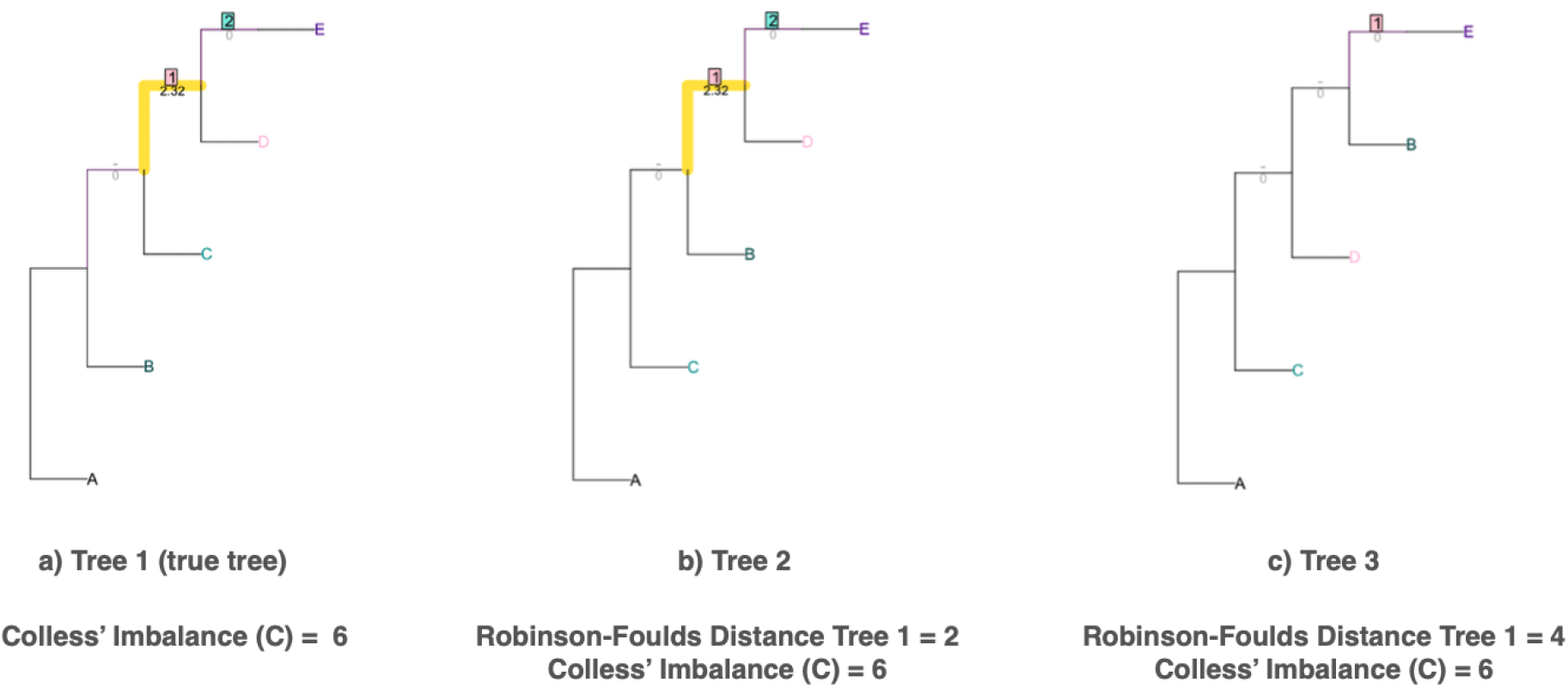
The imbalance of a topology and its distance with respect to the true topology can vary independently. As before (Fig. S30), highlighted (yellow) branches indicate shared splits between the focal tree and Tree 1 (i.e. the split isolating clade DE for Tree 2 and Tree 1). For Tree 2, the split BDE is unique compared to Tree 1 while the split CDE in Tree 1 is unique compared to Tree 2, resulting in RF distance = 1 + 1 = 2. For Tree 3, no splits are shared (minus the root edge split, which again is ignored) with Tree 1 resulting in RF distance = 2 + 2 = 2. All three topologies are pectinate and have the highest Colless imbalance score (C) of (4 *−* 1) + (3 *−* 1) + (2 *−* 1) = 6.

## Notes

http://dx.doi.org/10.5281/zenodo.5940691

### Competing Interest Statement

The authors have declared no competing interest.

### Summary of Updates

Figures 1, 3, 4, 5, and 8 have been improved, and other small edits have been made. Author affiliations have also been updated.

